# SLC3A2 N-glycosylation and alternate evolutionary trajectories for amino acid metabolism

**DOI:** 10.1101/2022.11.15.516651

**Authors:** Cunjie Zhang, Massiullah Shafaq-Zadah, Judy Pawling, Deanna Wan Jie Ng, Geoffrey G. Hesketh, Estelle Dransart, Karina Pacholczyk, Joseph Longo, Anne-Claude Gingras, Linda Z. Penn, Ludger Johannes, James W. Dennis

**Author notes:** Address correspondence and requests for materials to J.W.D.

## Abstract

SLC3A2 (4F2hc, CD98) is an adaptor to the SLC7A exchangers and has undergone extensive repositioning of N-glycosylation sites with vertebrate evolution, presumably in synchrony with the species-specific demands of metabolism. The SLC3A2*SLC7A5 heterodimer imports essential amino acids (AA) and thereby stimulates mTOR signaling, while SLC3A2*SLC7A11 imports cystine required for glutathione synthesis and mitigation of oxidative stress. Analysis of SLC3A2 N-glycans revealed stable site-specific profiles of Golgi remodeling, apart from the conserved N365 site where branching and poly-N-acetylglucosamine content were sensitive to the insertion of lost ancestral sites and to metabolism. N-glycans at N381 and N365 stabilized SLC3A2 in the galectin lattice and opposed endocytosis, while N365 which is nearest the membrane, also promoted down-regulation by galectin-driven clathrin-independent endocytosis (glycolipid-lectin GL-Lect). This is the first report of both positive and negative regulation by galectin binding to N-glycans that are strategically positioned in the same membrane glycoproteins. Proteomics analysis in SLC3A2 mutant HeLa cells with induced re-expression of SLC3A2 as bait revealed the canonical non-N-glycosylated interactors, SLC7A5 and SLC7A11 exchangers, but also AA transporters that were dependent on SLC3A2 N-glycosylation, and are themselves, N-glycosylated AA/Na^+^ symporters (SLC7A1, SLC38A1, SLC38A2, SLC1A4, SLC1A5). The results suggest that the N-glycans on SLC3A2 regulate clustering of SLC7A exchangers with AA/Na^+^ symporters, thereby promoting Gln/Glu export-driven import of essential AA and cystine, with the potential to adversely impact redox balance. The evolution of modern birds (Neoaves) led to improved control of bioenergetics with the loss of genes including SLC3A2, SLC7A-5, -7, -8, -10, BCAT2, KEAP1, as well as duplications of SLC7A9, SLC7A11 and the Golgi branching enzymes MGAT4B and MGAT4C known to enhance affinities for galectins. Analyzing the fate of these and other genes in the down-sized genomes of birds, spanning ∼10,000 species and >100 Myr of evolution, may reveal the mystery of their longevity with prolonged vitality.

**Key Points:**

- Golgi N-glycan remodeling at each site on SLC3A2 differs with the microenvironment.
- The galectin lattice and GL-Lect mediated endocytosis act as opposing forces on trafficking, controlled by N-glycans at the distal N381 and membrane proximal N365 sites, respectively.
- Mutation at N381 or N365 decreased SLC3A2 association with SLC7A5, SLC7A11 and N-glycosylated AA/Na^+^ symporters as well as the capacity to mitigate stress.
- Clustering of SLC3A2*SLC7A exchangers, with AA/Na^+^ symporter and ATPase Na^+^/K^+^ exchanger promotes growth but continuously consumes ATP in non-proliferating cells.
- Bird evolution has improved bioenergetics with the deletion of SLC3A2 and associated transporters; - replaced by transporters of keto acids and a re-enforced galectin lattice.

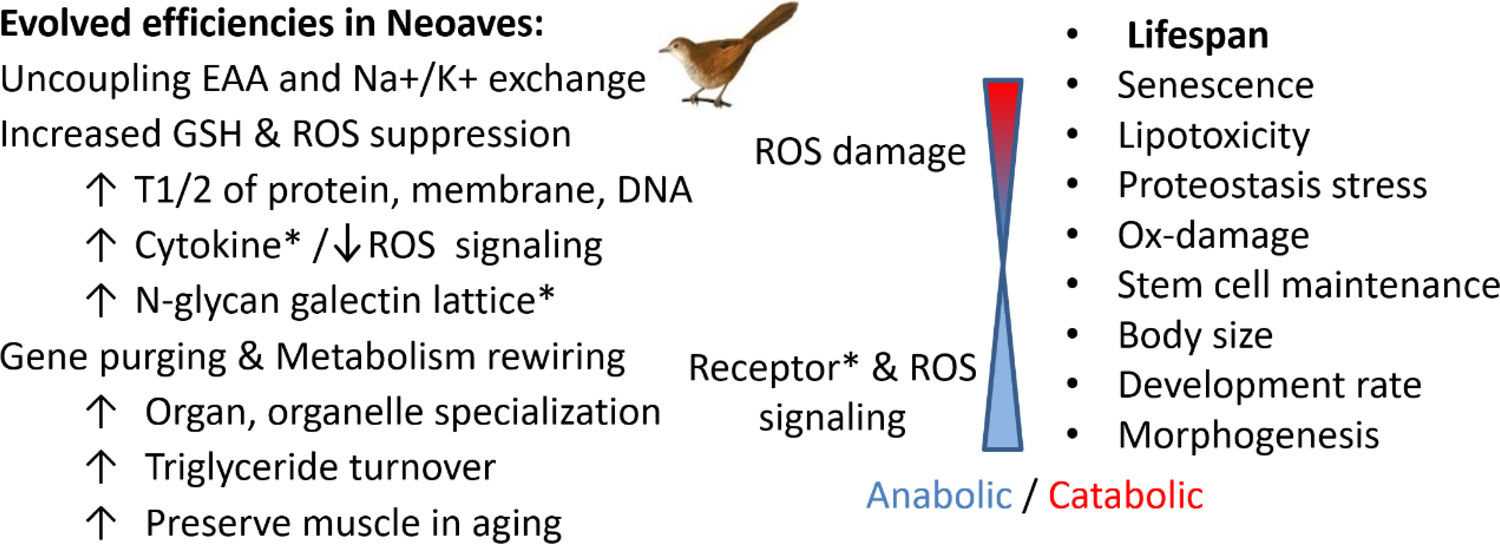

## INTRODUCTION

Evolutionary trajectories are bounded by lineage history and mobility within changing environments. The energetics of mobility requires the cellular uptake of carbohydrates, amino acids (AA) and lipids by membrane transporters that fuel ATP production by oxidative phosphorylation as well as heat and damaging entropy in the form of reactive oxygen species (ROS). Control of these forces requires posttranslational modifications (PTMs) of signaling intermediates in the regulation of gene expression and metabolism (1, 2). This includes N-glycosylation in the endoplasmic reticulum (ER), and N-glycan remodeling in the Golgi. Both steps require uridine diphosphate N-acetylglucosamine (UDP-GlcNAc) generated by the hexosamine biosynthesis pathway (HBP) from glucose, glutamine (Gln) and acetyl-coenzyme A (Ac-CoA), thus competing for these substrates with glycolysis, glutaminolysis and sterol turnover (3–5). The Golgi N-acetylglucosaminyltransferases branching pathway (MGAT1, 2, 4 and 5, and avian-derived MGAT6) requires UDP-GlcNAc to initiate branching of N-glycan on growth receptor kinases (6, 7), integrins (8), glucagon receptor (9), and nutrient transporters in transit to the cell surface (10–12). The N-glycan branching pathway is ordered and requires Mgat5 to generate higher affinity ligands for galectins, a family of lectins that cross-link glycoproteins, thereby inducing phase transition at critical concentrations (termed galectin lattice) that slows the diffusion of receptors and nutrient transporters into clathrin-coated pits and loss to endocytosis (6,9,13). Galectin cross-linking is highly dynamic and intervenes into cell-cell synapses and focal adhesions, transferring entropy that allows exploratory behavior (7,8,14). Galectins also recruit transmembrane glycoproteins into glycolipid-rich invaginations according to the GlycoLipid-Lectin (GL-Lect) hypothesis, which are then internalized within tubular-shaped clathrin-independent carriers, destined for either lysosomal degradation, retrograde trafficking to the Golgi or recycling to the plasma membrane (15, 16).

With support for this physical model of lectin / N-glycan dynamics and the functionality gleaned from cell biology (17) and glycosyltransferase KO mice (18), we are challenged to expand this interaction network at the molecular and physiological levels. Briefly, recent genome-wide CRISPR screens in HAP1 cells revealed correlated essentiality between fatty acid synthase (FASN) and multiple N-glycosylation enzymes, indicating a role for ER and Golgi pathways in lipogenesis (19). Transgenic mice overexpressing either glutamine: fructose-6P-aminotransferase (Gfpt) in the HBP or Golgi Mgat5 in the liver display obesity, steatosis, impaired glucose tolerance and insulin resistance (20, 21). Conversely, Mgat5 deficient mice are resistant to weight gain on a high fat diet (22), similar in this regard to wild type mice on a high fat diet in which branched chain amino acids (BCAA: Leu, Ile, Val) have been reduced (23). Dietary supplementation with GlcNAc markedly increases UDP-GlcNAc and hepatic N-glycan branching, the Gln/essential AA ratio, lipogenesis and weight gain in wild type mice, but much less in Mgat5^-/-^ mice (24). GlcNAc-treated mice are not indolent, but rather utilize equivalent calories more efficiently, as determined by increased conversion to body-mass. This apparent gain in efficiency may include enhanced absorption (uptake) of nutrients and/or efficiencies in oxidative phosphorylation that do not violate the first law of thermodynamics (24). Mgat5 is required, regardless of HBP stimulation, to maintain lean muscle mass and strength with aging. Indeed, Mgat5^-/-^ mice age prematurely with early losses of satellite cells and osteogenic activity, insensitivity to growth factor stimulation (22) and a hepatic AA imbalance (9).

SLC3A2 (4F2hc, CD98) is a type 2 transmembrane glycoprotein that forms disulfide-linked heterodimers (indicated by *) with the large AA transporters (SLC7A-5,6,7,8,10,11), bringing N-glycans to these transporters which are not N-glycosylated (25). SLC3A2*SLC7A5 exchanges essential AA, thereby balancing supply with consumption inside the cell. However, Gln also acts as an export substrate that can drive essential AA import, thereby Leu and Met dependent activation of mTOR and anabolic metabolism (26–28). SLC7A5 expressed with and without SLC3A2 in yeast, which lack the Golgi N-glycan branching pathway, have similar activity indicating that bidirectional transporter does not require SLC3A2 (29, 30). However, SLC3A2 in mammalian cells promotes cell surface residency and transport activity (31, 32), suggesting its adaptor function is dependent on Golgi remodeled N-glycan.

Oxidative stress (ox-stress) induces SLC3A2 and SLC7A11 (xc(-)) expression, thereby Cystine/Glu exchanger activity suppling Cys to glutathione (GSH) biosynthesis (33, 34). GSH mitigates oxidative damage to proteins and lipids thereby opposing cell death by ferroptosis and pyroptosis (35, 36). However, ROS in measured amounts also stimulates growth signaling by reacting with thiol groups which inactivate phosphatases that regulate growth signaling (e.g. PTEN, PTP1B) (37, 38). Conversely, impaired repression of SLC7A11 in cells expressing the Pro47Ser variant of the tumor suppressor p53 leads to increased GSH, which alters the redox state of GAPDH, and in turn, increases RHEB binding to mTOR, thereby stimulating growth (39). In mice, p53 Pro47Ser increased body size, improved metabolic efficiency and fitness, but also increased the incidence of cancer. Thus, SLC3A2*SLC7A11 and SLC3A2*SLC7A5 balance cystine/Glu and BCAA/Gln exchange, respectively, thereby modulating redox state by GSH and mTOR signaling by Leu (26,34,40). Lifespan extension can be achieved in mice by single gene deletions that reduce redundancies in insulin/mTOR signaling (41), or deletions that suppress ROS (e.g. KEAP1, p66SHC) (42–44), but perhaps leaving vulnerabilities (e.g. smaller mass) when challenged in the wild.

SLC3A2 levels increase with cancer progression (45), and conditional deletion of SLC3A2 in the basal epidermis protects against Ras-driven skin carcinogenesis (46), but leads to premature dermal aging with an accelerated loss of extracellular matrix (47). Cancer cells often over-express SLC7A5, SLC7A11, as well as Gln/Glu transport by AA/Na^+^ symporters SLC1A5, SLC38A1 and SLC38A2 (48), perhaps interacting synergistically as explored herein. The efficient import of essential AA and cystine by SLC7A exchangers driven by Gln/Glu promotes mTOR signaling and clonal selection in tumors (49). However, in non-proliferating cells, excessive uptake of BCAA is associated with metabolic stress, insulin resistance and shortened lifespan (23,50–52). BCAA catabolism in mammals contributes to oxidative phosphorylation, lipogenesis, the sterol pathway, glucogenesis, ketogenesis and thermogenesis (53–55); - a complexity in regulation that may have been streamlined to optimize bioenergetics with Neoave evolution.

Herein we asked how the evolution of N-glycosylation sites in SLC3A2 has impacted N-glycan structure, protein interactions and functionality. We deleted the SLC3A2 gene in Flp-In T-REx HeLa cells rescued with a doxycycline (dox)-inducible FLAG-tagged wild type SLC3A2, or SLC3A2 variants with N-glycosylation sites deleted or added at the positions where loss had occurred on the evolutionary path to primates. The four sites on human SLC3A2 displayed distinct N-glycan profiles; those at N365 and N381 were required for Gal3 binding, retention of SLC3A2 at the cell surface, and resistance to ox-stress (H_2_O_2_). The N-glycan at N365 is close to the plasma membrane and promoted Gal3 driven clustering that may facilitate AA exchange between SLC3A2*SLC7A exchangers and AA/Na^+^ symporters. With Neoaves evolution, SLC3A2 and associated transporters were purged, and other genes duplicated, indicating an intriguing shift in AA metabolism supported by additional N-glycan branching and a repositioning of existing adaptor glycoproteins.

## RESULTS

### Evolution of N-glycosylation positions on SLC3A2

The number of N-glycosylation sites in SLC3A2 varies significantly between fish, rodents, and primates (**Fig.1A-C**). The consensus motif is Asn-X-Ser or Thr where X≠ Pro, (NXS/T(X≠ P). Sites in SLC3A2 also vary locally by about 3-6 AAs over mammalian evolution, suggesting a conservative type of “experimentation” with four positions (i.e., 271-273, 355-352, 407-409 and 492-seqiences 498) (56). Sites at all four positions were lost in primates while a site at N381 in a new position was gained. The remaining sites at N365, N424, N506 have 98%, 80% and 97% conservation in mammals, respectively (**Fig. S1A**). The bipartite N-glycosylation motif, <u>N</u>X<u>S/T</u>(X≠ P), displays an encoding asymmetry that accelerates experimentation with N-glycan positions; a feature that enhances the evolvability of glycoproteins (56). The site losses in SLC3A2 were mutations at the Asn, which is more likely due to the encoding (2 Asn and 10 Ser/Thr codons) (57). N381 site gain created an Asn (DSS→ NSS); this is the less probable path, but readily adopted by positive selection in primates. X-ray crystallography and cryo-electron microscopy of human SLC3A2 in complex with SLC7A5 has revealed polar interactions on the extracellular side in close proximity to the membrane (58), where the N-glycan at N365 may contact the membrane, while the primate-derived site at N381 is more distal, in proximity to the galectin lattice (**Fig. 1C**). Intriguingly, selective pressures on SLC3A2 N-glycosylation sites have been greater than other transporters with a similar heterodimeric structure. Notably the Ca^++^ channel (ATP2B1) and Na^+^/K^+^ channel (ATP1A1) (59) have highly conserved N-glycan positions in their adaptors; neuroplastin (NPTN) and ATPase subunit β1(ATP1B1), respectively.

**Figure 1.**
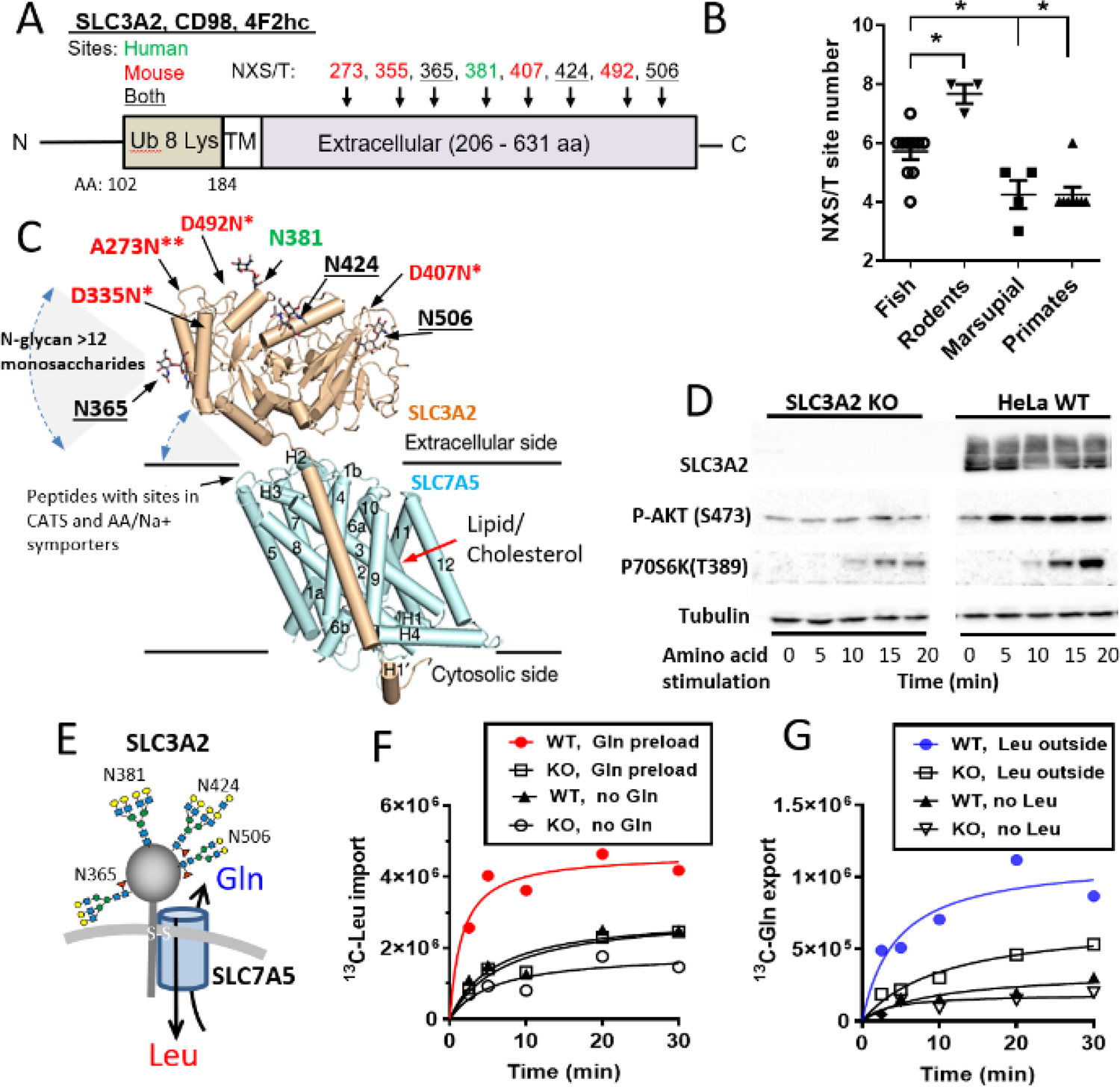
SLC3A2 N-glycosylation sites. **(A)** N-glycosylation sites NXS/T(X≠ P) in the human SLC3A2 sequence (black, green) and losses (red) since a common ancestor with mouse. Eight Lys near the N-terminus are targets for ubiquitination. **(B)** N-glycosylation site number in fish (n=10 species), rodents (n=4), marsupials (n=4) and primates (N=8) * paired t test p <0.02 (see Fig. S1A). **(C)** X-ray crystal structure of SLC3A2 in complex with SLC7A5 AA exchanger (from Yan R. et al. *Nature* 568:127, 2019), and N-glycosylation sites with the linkage monosaccharide attached. The full-length N-glycans are commonly >12 monosaccharide and extend into solution with rotational flexibility about the glycosidic linkages. * and ** indicate one or two mutations away from Asn for site lost since common ancestor with mouse. **(D)** Stimulation of mTOR kinase activity revealed by phosphorylation of p70S6 kinase in WT and SLC3A2 KO HeLa cells starved of amino acids for 1h, followed by 15 min stimulation with Leu, Gln, Arg. **(E)** Schematic representing the major de-sialylated N-glycan structures at each site in SLC3A2 as documented herein (see Fig. 4). SLC3A2 is required for Leu/Gln exchange activity. **(F)** Time course of Leu transport in HeLa cells preloaded ± Gln and pulsed with [U^13^C]-Leu in the medium. **(G)** Cells preloaded with [U^13^C]-Gln and stimulated ± Leu in the medium and measuring the released of [U^13^C]-Gln to the medium.

### Reduced essential AA transport and mTOR activity in SLC3A2 deficient HeLa (KO) cells

The Flp-In T-REx HeLa cells (designated as WT below) were targeted for SLC3A2 deletion by CRISPR-Cas9. Mutant cells displayed reduced capacity for mTOR activation by Leu (**Fig. 1D**), slower rates of growth and cell migration (**Fig. S2A-C**); consistent with phenotypes previously reported for SLC3A2 KO cells (26, 60). ^13^C-Leu uptake and ^13^C-Gln export were impaired by ∼ 3 fold under maximal inside-to-outside gradient conditions, indicating reduced SLC7A5 exchanger activity (**Fig. 1E-G**). In Gln-free medium, D-phenylalanine (D-Phe), an inhibitor of SLC7A5, reduced intracellular Leu, Ile, Val, His by ∼70% in WT cells to levels observed in untreated SLC3A2 KO cells (**Fig. S2D**). D-Phe also reduced these AAs in SLC3A2 KO cells an additional ∼50%, consistent with residual SLC7A5 transporter activity in the absence of SLC3A2 (31). Restoring Gln with or without Torin did not affect BCAA levels in WT or KO cells. However, restoring Gln to the medium increased Glu and reduced Asp levels in WT cells which was reversed by D-Phe. In SLC3A2 KO cells, restoring Gln resulted in only a smaller increase in Glu, and D-Phe had no effect. (**Fig. S2D**). This suggests that SLC3A2 enhances Glu levels in an essential AA-dependent manner, perhaps by spatially coupling SLC3A2*SLC7A5 with AA/Na^+^ symporters (SLC1A and SLC38A families).

To explore these dynamics, cells were conditioned in AA-free medium for 3h, followed by a change of the AA-free medium plus either Bovine Serum Albumin (BSA) or BSA+ Gln, and extracellular AAs were measured at times thereafter (**Fig. S3A,B**). Phagocytosis of BSA and proteolysis in the lysosomes supplies intracellular AAs, and ensuing imbalances are reflected in AA released to the medium (28,61,62). During the 3h AA-fast, WT and mutant cells released 15 of the 19 AAs measured, while Cys, His, Arg and Met were absent in SLC3A2 KO cell medium, and Lys was reduced (**Fig. S3D**). Consistent with cellular control over release, AA levels in medium after 21h did not correlate with the AA composition of BSA (**Fig. 3E**). With the addition of BSA, all AAs were present in the medium by 21h except for Met. Gln levels declined rapidly to a new equilibrium by 2 hours in WT and SLC3A2 KO cell medium. In contrast, Glu concentrations fluctuated in the medium of WT cells indicating a reversibility of transport direction by the cell population, presumably driven by competition between metabolic pathways (**Fig. S3A,B**). In SLC3A2 KO cell cultures, Glu levels were low, and fluctuations suppressed, consistent with reduced cooperativity between transporters and metabolism.

Importantly, Leu, Ile and Val fluctuated in synchrony with Glu levels in the WT cell medium. Trp, Phe, His, Arg and Lys increased continuously, reaching higher levels in SLC3A2 KO than WT medium by 21 and 24 hours. The initial fluctuations (0-3 hours) in Glu and BCAA were suppressed as Trp, Phe and His, which are known to act as counter-ions for essential AA import by SLC7A5 (29), increased (**Fig. S3B**). Rates of metabolism in WT and KO cells were similar as indicated by consumption of hypoxanthine and pyruvate, which were provided in the AA-free DMEM medium (**Fig. S3C**). In BSA+Gln, the release of BCAA as well as Trp, Phe, His, Arg and Lys from WT and KO cells was largely suppressed (**Fig. S3B**). Results in BSA alone are consistent with spatial proximity of SLC3A2*SLC7A5 and AA/Na^+^ symporters that couple BCAA uptake with Glu export in WT cells but are lacking in KO cells. The efficiency of AA capture by transporters decreases exponentially with distance as described by Fick’s laws of diffusion; a limitation that may be circumvented by Gal3-mediated clustering (**Fig. S3A**). The proximity of SLC3A2*SLC7A5 and AA/Na^+^ symporters is expected to increase AA exchange, and thereby ATPase dependent AA/Na^+^ activity. Indeed, WT cells are more sensitive than KO cells to NaCl, which is partially rescued by KCl (**Fig. S3F**).

### Central metabolism is modified in SLC3A2 KO cell

With only a marginally slower growth rate, SLC3A2 KO cells have adapted to a lower capacity for essential AA import. Metabolite profiling by liquid chromatography mass spectrometry (LC-MS/MS) revealed 52%-18% lower levels of His, Val, Ile, Trp, Met, Leu, Tyr Phe; as well as lower Cys and GSH, consistent with reduced SLC7A5 and SLC7A11 activities, respectively, and similar to previous reports on SLC3A2 KO mouse ES cells (63) and human tumor cells (31, 64). Increased Gln, Asp, Arg, Lys, Asn and Ser levels are apparently insufficient as efflux substrates to maintain intracellular essential AA levels by SLC7A5 exchange in the absence of SLC3A2 (**Fig. 2A, S2D, Table S1**).

**Figure 2.**
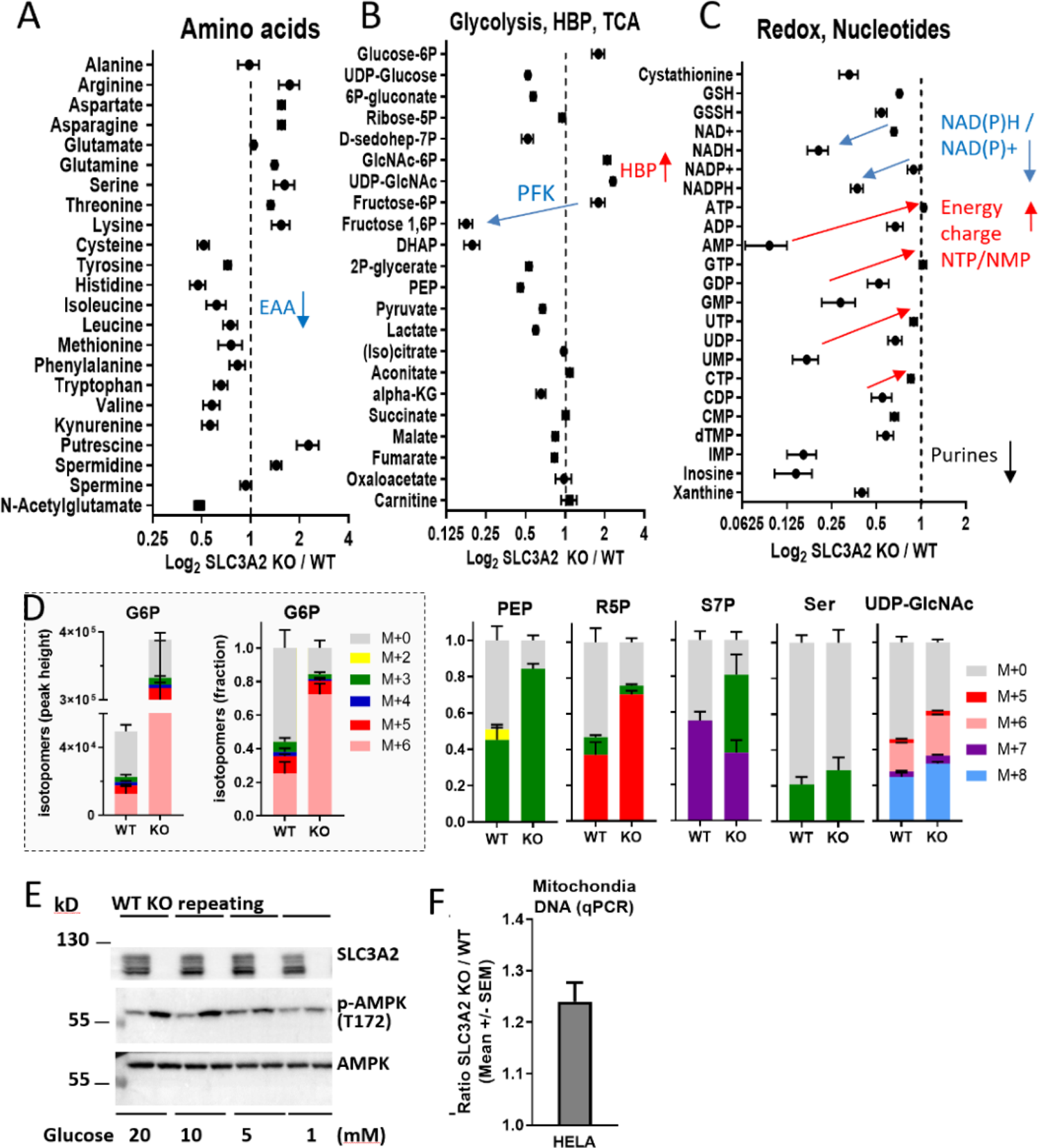
SLC3A2 WT and KO metabolite profiles. **(A-C)** Intracellular metabolites by LC-MS/MS are displayed as KO/WT ratios, mean ± SD biological replicates (n= 6). Phosphofructokinase (PFK) **(D)** Cells were cultured in [U-^13^C]-glucose for 24 h to label downstream metabolites. G6P in the box is displayed as mass intensity/cell (left) and fractional distribution of ^13^C-label (right). Phosphoenolpyruvate (PEP), ribose 5-phosphate (R5P), sedoheptulose 7-phosphate (S7P), Serine (Ser), UDP-GlcNAc. **(E)** AMPK activation in SLC3A2 KO and WT cells cultured in DMEM 1 mM Gln +10% dialyzed FCS with 1 to 20 mM glucose for 48 h. **(F)** relative mitochondrial content by qPCR.

Fructose-6P (F6P) was 4-fold higher and fructose 1,6-bisphosphate (F1,6BP) ∼8 times lower in SLC3A2 KO cells, indicating a decrease in phosphofructokinase (PFK) activity, thereby increasing glucose-6P flux into the oxidative pentose phosphate pathway (ox-PPP), and F6P into the HBP pathway (**Fig. 2B,C**). Indeed, [U-^13^C]-glucose flux to ribose-5P and sedoheptulose-7P was increased, the latter consistent with ribulose-5P conversion to F6P and glyceraldehyde 3-phosphate by the non-oxidative PPP (**Fig. 2D**). Thus, the oxidation of glucose through the PPP has increased while depleting F1,6BP and phosphoglycerate flux to the one-carbon folate cycle, which generates tetrahydrofolates 5,10-meTHF and 10-formylTHF (65). IMP, AMP, GMP levels were severely depleted in SLC3A2 KO cells, suggesting a deficiency in formate production and/or its transfer from mitochondrial to cytosolic 10-formylTHF and into the purine pathway. 10-formylTHF is a substrate in the reduction of NAD(P)^+^ to NAD(P)H, and also provides one-carbon transfer in purine and Met recycling from S-adenylhomocysteine (66). IMP and AMP levels were low while ATP levels were maintained, suggesting enhanced coupling of oxidative phosphorylation, perhaps driving the NADH/NAD^+^ ratio lower (**Fig. 2C**). Despite the bottleneck at PFK, [U-^13^C]-glutamine flux into the TCA cycle was slightly increased, perhaps contributing to elevated Asp levels and the Asp/malate shuttle (**Fig. S4A**) (67). Furthermore, SLC3A2 KO HeLa cells were more sensitive to the glutaminase inhibitor CB839 (**Fig. S4B**). The SLC3A2 KO phenotype extends beyond AAs to central metabolism and pathways that are revisited below in our discussion of the natural SLC3A2 deficiency in birds.

Low levels of F1,6BP provides a means of activating AMPK independent of the AMP/ATP ratio (68). AMPK activation protects against ox-stress in part by increasing glucose flux, mitochondrial biogenesis and autophagy (69). With increasing glucose availability from 1 to 20 mM, AMPK activation increased in SLC3A2 KO cells, while remaining stable in WT cells, consistent with a defect in negative feedback control in high glucose DMEM medium, and increased mitochondria content in KO cells (**Fig. 2E,F**). However, reduced levels of cystathionine, Cys, Met and GSH predicts an increased vulnerability of the KO cells to ox-stress.

### SLC3A2 KO cells are hypersensitive to oxidative-stress

Mutant cells in normal culture conditions displayed a significant increase in the ER stress-response transcripts CHOP, GADD34, XBP1-U and XBP-S (**Fig. 3A**). Cells transferred from normal high glucose/Gln (25mM / 4mM) to lower (10 mM / 1mM) medium stimulated the expression of these stress-response genes in WT cells, but considerably less in SLC3A2 KO cells, suggesting a defect in stress sensing. SLC3A2 KO cells were more sensitive to H_2_O_2_, which was intensified in low nutrient medium or with removal of serum, indicating catabolism and growth factor signaling were required to oppose ox-stress (**Fig. 3B**). High nutrient conditions provided some protection to WT cells in the absence of serum, but less for SLC3A2 KO cells. CHOP, GADD34, XBP1 along with ATF4 and ATF6 regulate expression of antioxidant genes, including SLC3A2, SLC7A11, SLC7A1 and SLC1A5 in nutrient depleted conditions (70, 71).

**Figure 3.**
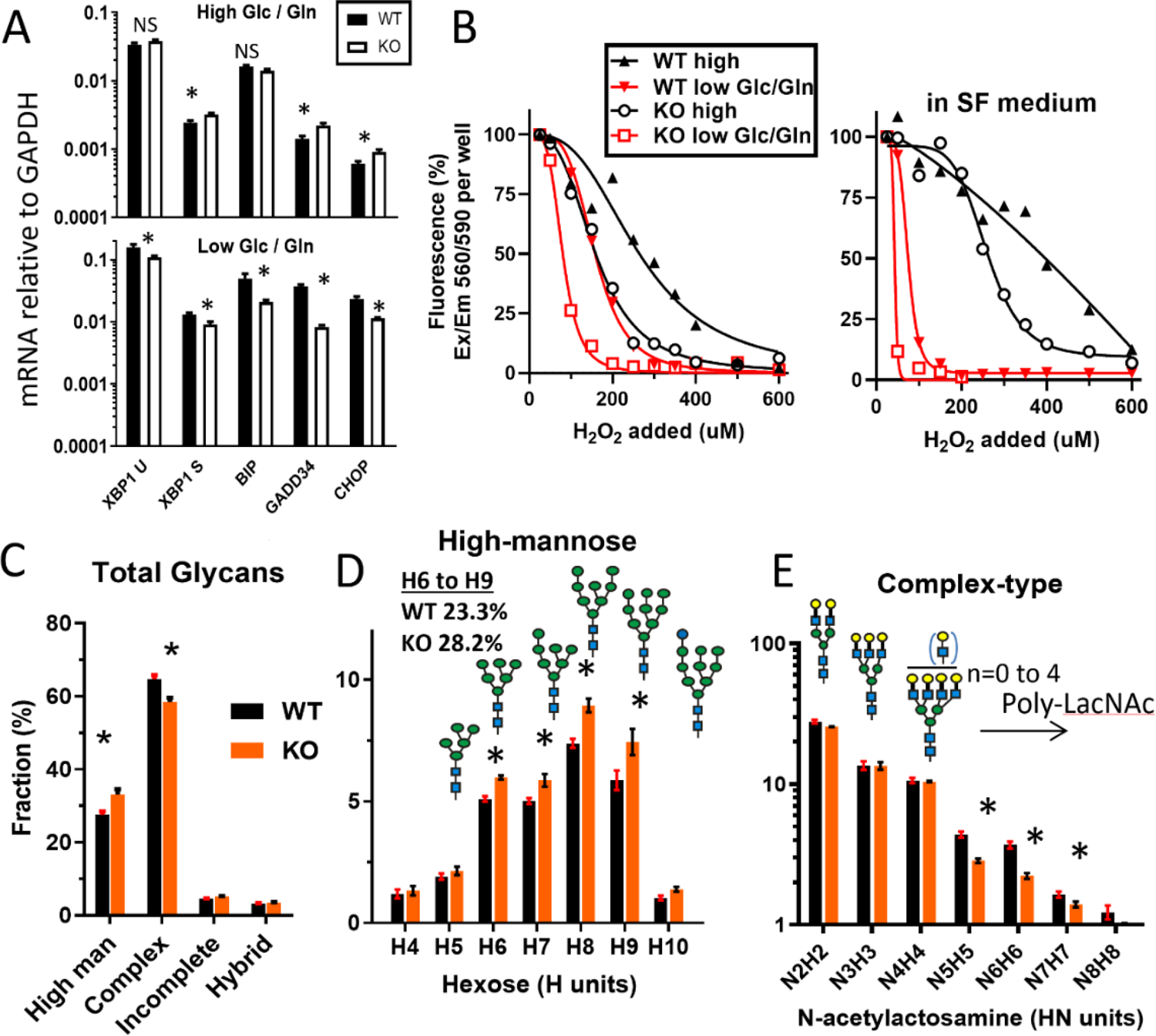
Hypersensitivity of SLC3A2 KO cells to Ox-stress is nutrient dependent. **(A)** qPCR analysis for stress-inducible genes; XBP1, XBP1 splice, BIP, GADD34 and CHOP. Transferring cell from normal high glucose/glutamine (Glc/Gln) 25 mM/4 mM to lower Glc/Gln (10 mM/1 mM) culture medium for 24h stimulated expression. *p <0.01 paired t test. **(B)** Survival after H_2_O_2_ treatment of SLC3A2 WT and KO HeLa cells cultured in normal and low Glc/Gln (10 mM/1 mM) medium with and without serum were measured with Alamar Blue. **(C-E)** N-glycans released by PNGase F from WT and SLC3A2 KO cell membranes were analyzed by LC-MS/MS; **(C)** Total N-glycans, (D) High-mannose and **(E)** Branched complex-type N-glycans with poly N-acetyllactosamine. Mean ± SD, * p <0.05 paired t test; 3 experiments with 2-3 technical replicates (See Table S3).

Global changes in N-glycan processing may be a feature of the stressed phenotype in SLC3A2 KO cells. Analysis of total cellular N-glycans released by PNGase F revealed lower levels of complex-type N-glycans and more high-mannose structures [Man(6 to 9)GlcNAc2] in SLC3A2 KO cells, consistent with a chronic increase in misfolded glycoproteins (**Fig. 3C-E, Table S2**). The removal of two glucoses from the initial N-glycan added by oligosaccharidyltransferase (OST) allows newly synthesized glycoproteins to enter the glucosidaseI/II Cnx/Crt cycle, a chaperone and quality control system in the ER. If folding fails after multiple cycles, a slow-acting ER α-mannosidase I cleaves the terminal mannose leaving Man(6 to 9)GlcNAc2, which targets the glycoprotein for degradation by ERAD (72). Poly-N-acetyllactosamine (poly-LacNAc) extensions to branched N-glycans were also reduced, perhaps due to ER stress impacting *trans*-Golgi β1-3 N-acetylglucosaminyltransferase activity. Elevated F6P and UDP-GlcNAc observed in the KO cells may be a response to stress, but it is insufficient to normalize ER and Golgi defects in N-glycosylation. Both the efficiency of N-glycosylation in the ER and Golgi remodeling of the SLC3A2 N-glycans are developmentally and stress regulated (5). This is explored further below (Heading: Reducing NXS/T site occupancy shifts SLC3A2 interactions and response to ER stress).

To summarize the above, SLC3A2 KO HeLa cells display decreased SLC7A5 dependent Leu/Gln exchanger activity, consistent with a decrease in intracellular essential AA and mTOR activity. SLC3A2 KO cells display an imbalance in central metabolism pathways and impairment of N-glycan processing, consistent with ER stress and increased AMPK activation. Resistance to ox-stress (H_2_O_2_) is shown to be dependent on Glucose/Gln supply in the medium and expression of SLC3A2. Next, we asked whether this dependence includes SLC3A2 N-glycans; their Golgi-modified structures in a position-dependent manner, and endocytosis by galectins.

### N-glycan branching is site-specific on SLC3A2

To assess the potential for SLC3A2 to interact with galectins, the N-terminal FLAG-tagged SLC3A2 wild type (abbreviated, WTseq), and gain-of-site FLAG-tagged variants A273N, D355N and D492N were inserted into the Flp-In site of SLC3A2 KO cells for expression under the control of a tetracycline (dox)-inducible promoter. Dox-induced FLAG-SLC3A2 levels were similar among the cell lines, and low in the absence of dox (**Fig. 4A**). Dox-induced FLAG-SLC3A2 rescued the stress response and resistance to H_2_O_2_ (**Fig. S5A,B**). FLAG-SLC3A2 variants displayed similar levels of dox-induced expression and increases in molecular weight consistent with N-glycosylation at A273N and D355N (**Fig. S5C**). However, D492N was unoccupied based on peptide analysis by LC-MS/MS and mobility in SDS-PAGE. Each site in FLAG-SLC3A2 displayed distinct and reproducible post-Golgi modified N-glycan profiles (**Fig. 4B-G, Tables S3**). The tetra-antennary content at N381 and N424 was higher than the other sites, and at N424, N-glycans were more frequently extended with linear poly-LacNAc chains. N-glycans at N506 were mostly bi- and tri-antennary, while N365 displayed predominantly tri- and tetra-antennary structures. The total LacNAc content ranked by site was N424 > N381∼ N365 > N506 (**Fig. 4I**). The endogenous SLC3A2 in WT HeLa cells displayed site specific N-glycan profiles consistent with that of dox-induced FLAG-SLC3A2 (**Fig. S6, Table S4**). Thus, the microenvironment of each N-glycan site uniquely influences the activity or access of Golgi remodeling enzymes.

**Figure 4.**
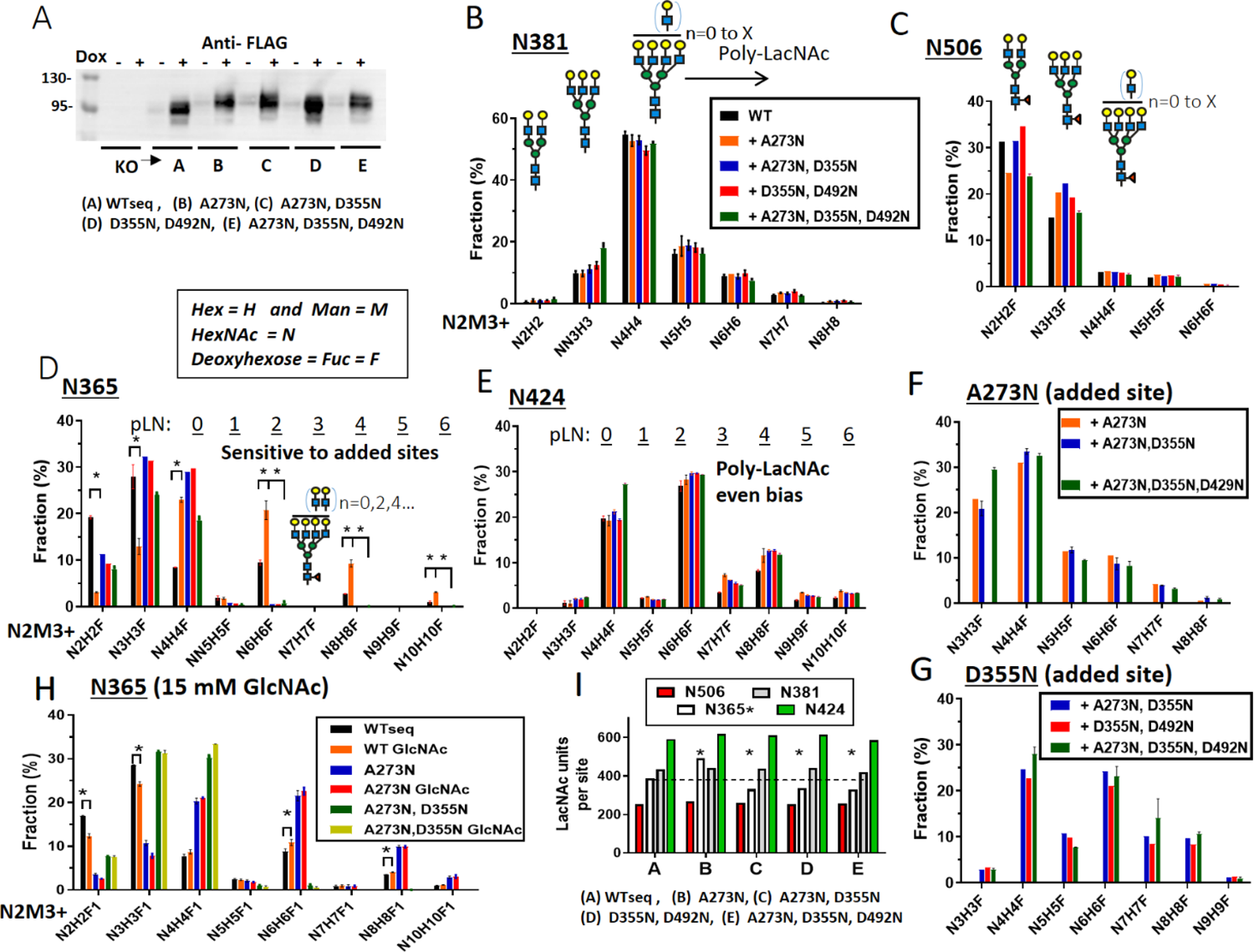
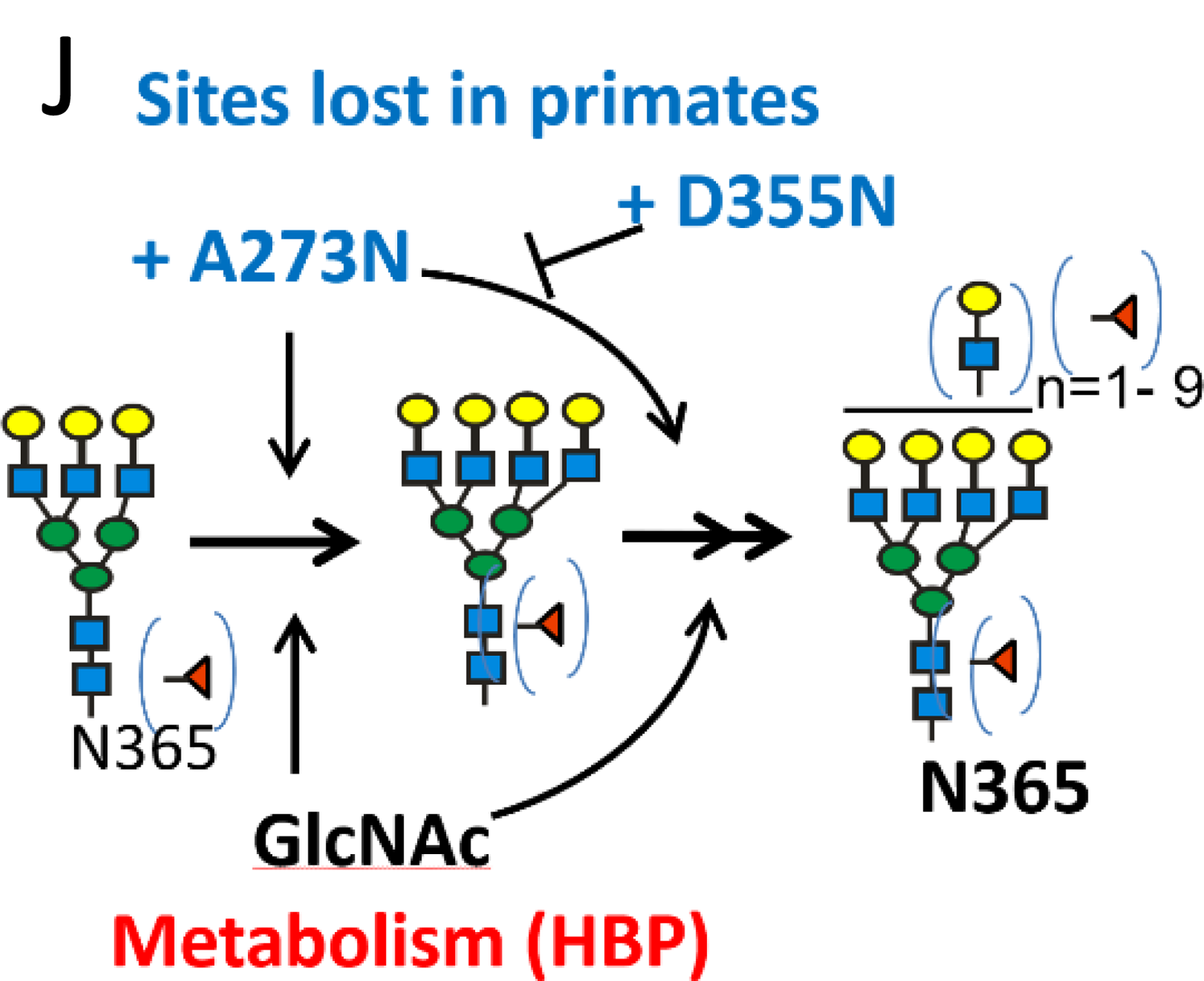
Site specific profiling of N-glycans on FLAG-SLC3A2 WTseq and gain-of-site variants. **(A)** SLC3A2 Western blot of anti-FLAG pulldowns from lysates of cells treated with and without dox. **(B)** Glycan structures at N381, **(C)** N506, **(D)** N365, (E) N424, and at added ancestral site (F) A273N, (G) D355N. The added site at 492 was not glycosylated. (Table S5). Mean ± SD of 3 experiments by LC-MS/MS with 3 technical replicates. **(H)** GlcNAc supplementation (48h) increases poly-LacNAc content on tetra-antennary N-glycans at N365 with the reciprocal decrease in bi- and tri –antennary glycans *p<0.05, n=3 (see Fig. S7). **(I)** N-acetyllactosamine (LacNAc) content at each site on FLAG-SLC3A2 in WTseq and gain-of-site variants. N365 LacNAc content is altered by added sites p<0.05, n=3. **(J)** graphical summary of positive and negative effect of added sites and GlcNAc on branching at N365.

Further LC-MS/MS fragmentation of the N-glycans provided additional information (**Fig. S7A-C**). We observed a remarkable bias towards even numbers of LacNAc units in poly-LacNAc chains at N424, N365 and the inserted site D355N, while N-glycans at N381 displayed the expected second order decay in elongated poly-LacNAc chains (**Fig. 4B-G, Table S3**). Unexpectedly, poly-LacNAc content at N365 was increased with the addition of A273N, while the co-addition of D355N suppressed the enhancing effect of A273N (**Fig. 4D,I**). Importantly, the N-glycan added at D355N completely suppressed the A273N enhancement and WTseq levels of poly-LacNAc at N365. N-glycan profiles for N424 and N381 were not significantly affected by the presence of the added sites. N365 was also the only site displaying increased branching when the cultures were supplemented with GlcNAc (**Fig. 4H, Table S5**).

Interactions with enzymes in the Golgi may depend on the intramolecular proximity of sites, as well as SLC3A2 heterodimer partners which assemble earlier in the ER. The distance from N365 to the enhancing site at A273N is ∼27Å, and N365 to inhibitory D355N site is ∼ 15Å. The results suggest that N-glycan remodeling at N365 has evolved away from a dependence on neighboring sites and directly on HBP flux and metabolism (**Fig. 4J, S8**). As discussed below, OST activity is also sensitive to metabolism whereby under-glycosylation of NXS/T sites in some substrates is an additional level of stress regulation (4).

### Site-specific regulation of both endocytosis and surface retention by galectin-3

SLC3A2 is both a positive and negative regulator of SLC7A exchangers. SLC3A2 is internalized by clathrin-independent endocytosis and tagged for lysosomal degradation by membrane-associated RING-CH E3 ubiquitin ligases (MARCH-1 or −8) (73). Mutation of Lys sites in the N-terminal cytoplasmic tail of SLC3A2 has been shown to increase T cells clonal expansion (32), suggesting that while SLC3A2*SLC7A5 stimulates growth, it is also a sensor of excessive metabolic rates. The extracellular domain of SLC3A2 projects branched N-glycans from N381 and N424 above the membrane by >100 Å where cytokine and adhesion receptors interact with the galectin lattice (7, 74). However, N365 is close to the membrane (∼10 Å), in proximity to glycolipids where Gal3 binding may promote internalization by GL-Lect mediated endocytosis (**Fig. 1C**). The N-glycan at N365 is also positioned for Gal3 crosslinking with solute transporters that are N-glycosylated on their short extracellular peptide loops. For example, Gal3 in pancreatic β cells binds N-glycans on the glucose permease SLC2A2 (10) and on the Arg transporter SLC7A3 (75), stabilizing both against endocytosis, thereby sensing elevated glucose and supplying Arg to the synthesis of nitric oxide, respectively, as required to stimulate insulin secretion.

To test the hypothesis that SLC3A2 N-glycans link membrane proximal and distal interactions, dox-induced FLAG-SLC3A2 WTseq, N381D and N365D site variants were tracked in live cells by immunocytochemistry. Approximately 80% of the FLAG-SLC3A2 expressed in HeLa cells was disulfide-linked as heterodimers (**Fig. S5D**). The levels of FLAG-SLC3A2 WTseq, N381D and N365D expressed on the plasma membrane after 48 h of dox induction were similar when stained with anti-CD98 (ie. SLC3A2) antibodies at 4°C (**Fig. 5A**). However, upon warming to 37 °C for 10 min, endocytosis of FLAG-SLC3A2 was observed with intensity N381D > N365D > WTseq, indicating a position dependent effect (**Fig. 5B**). N381D and N365D showed a more perinuclear accumulation, whereas the signal remained mostly peripheral in WTseq cells.

**Figure 5:**
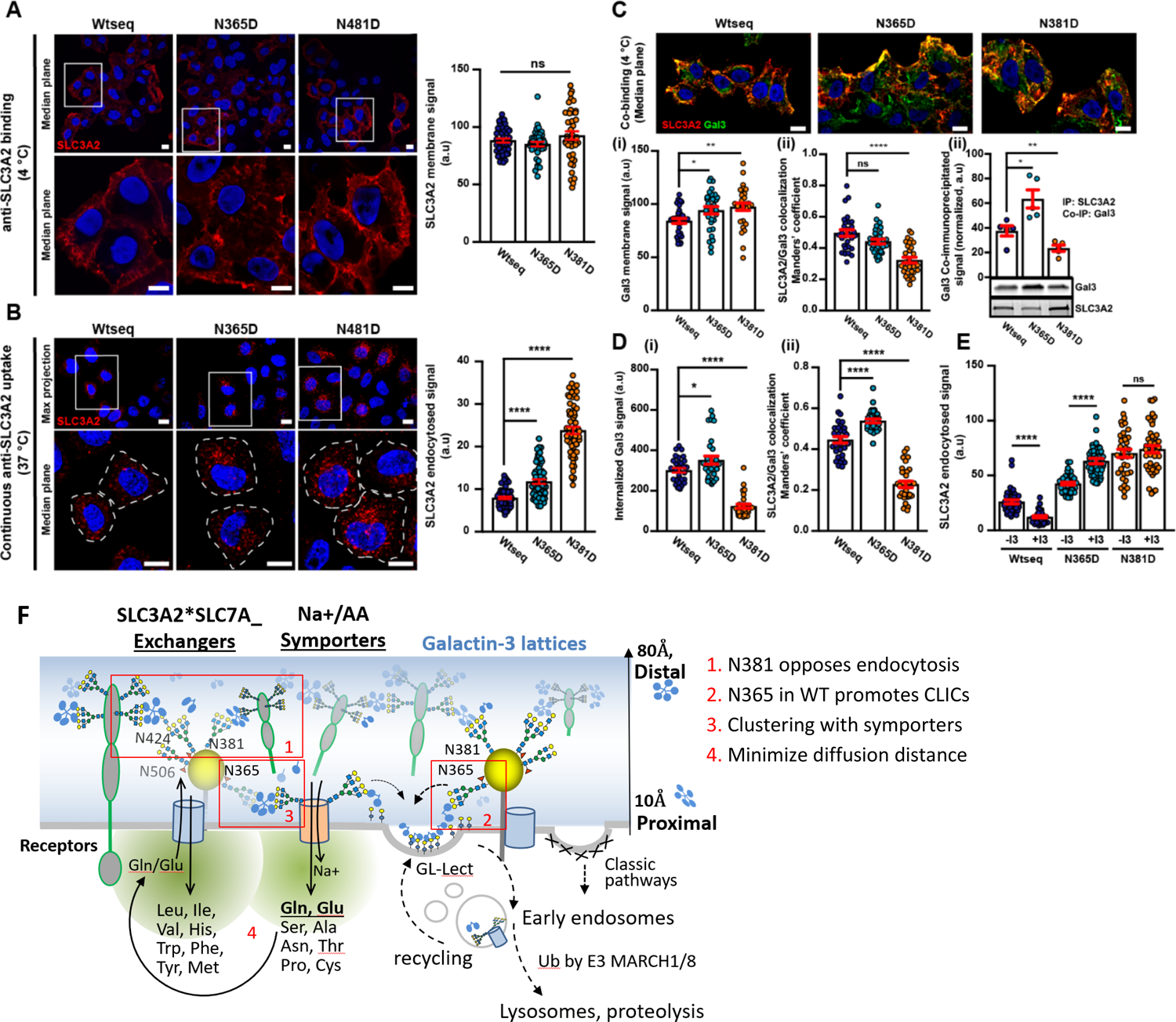
Regulation of SLC3A2 endocytosis. **(A)** Surface levels of FLAG-WTseq and variants are similar. Cells were incubated for 30 min at 4 °C with 10 µg/ml of anti-SLC3A2 antibody (red). Unbound antibody was removed with ice-cold PBS, cells fixed with 4 % PFA, and immunolabeled. The SLC3A2 signal was quantified (right histogram). Zooms are from boxed areas above. Nuclei in blue (DAPI). Scale bars = 10 µm. **(B) Internalization of SLC3A2 site variants is increased.** Cells were continuously incubated for 10 min at 37 °C with 10 µg/ml anti-SLC3A2 antibody (primary antibody), and then shifted to 4 °C. Cell surface-accessible antibody was removed on ice with acid wash solution. Cells were then fixed, permeabilization with saponin, immunolabeled, and images were quantified ****p<0.0001. A more perinuclear accumulation of the signal from internalized antibody was observed in N365D and N381D expressing cells, whereas the signal remained mostly peripheral in WT cells. **(C) Co-distribution of SLC3A2 and Gal3.** Cells were incubated for 30 min at 4 °C with 5 µg/ml of purified Gal3-488, and after washing for 30 min at 4 °C with 10 µg/ml anti-SLC3A2 antibody (primary antibody). After extensive washes with ice-cold PBS, cells were fixed and immunolabeled with secondary antibody. Gal3 signal at the membrane and its colocalization with SLC3A2 were quantified in the histograms. (i) Surface levels of Gal3 are similar; (ii) Mander’s coefficient dropped significantly for the N381D variant; (iii) co-immunoprecipitation with Gal3 was increased for N365D and decreased for N381D. **(D)** Same as previous with the additional 10 min at 37°C for measuring internalization. (i) Endocytosis of Gal3 is increased in N365D expressing cells and decreased in N381D expressing cells; (ii) Mander’s coefficient of Gal3 with SLC3A2 is increased in N365D expressing cells and decreased in N381D expressing cells. **(E) Gal3 inhibition and SLC3A2 endocytosis.** Cells were preincubated for 5 min at 37 °C with 10 µM of Gal3 inhibitor GB0149-03. Inhibitor solution was removed, and cells were continuously incubated for 10 min at 37 °C with 10 µg/ml of anti-SLC3A2 antibody, placed on ice to stop endocytosis, acid-washed to remove cell surface-accessible antibodies, fixed, permeabilized and immunolabeled. Internalized SLC3A2 signal quantified. **(F)** Working model. Surface expression, trafficking and interactions of SLC3A2*SLC7A5/11 regulated by Gal3 are highlighted in red 1 to 4. Clathrin-independent carriers (CLIC).

Next, we asked whether N-glycan positions influenced galectin binding. Fluorophore tagged Gal3-488 was added to cells and binding at 4°C was again similar (**Fig. 5C(i)**), although the Manders coefficient for co-localization (clustering) was decreased in N381D expressing cells (**Fig. 5C(ii)**), consistent with loss of the higher affinity tetra-antennary N-glycans at N381. Moreover, anti-SLC3A2 antibodies also pulled-down less Gal3-488 in lysates of N381D expressing cells, and surprisingly, more from N365D lysates compared to WTseq (**Fig. 5C(iii)**). After washing to remove excess Gal3, addition of anti-SLC3A2 antibody and warming to 37°C, Gal3-488 uptake was decreased with N381D, when compared to the WTseq, while uptake was increased with N365D compared to WTseq (**Fig. 5D(i)**). Furthermore, co-internalized Gal3 and SLC3A2 colocalized strongly in the N365D, but not in the N381D expressing cells (**Fig. 5d(ii)**). consistent with the co-immunoprecipitation data in **Fig. 5C**.

To explore this idea further, cells were pre-incubated (5 min) with a Gal3 inhibitor GB0149-03 (76, 77) in the same assay protocol. GB0149-03 treatment further increased the already augmented endocytosis of the N365D variant, but no additional effect on already high level of internalization of the N381D variant (**Fig. 5E**). This again was consistent with a decreased interaction between the Gal3 and the N381D variant. In sharp contrast, GB0149-03 inhibited uptake of WTseq by ∼50%, consistent with a requirement for Gal3 in the formation of tubular endocytic pits, GL-Lect mediated endocytosis and internalization via clathrin-independent carriers (78). These findings strongly suggest that N-glycans at the membrane-distal position N381 are dominant for surface retention by the galectin lattice, slowing loss to endocytosis by other mechanism (6). In contrast, the N-glycan at N365, nearest the membrane is critical for GL-Lect mediated endocytosis associated with glycolipid-enriched raft domains (79).

The localization of the endocytosed N365D and N381D variants to perinuclear endosomes instead of peripheral ones (as observed with WTseq; **Fig. 5B**) indicates a reduced affinity for the galectin lattice resulting in a loss of SLC3A2 internalization by other pathways (6), thereby masking or rendering the GL-Lect mechanism nonfunctional. To test this more directly, we compared the endocytic fate of SLC3A2 WTseq under 2 conditions: (i) acute GB0149-03 or lactose treatment of 5 min versus (ii) extended treatment of 30 or 60 min. As already described in Fig. 5D, the acute treatment protocol led to ∼50% inhibition of SLC3A2 WTseq endocytosis (**Fig. S9A**). In contrast, SLC3A2 WTseq endocytosis was increased under the conditions of the extended treatment protocol (**Fig. S9A**). As with the N365D and N381D variants, extended treatment drove a significant fraction of the SLC3A2 WTseq into perinuclear endosomes in which SLC3A2 WTseq overlapped more strongly with the early endosomal marker EEA1 (**Fig. S9B**). The results suggest that the galectin lattice and the GL-Lect mechanism act as opposing but complementary forces on SLC3A2 (**Fig. 5F**). The fate of SLC3A2*SLC7A trafficking to lysosomes or recycling endosomes may be regulated by changing affinities for galectins, particularly at N365, where N-glycan branching is sensitive to the HBP and microenvironment.

### SLC3A2 interacts with transporters and other glycoprotein adaptors: SLC3A2 interacts with transporters and other glycoprotein adaptors

SLC3A2 interactions with SLC7A transporters and integrin-β1 are well documented (25, 48), but other interactions and the role of N-glycans are largely unknown. As a first step, affinity purification - mass spectrometry (AP-MS) analysis of dox-induced FLAG-SLC3A2 WTseq identified 155 significantly enriched proteins (Fig. 6A Table S6). Dox-induced FLAG-SLC3A2 rescued the stress-response and resistance to H_2_O_2_, indicating that the protein interactions observed were that of a functional bait (Fig. S5A,B). Dox-induced expression and pull-down efficiencies were similar for wild-type and variants (Fig. S5C,D). By gene ontology (V11.0), the list of interactors included 115 integral components of membranes, of which 70 were nutrient and ion transporters, including AAs, monocarboxylate, ATPase Na^+^/K^+^ exchangers and, Zn^++^ transporters (Fig. 5B). The top interactor was SLC7A5 by peptide intensity. The transmembrane transporters represented ∼13 subfamilies of SLCs and mitochondrial transporters. The set of 155 proteins also included eight complex V ATP synthases (ATP5 family), ADP/ATP translocator (SLC25A5, SLC25A6), Asp/Malate and αKG/malate shuttle (SLC25A11), thiamine pyrophosphate transport (SLC25A19) and Glu transporters (SLC25A22, SLC25A13, SLC25A11). Other proteins in the list appear to be novel interactors, that may be dependent on dox-induced FLAG-SLC3A2 rescue of the KO cells. These include a mitochondrial transmembrane transport complex; purine ribonucleotide biosynthesis, protein N-glycosylation, and lipid biosynthesis-associated proteins SLC3A2 interacted with other glycoprotein adaptors; CD44, BSG, ATP1B1 and ATP1B3, which are regulated by clathrin-independent carriers, and partner with non-glycosylated transporters (78, 80) (Fig. 6B). GO:KEGG points to diseases associated with oxidative and mitochondrial stress (Huntington, Parkinson, Alzheimer, Diabetic cardiomyopathy) (Fig. 6D,E).

**Figure 6.**
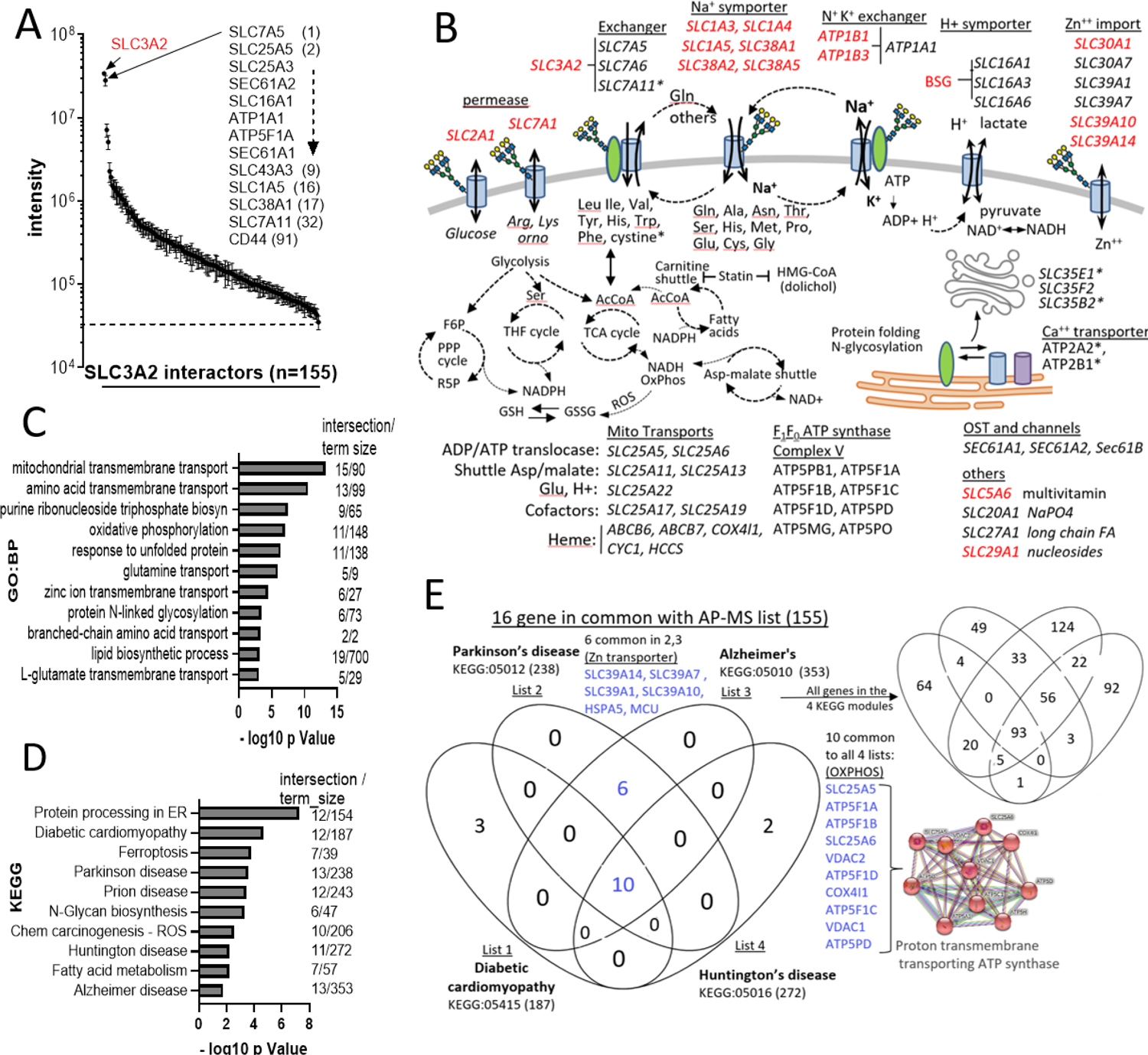
SLC3A2 interacting proteins. **(A)** AP-MS with anti-FLAG antibodies in dox-induced FLAG-SLC3A2 cells revealed peptides from 155 interacting proteins with mass intensities > 3 SD above control pulldowns from dox-treated SLC3A2 KO cell lysates (mean ± SD, n=5 experiments). The results are plotted in rank order of peptide intensity. **(B)** SLC3A2 interacting proteins with potential for regulation of AA and ion gradients impacting metabolism. N-glycosylated proteins are marked in red. SLC3A2, ATP1B, ATP1B3 and BSG are all co-receptors that partner with non-glycosylated transporters. Gene ontogeny **(C)** Biological Process and **(D)** KEGG by gProfiler using ordered query for the 155 FLAG-SLC3A2 interacting proteins. **(E)** Sixteen SLC3A2 interacting proteins associated with four degenerative diseases, which segregate into two functional classes (Venny Diagram, J.C. Oliveros). The STRING network represents the 10 transmembrane protein transporters. Four SLC39A transporters regulate zinc import/export.

### SLC3A2 N-glycans are required for interactions with AA/Na^+^ symporters

Anti-FLAG pulldowns effectively depleted FLAG-SLC3A2 from cell lysates, and ∼75% of the FLAG-SLC3A2 was disulfide-linked as heterodimers (**Fig. S5D**). We focused on 85 of the 155 SLC3A2 interactors that displayed significant differences between wild type and site variants in a Kruskal Wallis test of all and in pairwise comparisons. With N381D and N365D site mutations, FLAG-SLC3A2 interactions decreased including SLC7A5 by 44±14% and SLC7A11 by 67±15%; as well as neutral AA/Na^+^ symporters SLC1A5, SLC1A4, SLC38A2 and SLC38A1; Arg/Lys/Ornithine high-affinity permease SLC7A1; the Zn^++^ transporters SLC30A1; and non-catalytic subunit of Na^+^/K^+^ ATPase-dependent exchanger ATP1B1 (**Fig. 7A, Table S6**). Importantly, SLC1A5, SLC1A4, SLC38A2, SLC38A1, SLC7A1 and SLC30A1 are N-glycosylated, and have also been identified as prey correlating with SLC3A2 in a global BioID proteomics network (81). The BioID method depends on proximity over time rather than affinity and can therefore detect dynamic interactions as they occur in phase transition condensates such as the galectin lattice (17, 82). Experimental and inferred interactions between SLC7A exchangers and AA/Na^+^ symporters occur frequently in the literature, perhaps in this case by dynamic galectin crosslinking, which is nearer the affinity cut-offs in most proteomics methods.

**Figure 7.**
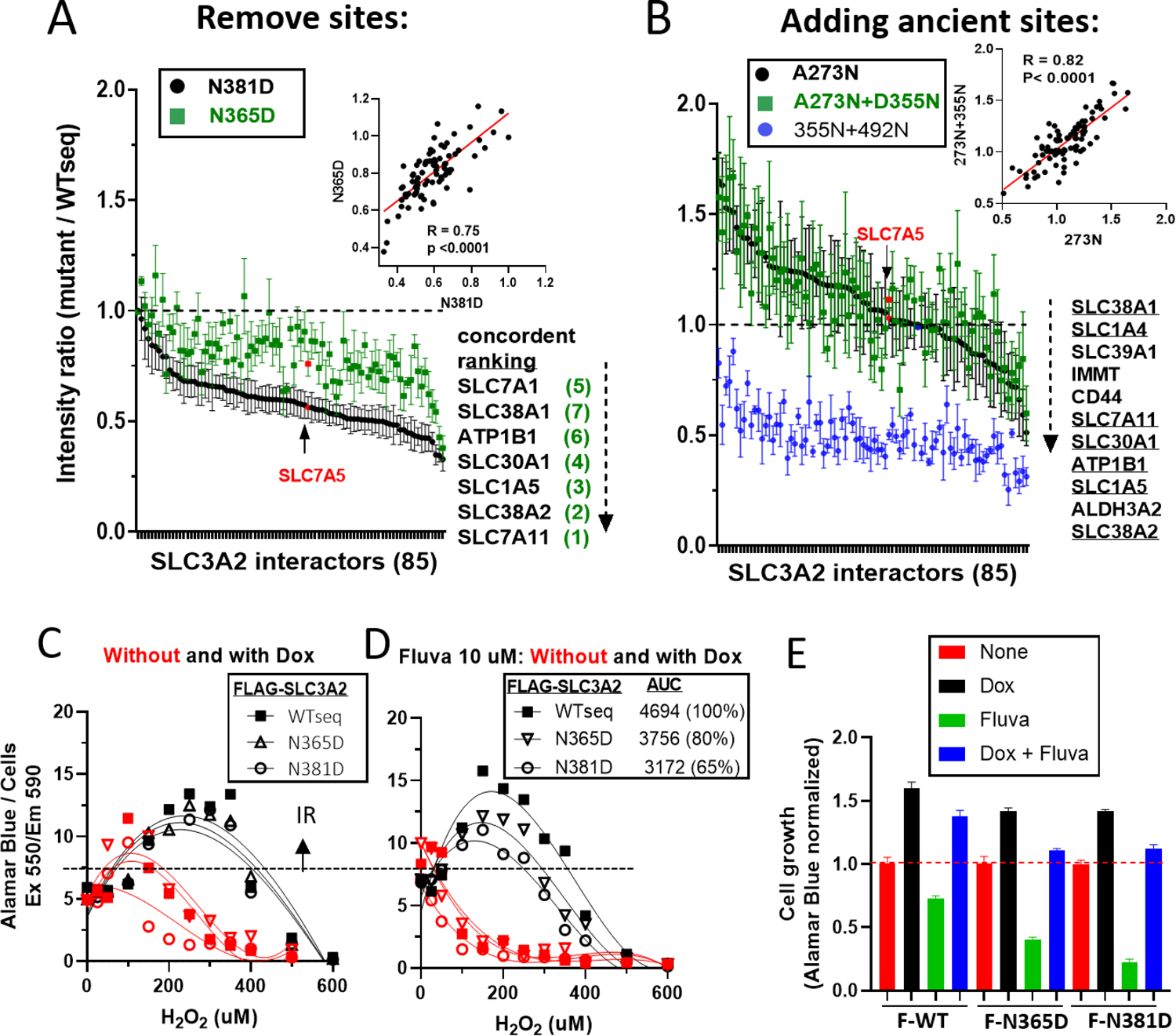
Protein interaction comparing FLAG-SLC3A2 WTseq and site variants as baits. **(A)** AP-MS analysis of anti-FLAG antibody pulldowns from dox-induced FLAG-SLC3A2 WTseq and site-deletion variants, shown as mutant/WTseq ratios, mean ± SE, WTseq, N381D (n= 8 each). Graphed relative to ranking of N381D/WTseq. **(B)** Ratios from WTseq n=4, A273N n=3, A273N+D355N n=3, and D355N+D492N n=3 independent pulldowns. Graphed relative to ranking of A273N/WTseq (listed in descending order for all three variants, and <u>seven</u> of the most decreased are common to N381D and N365D variants in panel A. Inserts shows correlations between variant pairs. For the variant in blue, added sites D355N was glycosylated and D492N was not. **(C,D)** Cells pre-treated +/- dox for 2 days to induce expression of FLAG-WTseq and variants, and the last 24h without **(C)** and with **(D)** 10 *μ*M fluvastatin followed by H_2_O_2_ for 16h. The Alamar Blue signal is normalized to viable cell number by InCell imaging. **(E)** Growth of cells +/- dox-induced FLAG-SLC3A2 +/- 10 *μ*M fluvastatin for 24h.

SLC3A2 with sites added, A273N and A273N+D355N, also showed reduced association with many of the same proteins including SLC38A2, SLC1A5, ATP1B1, SLC30A1 SLC7A11, SLC39A1, SLC1A4 and SLC38A1(**Fig. 7B, Table S6**). However, unlike the site deletion variants, site insertion at A273N increased SLC3A2 association with ER chaperones (i.e., HSPA5, HSP90AB1, HSP90AA1 and HSPA8) indicating impaired SLC3A2 folding, heterodimer formation and/or additional ER stress in cells rescued by these variants. N-glycan addition at D355N suppressed interactions with most of the 85 candidates (**Fig. 7B**). Importantly, mutations at N381, N365 or gain of ancestral sites decreased SLC3A2 association with SLC7A11 and a subset of glycosylated AA /Na+ symporters associated with the response to ox-stress.

### Reducing NXS/T site occupancy shifts SLC3A2 interactions and response to ER stress

The set of 155 SLC3A2 interacting proteins included OST proteins (STT3A, DDOST, RPN1, RPN2), and the channel-forming translocon (SEC61A2, SEC61A1, SEC61B), suggesting an interaction with the initial steps of N-glycosylation in the ER. Glucose starvation depletes the donor substrate for OST, lipid-linked oligosaccharide (Glc_3_Man_9_GlcNAc_2_-pp-dolichol), leading to ER stress and the unfolded protein response (4). Statins inhibit HMG-CoA reductase, thereby lipid-linked oligosaccharide biosynthesis (83, 84), providing an alternate means of reducing N-glycosylation independent of starvation (85). We treated cells with fluvastatin for 48h, then measured changes in N-glycosylation and the antioxidant capacity of SLC3A2 WT and KO cells (**Fig. S10A**). Titration with increasing H_2_O_2_ revealed an early transient increase in the Alamar Blue signal (i.e., NAD(P)H / viable cell) in WT cells, indicating an induced resistance which was reduced in SLC3A2 KO (**Fig. S10B**). Fluvastatin treatment suppressed this induced resistance to H_2_O_2_ in both KO and WT cells. In low glucose/Gln conditions, the induced response was suppressed in both KO and WT cells, and Fluvastatin treatment had no additional effect (**Fig S10C**). The SLC3A2 KO and fluvastatin treatment caused a similar imbalance observed in AAs, glycolysis, HBP, GSH and purine pathways (**Fig. S11**).

We recently reported that fluvastatin treatment reduces N-glycosylation at N365 and N381, but not at the N424 site (5), suggesting these SLC3A2 sites have an evolved OST sensitivity to stressors that reduce lipid-linked oligosaccharide levels. Thus, fluvastatin treatment is expected to interact with the SLC3A2 single-sites loss variants to impair stress mitigation. Indeed, rescue of the induced-response to H_2_O_2_ in the presence of fluvastatin was comparatively WTseq > N365D > N381D, consistent with a requirement for N-glycosylation at both sites to promote SLC3A2-heterodimer surface residency and/or function (**Fig. 7C,D,E**).

In SLC3A2 KO cells rescued by dox-induced WTseq FLAG-SLC3A2, fluvastatin treatment increased SLC3A2 association with AA transporters and ER protein chaperones (CANX, HSPB1, TMX3, SEC61A1, SEC61A2, HSPA5, HSP90AB1, HSP90AA1, HSPA8), which were also increased with the A273N gain-of-site variant (**Fig. S10D,E, Table S6**). CD44 was one of the most enhanced SLC3A2 interactors, ranking 4^th^ in fluvastatin treated cells, and is known to bind SLC3A2*SLC7A11 under stressed conditions, thereby promoting surface expression, import of cystine and stress mitigation (86, 87). The long splice form, CD44v has N-glycan sites in the neck region nearer the membrane that are required for binding to SLC3A2*SLC7A11 (86). Although N381 and N365 sites in SLC3A2 are hyper-sensitive to statin-induced suppression of N-glycosylation (5), the loss of N-glycans at these sites may allow recruitment of CD44v to SLC3A2*SLC7A11 (86), thereby compensating in the galectin lattice to recruit the AA/Na^+^ symporters required for GSH synthesis (Gln, Glu, Cys and Ser). Indeed, six of the enhanced interacting transporters in fluvastatin-treated cells (SLC7A1, SLC7A11, SLC1A4, SLC38A1, SLC1A5 and SLC38A2) were also identified as depleted with N365D and N381D variants in non-stressed conditions (**Fig S10E**). Furthermore, a gProfiler multiquery ordered by intensity ratios to compare data in **Fig. S10D and Fig. 6A**, revealed GO:BP modules consistent with cell death, AA transport and ATP biosynthesis (**Fig. S10F**). Presumably, upon reversal of ox-stress, SLC3A2 and CD44 can be internalized by GL-Lect endocytosis thereby normalizing the levels of the associated transporters (78,80,88).

### Purging and duplication of genes in Neoaves evolution

The report by Castiglione et al. (89) on the absence of KEAP1 in Neoaves drew our attention to the possibility of extending their observations. Neoaves are modern volant bird, representing ∼95% of all Ave species. KEAP1 encodes an E3 ubiquitin ligase that targets the transcription factor NRF2 for proteolysis, a key suppressor of antioxidant gene expression (89). Thus, loss of KEAP1 allows constitutive NRF2-driven expression of SLC7A11, glutathione synthesis (GCLC, GCLM) and thiol redoxin proteins (TXNRD1, PRDX1). Castiglione et al. (89) suggested that an enhanced NRF2 antioxidant response may lower the risk of oxidative damage associated with higher metabolic rates in Neoaves and contribute to their longevity. With this connection to the SLC7A11 transporter, we noticed that SLC3A2 is also absent in Neoaves suggesting further adaptation that includes both BCAA and cystine uptake. Indeed, SLC3A2 function connects these two recognized features of metabolism associated with longevity: - regulation of BCAA levels - thereby mTOR activity, and cysteine flux to GSH synthesis. C4 to C6 acyl-CoA biproducts of Leu, Ile, Val, Phe, and Tyr catabolism can compete for carnitine and interfere with fatty acid metabolism (50,90,91), which is critical for both fat stores and fueling muscle in prolonged flight. Indeed, reducing dietary BCAA in mice increases fatty acid catabolism, reduces triglyceride stores, and improves insulin sensitivity (92). However, essential AA catabolism has a remarkably higher level of complexity. Catabolism of Ile, Leu, Val, Met, and Thr generates acyl-CoAs which are labile and form reactive intermediates that non-enzymatically modify Lys residues in mitochondrial enzymes and nuclear proteins (91, 93). Although poorly understood, these modifications appear to be dependent on local concentrations of the auto-catalytic acyl-CoAs, and turnover of acyl-Lys sites occurs through deacetylation by sirtuins (SIRT1-7).

Briefly, the progenitors of birds survived the Permian–Triassic extinction 250 million years ago (MYA) that caused one of the greatest loss of species (∼95%) under sever anoxic conditions, and it took millennia to restore atmospheric oxygen and species diversity. The evolution of birds from bipedal dinosaurs involved a progressive reduction in body size, where the energy costs of being large were traded for those of being endothermic and adapted for flight (94, 95). The benefits were greater mobility and foraging opportunities, predator avoidance, and tolerance to a range of environments (96). Body mass correlates well with lifespan and metabolic rate in vertebrates, however birds are outliers often showing greater longevity and higher metabolic rates with smaller masses (96). Phylogenetic analysis of modern birds (Neoaves) suggests a massive protein-coding sequence convergence, and incomplete lineage sorting during rapid radiation after the Cretaceous-Paleogene mass extinction event ∼66 MYA (97). Selective pressures drove changes in physiology and metabolism and a surprising ∼25% reduction in genome size, discarding many genes and duplicating a few others (98–100). Evolution generally makes use of mutations in existing genes, and less frequently gene inactivation and losses occur when their contribution to fitness wanes. However, Neoave evolution has apparently been driven by selective purging of genes (97), perhaps reducing network complexities (gene paralogues) and streamlining metabolism (compartmentalize) thereby improving energetics.

The canary (Serinus canaria, a Neoave) and chicken (Gallus gallus, a Neoave progenitor), have an estimated 15,281 and 17,478 protein encoding genes, respectively. Human and mouse (Mus musculus), separated by a similar ∼80 MYA, have ∼22,389 and ∼23,317, respectively, with > 98% functionally annotated by homology (101). Gene loss in Serinus canaria for 47 KEGG modules of interest is 17.4 ± 3.67% (624 of 3730 genes) (**Fig. S12A, Table S7**). Greater gene loss is observed in pyruvate metabolism (KEGG:00620); glycolysis/gluconeogenesis (00010) and oxidative phosphorylation (00190). Conserved KEGG modules with more genes retained included: HBP nucleotide-sugars (00520); N-glycan biosynthesis (00510); fatty acid biosynthesis (01212); mTOR pathway (04150); ubiquitin proteolysis (04120). HBP enzymes, ER and Golgi N-glycan processing, as well as galectins LGALS1, 2, 3 and 8, are conserved in Neoaves, and duplications MGAT4B and MGAT4C point to the importance of branched N-glycans biosynthesis, as discussed below (**Fig. S12C**).

#### Saving on energy expenditure

Three activities contributing to ∼75% of energy expenditure in non-dividing cells: (1) protein turnover, (2) Na^+^/K^+^ ATPase exchanger and (3) mitochondrial proton leakage (102), have been addressed with Neoave evolution as indicated by gene losses. Protein turnover in mammalian cells is estimated to account for ∼25% of energy expenditure. At scale, gene purging and reduction in total base-pairs (99) reduces the energy cost of maintaining a larger than needed genome and proteome. Indeed, lifespan is negatively correlated with rates of protein turnover measured in confluent cultures of primary fibroblasts from diverse mammalian species. Swovick at al. (103) conclude that fast protein turnover is sufficient in shorter-lived organisms, but the consumption of ATP and generation of ROS over time would disadvantage longer-lived species. Thus, the beneficial effects of protein turnover on proteostasis are balanced against the energy costs of protein re-synthesis and the associated cost in damage by ROS.

However, beyond a smaller proteome in Neoaves, selective genes losses and duplications have contributed to efficiencies in ATP utilization and ROS mitigation. Another ∼25% of energy allocation is accounted for by ATPase-driven Na^+^/K^+^ exchange (102), which is functionally linked to SLC7A exchangers in mammals (**Fig. S3A,F**). Export of Glu/Gln with the import of essential AA by SLC3A2*SLC7A exchangers (26) is accompanied by recovery of Glu/Gln by AA/Na^+^ symporters, which requires the energy-consuming ATPase Na^+^/K^+^ exchanger to maintain the Na^+^/K^+^ gradient across the membrane (**Fig. 8A**). SLC7A exchangers mediate bidirectional flux in real time (milliseconds) balancing essential AA and continuously consuming ATP in this manner. Selection against what appears to be a burden on energetics is supported by losses of SLC7A7, SLC7A8 and severe truncations of SLC7A5 and SLC7A10 exchangers in Neoaves (**Fig. S12C, S13**). Two other sodium-dependent large AA transporters, SLC43A1 and SLC43A2 are also absent, leaving only SLC6A15 as a known transporter of BCAA found in mammalian brain. In human tumors, increases in SLC3A2 and SLC7A5 expression are accompanied by SLC1A4 and SLC1A5, the interacting Glu/Gln symporters, which are also absent in Neoaves. Thus, purging nine genes encoding transporters that interact strongly supports selection against the coupled mechanism of essential AA regulation by SLC3A2*SLC7A exchange - Glu/Gln AA/Na^+^ symporter - ATPase Na^+^/K^+^ exchange.

**Figure 8.**
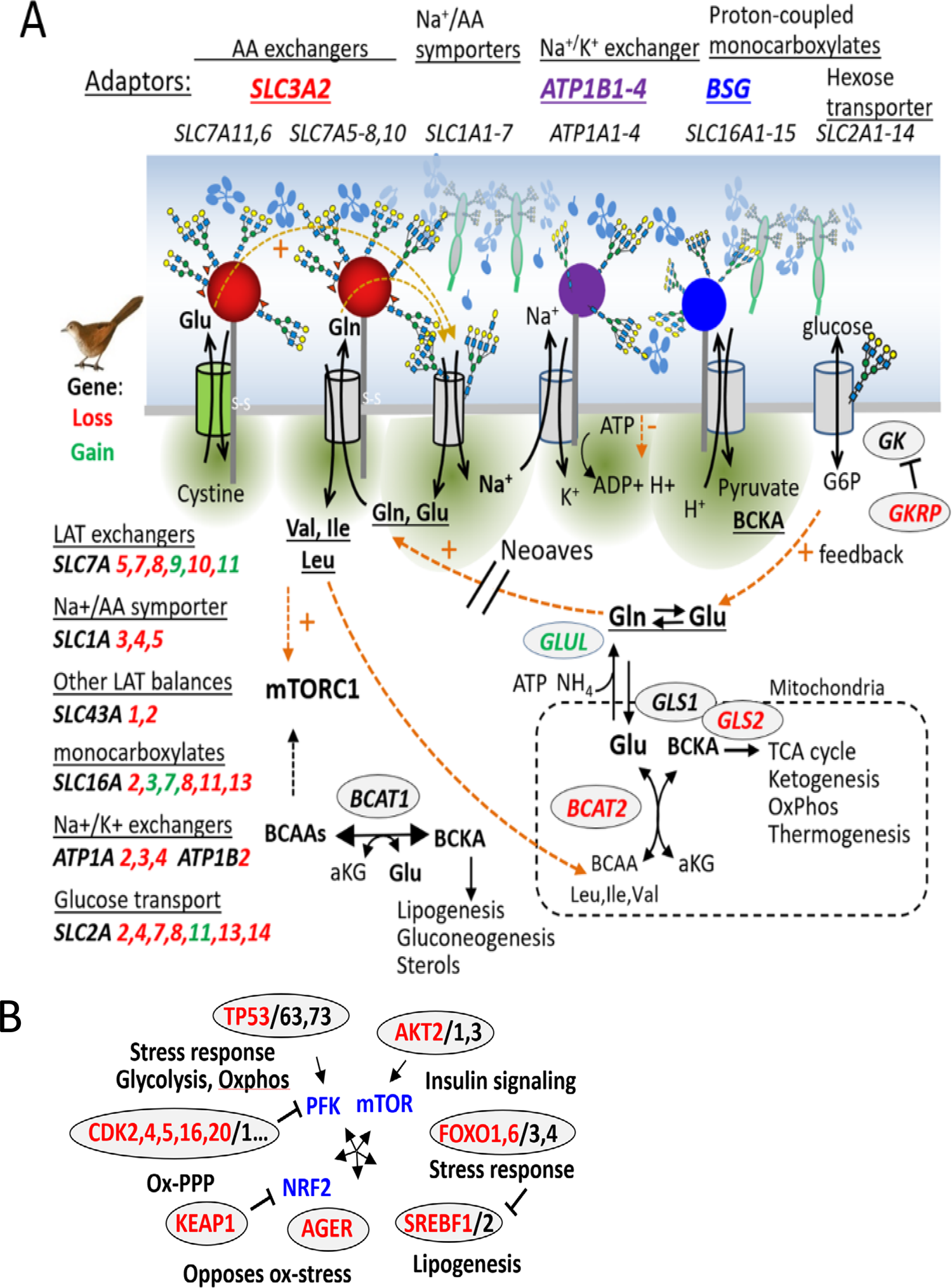
Positive feedback model in mammals is deleted in birds. **(A)** Interactions of the mammalian network with positive feedback regulation (dashed arrows) by SLC7A exchanges and adaptor mediated clustering. Intracellular Glu and Gln drives essential AA import, mTOR activation and catabolism of BCAA. Gene purging with the evolution of Neoaves has rewired the AA transporter network, redox regulation and signaling (see Fig.S14 for gene-enzyme list). **(B) Genes lost in birds** (red) are often one or more paralogues with overlapping activities and tissue or organelle selectivity (see Fig. S14). Genes shown are of interest in cancer, metabolic disease, inflammation, and aging. Tumor protein (TP53), RAC-alpha S/T protein kinase (AKT), Forkhead box protein O, (FOXO), Sterol regulatory element-binding protein (SREBF), Advanced glycosylation end product-specific receptor (AGER), Kelch like ECH associated protein 1 (KEAP1), Cyclin-dependent kinase (CDK1-20).

If fitness has been improved by conserving ATP, then losses >25% in other ATPases-dependent transporter families might also be expected. Indeed, 38% (16/42) ATPase dependent ABC transporters (KEGG:02010) are absent or severely truncated. Remarkably, 85% (6/7) of the purinoreceptors (P2RX1-7), a family of ATPase-gated cation-permeable ion channels (Ca^++^/K^+^/Na^+^) are also missing in Neoaves. Only P2RX5 is well conserved in birds (∼93%) and upstream ATP-channel (PANX1-3), downstream ectonucleotides (ENPP-1, 3) and receptor (P2RY-1, 2) are present suggesting the pathway is intact.

#### BCAA metabolism and mTOR

Mitochondrial BCAA aminotransferase (BCAT2) is also absent, leaving only cytoplasmic BCAT1 as gatekeeper for BCAA catabolism. The BCATs catalyze bidirectional transamination between BCAA + α-ketoglutarate and branched chain α-keto acids (BCKA) + Glu. In BCAT2 deficient mice, plasma BCAA levels and energy expenditure are increased, adiposity decreased, and insulin sensitivity improved (104). Endurance is reduced with the loss of BCAT2 or SLC7A5 (105, 106) consistent with the known contribution of BCAA to oxidative respiration in mammalian muscle (53, 92). However, endurance in Neoaves may be enhanced by deleting the SLC7A exchanger mechanism, reducing BCAA uptake and oxidation in muscle, where fatty acids and ketones are the preferred fuel (**Fig. 9A, S14**). Indeed, fat is anhydrous and highly reduced with a 6-fold higher density of stored energy than glycogen. Transport and exchange of monocarboxylates including BCKAs and ketones are mediated by SLC16A family members present in Neoaves, including SLC16A3 and SLC16A7 present as two and three paralogs in Neoaves, respectively. BCKA may largely support hepatic lipogenesis, spared from oxidation in muscle and re-aminated by BCAT1 in other tissues as needed for protein synthesis. The absence of BCAT2 may reduce ammonia release in the mitochondria and its potential toxicity. Ammonia is directed into the purine pathway to uric acid, by glutamine synthetase (GLUL) - duplicated in bird, an enzyme that catalyzes the ATP-dependent conversion of NH_4_ and Glu into Gln. Bird physiology conserve water by shifting ammonia disposal to this less soluble end products. Bird guano contains ∼10% protein which is needed to package uric acid at ∼80% in colloidal suspension, whereas loss of protein and AAs in the urine of mammals is minimal. This loss is compensated for by a shift away from the use of glucuronidation to remove water insoluble compounds (17/20 UDP-glucuronosyltransferases are absent in birds).

**Figure 9.**
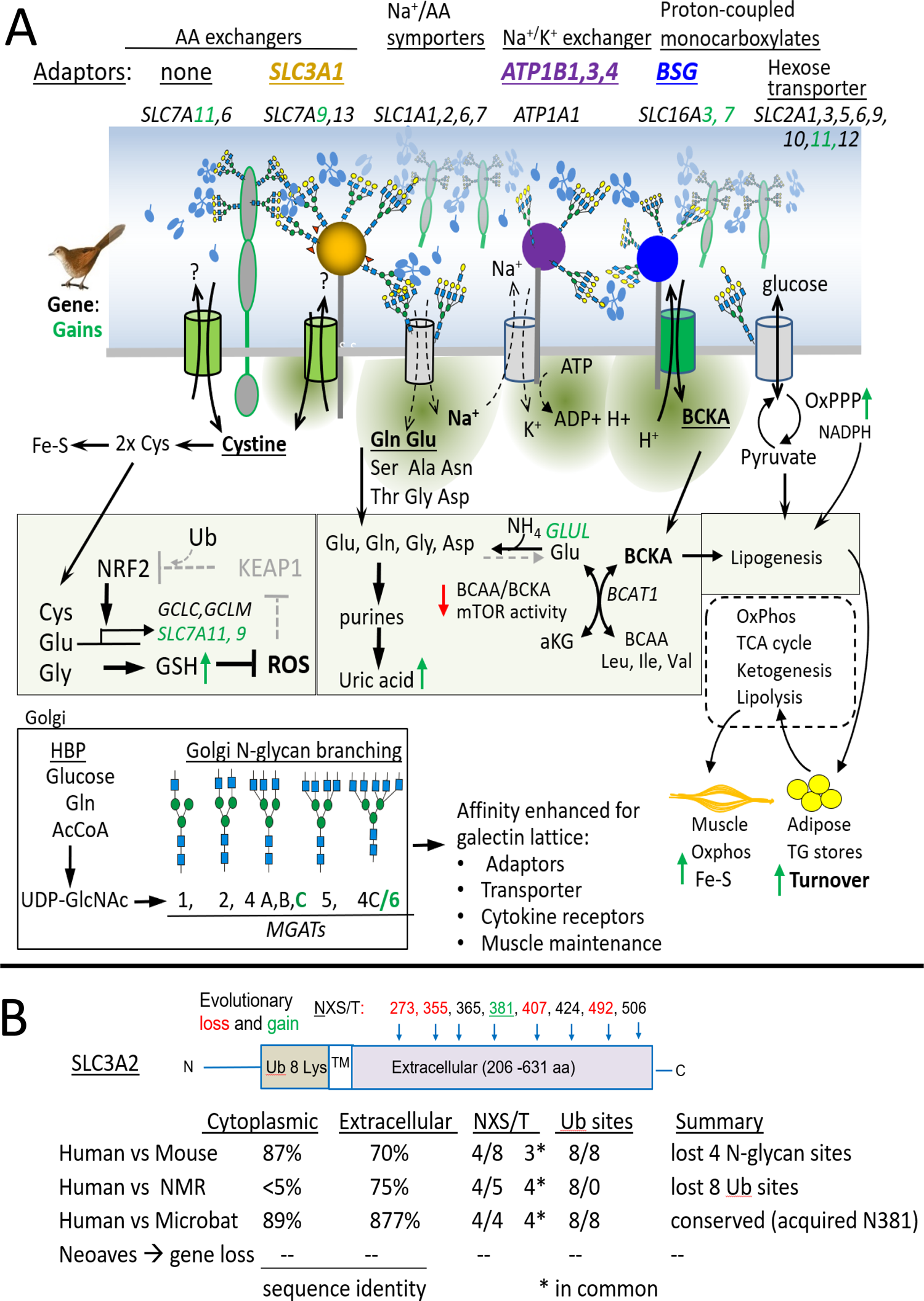
**(A) Alternate network of adaptors and transporters in Neoaves.** The interaction of SLC3A1 with ATP1B1,3,4 and BSG adaptors promotes BCKA transporter and hepatic AA flux into fatty acids to fuel muscle during flight. Galectins may regulate the transporter network in Neoave, with a greater dependence on duplicated genes: SLC7A9, SLC7A11 for cystine (into GSH and Fe-S complex), and SLC16A1 and SLC16A3 for BCKA uptake and exchange. HBP and N-glycosylation pathways are well conserved and increased branching by duplication of MGAT4B and MGAT4C supports a higher affinity of membrane glycoproteins for galectins. Without SLC3A2 and Gln/Glu driven SLC7A exchangers, energy is saved by reducing the activity of ATPase-driven Na^+^/K^+^ exchanger, and BCAT1 control of BCAA flux in the cytoplasm, reducing its oxidation and mitochondrial stress. **(B) Divergence in SLC3A2 evolution.** Comparing the extracellular and cytoplasmic domains of SLC3A2 from human, mouse, naked mole rat (NMR) and Neoaves reveals four distinct patterns.

To summarize, the deletion of SLC7A exchangers reduces the continuous consumption of ATP required to balance essential AA inside the cell. The loss of BCAT2 leaves BCAT1 to balance α-keto acids, the BCAA/BCKA ratio, and flux into the mitochondria. Perhaps this contributes to enhanced lipogenesis and adipose storage as required for prolonged flight, in contrast to mammals where greater catalysis of proteins occur with extended exertion (95) (**Fig. 9A**). This presumably requires other unexplored features of metabolic adaptation in Neoaves (**Fig. S14**). The exchange of α-keto acids at the plasma membrane by BSG*SLC16A transporters is Na^+^ independent and subject to positive regulation by the galectin lattice which appears to be reinforced by additional N-glycan branching (MGAT4B,MGATC duplications). Retaining muscle mass and fatty acid turnover is a critical feature of healthy aging and requires higher N-glycan branching as revealed by the analysis of Mgat5 deficient mice (22, 24). A lower cytoplasmic BCAA/BCKA ratio and mTOR activity in Neoaves may reduce ox-stresses associated with BCAA in mammals, and thereby support longevity.

#### Savings by preventative measures

In the control of ROS, gene duplication of the cystine transporter SLC7A11, may compensate for the loss of its adaptor SLC3A2. Moreover, SLC7A9 is also duplicated, a transporter that heterodimerizes with SLC3A1 (rBAT) and re-absorbs cystine in kidney tubules (107, 108). Unlike its paralogue, SLC3A1 is conserved in Neoaves and has six N-glycan sites but lacks the ubiquitination sites at the N-terminus. SLC3A1*SLC7A9 may cluster with AA/Na^+^ symporters as well as BSG- and ATP1B-associated transporters by galectin cross-linking with enhanced affinity due to MGAT4B and MGAT4C gene duplications (Fig. 9A). Notably, one of the MAGT4C paralogues (aka; Mgat6, Uniprot: Q9DGD1) evolved a novel specificity that adds an additional N-glycans branch not found in mammals (109). Thus, a stronger galectin lattice is expected to enhance cell surface expression of SLC3A1*SLC7A9 and growth factor receptors (22,24,74), thereby providing Cys to GSH and Coenzyme A (Ac-CoA) synthesis, linking ox-stress mitigation and TCA cycle in muscle, respectively (110) (Fig. 9A). Cys is also required in the biosynthesis of iron-sulfur clusters (Fe-S), prosthetic groups required by aconitase and succinate dehydrogenase, and three reductases of the electron transport chain.

Proton leakage in mammalian cells account for an additional ∼25% of energy consumption, in this case, energy lost to the ATP pool and increased ROS (102). The higher ATP/AMP ratio in SLC3A2 KO HeLa cells, and FLAG-SLC3A2 interactions with the mitochondrial ATP synthesis complex V suggest that Neoave energetics may benefit further from the loss of SLC3A2, by enhanced coupling of oxidative phosphorylation (**Fig. 2A, 6C,E**). Mitochondrial uncoupling proteins (UCP-1, 2) are deleted perhaps reducing ROS generation, while UCP3 and UCP4 are present (ie. SLC25A-7,8,9,27) (111).

#### Central metabolism and compartmentalized

Further manual analysis using UniProt and sequence alignment provided insight on additional streamlining of paralogues, that are often compartmentalized at the cellular and organ levels. Muscle glycogen synthesis (GYS1) is absent, presumably replaced by higher levels of circulating glucose (98) and by mobilization of fatty acids and ketones during flight. Carnitine O-palmitoyltransferase 1a (CPT1A, kidney, heart, liver) is present, and CPT1B/C (muscle, brain) are absent, consistent with liver as the major supplier of ketones to other tissues (**Fig. S14**). Mitochondrial pyruvate carboxylase (PC) is absent, an enzyme required to transfer citrate-derived oxaloacetate from the cytoplasm into the mitochondria TCA cycle, thus oxaloacetate is more likely redirected into gluconeogenesis, ox-PPP and HBP pathways. Moreover, phosphoenolpyruvate carboxykinase (PCK2) is also absent, leaving cytoplasmic PCK1 to mediate the reversible conversion of phosphoenolpyruvate and oxaloacetate in the cytosol, a key factor in flux of carbon between glucose, ox-PPP and lipid metabolism.

### Bat gene losses in common with birds

Bats are the only mammal to have evolved flapping-wing flight, which has also resulted in smaller genomes and extended life spans. However, Myotis lucifugus (Microbat) has SLC3A2, associated transporters and conserved N-glycan biosynthesis, but genes of interest in common with those purged in Neoaves, includes BCAT2, GCKR, FOXO1, FOXO6, PTEN, SREBF1, AKT1 and TP73 (AKT2 and TP53 in Neoaves). (**Fig. 8B, S12B**). These genes involved in signaling, transcription and stress mitigation, are intensely studied for their involvement in degenerate conditions of human aging. However, current “omics” type analysis, and CRISPR sgRNA drop-out screens in cultured cell lines are difficult to interpret as networks and reproduce in other conditions. Evolution leading to modern birds and bats has done a very successful gain-of-function gene drop-out experiment as evidenced by species radiation and longevity that includes an extended period of vitality in many species. Therefore, a detailed analysis of this large genomic space (>10,000 species over >100 Myr of evolution) has the potential to go well beyond studies on the genetic variation in dogs and other domesticated species (112, 113).

## DISCUSSION

Recent evolution of the N-glycan positions in mammalian SLC3A2 suggests these modifications are a major functional feature. The four NXS/T sites on human SLC3A2 display different N-glycan structures, indicating a sensitivity of *medial* and *trans-* Golgi remodeling enzymes to the microenvironment of each site. The primate-derived site at N381 and the conserved site at N424 are furthest from the membrane, and have predominantly tetraantennary N-glycans, which are higher affinity ligands for Gal3 binding. Mutational analysis indicated that N-glycans at N381 and to a lesser extent at N365, promote galectin binding and thereby tune SLC3A2 residency and endocytosis.

Restoring a lost site at A273N enhanced branching and poly-LacNAc on the neighboring N-glycan at N365, while restoring D355N suppressed poly-LacNAc at N365. Supplementing WT HeLa cell cultures with GlcNAc enhanced N-glycan branching and poly-LacNAc content only at N365, the site nearest the membrane most likely to interact with glycosylated AA/Na^+^ symporters and mediate endocytosis by the GL-Lect induced clathrin-independent carriers. Selection against sites at A273 and D355 in primates has removed their influence on N-glycan processing at N365, while retaining the capacity for regulation at N365 by metabolism and HBP.

### A model of N-glycan function in adaptors to transporters

SLC3A2 interacted with other glycoprotein adaptors including CD44, BSG and ATP1B1/3 which provide a spatial context to clustering of receptors and transporters by galectins. For example, membrane-anchored glycoconjugates of different sizes from glycolipids to the much larger receptor and proteoglycans, create vertical zones as recently described for integrin receptor clustering and tension with substratum (114). Clustering and tension are regulated by the Gal3 lattice, which facilitates β1-integrin dynamics within stable focal adhesions, thereby promoting signaling, turnover by GL-Lect endocytosis and cell motility (8,15,17). SLC3A2 N-glycans at N381 and N424 project above the membrane-embedded exchangers and interacts with the distal Gal3 lattice comprised of growth factor receptors and other glycoproteins (6, 74). These branched N-glycans with poly-LacNAc can extend further (>30Å) above the SLC3A2 fold reaching >100A, while N365 is within ∼10Å from the membrane (Fig. 5E). In AA/Na^+^ symporters of SLC7A (CATs), SLC1A and SLC38A families, the third extracellular peptide loop is also within ∼10Å of the membrane and has 1 to 3 conserved N-glycosylation sites where galectin cross-linking is possible. Indeed, N-glycosylation of sites in SLC7A1, SLC7A3 and SLC1A5 promotes stability and activity at the cell surface by interacting with galectins, (75,115,116). The AA/Na^+^ symporters have the potential to cluster with SLC3A2*SLC7A5 and regulate diffusion-limited exchange as suggested by the synchronized flux of BCAA and Glu when BSA was the only source of AAs (Fig. S3).

In the atomic structure, a cholesterol and phospholipid molecule were sequestered by the SLC3A2 TM domain in contact with SLC7A5 (58), suggesting an affinity for raft-like domains. Glycolipid-enriched domains also contribute to clustering and reduce the distance for diffusion between SLC3A2*SLC7A exchangers and the AA/Na^+^ symporters (Fig. 7D). Fluvastatin treatment reduces SLC3A2 N-glycosylation by OST in the ER which shifts its interactions toward CD44, SLC7A11 and a few AA/Na^+^ symporters that are required for GSH synthesis and stress mitigation (86, 87). Fluvastatin mimicked the SLC3A2 KO phenotype in HeLa cells, disrupting AA levels, glycolysis, HBP, nucleotides and sensitized to H_2_O_2_ (Fig. S10, S11). However, fluvastatin likely reduces N-glycosylation at sensitive sites in other stress related glycoproteins, notably c-AMP-dependent transcription factor (ATF6), cleavage-activating protein (SCAP) and activating endopeptidases (MBTPS1) (117–120). Statins also reduce mevalonate required for CoQ10 biosynthesis, which along with SLC3A2*SLC7A11, GSH synthesis and the oxidoreductase activity of FSP1, play a role in preventing lipid peroxidation and ferroptosis (121). Perhaps this is a cautionary note for people on long-term high-dose statins, as adverse effects due to inhibition of N-glycosylation and CoQ10 synthesis have been reported (83,122,123).

### Enhanced N-glycan branching in birds and metabolism

In mammals, N-glycan branching on glucose transporter 2 (SLC2A2) in β-cells promotes association with the galectin lattice, increases insulin secretion, and thereby surface expression of SLC2A4 in peripheral tissues which mediates glucose uptake (74, 124). The insulin and glucagon receptors (INSR, GCGR) stimulate competing pathways that are regulated at multiple levels (125). GCGR activity is highly dependent on N-glycan branching and binding to the galectin lattice (9, 74). Circulating glucose in birds is 2-3 fold higher than mammals, and in a complementary manner, duplication of MGAT4B and MGAT4C is expected to increase the ratio of GCGR to INSR signaling in liver, thereby gluconeogenesis and flux to ox-PPP and HBP. The paralogue of MGAT4C (ie. Mgat6, Uniprot: Q9DGD1) acquired a novel activity, adding a fifth N-glycan branch expected to increase affinities for galectins (11, 109). Neoaves have streamlined other genes associated with glucose homeostasis, notably losses of insulin-inducible glucose transporter (SLC2A4), glucokinase regulatory protein (GCKR), S/T-protein kinase (AKT2), fibroblast growth factor 21 (FGF21) (98), and an evolved resistance of proteins to glycation (126). Perhaps as a consequence, the Advanced Glycation End product-specific Receptor (AGER) is absent in Neoaves, which in mammals, sends AGE-damaged extracellular matrix proteins to intracellular degradation, as well as signaling that opposes ox-stress (127).

Higher NAD(P)H/NAD(P)+ ratios supporting high rates of oxidative phosphorylation and ROS mitigation may be required to drive oxidative phosphorylation in Neoave. In humans, an elevated cytosolic NADH/NAD+ (reductive stress) has been linked with common loss-of-function genetic variants in GCKR associated with ∼130 metabolic traits and diseases by GWAS, including elevated serum glucose, BCAA, α-hydroxybutyrate, triglyceride levels and insulin resistance (128). GCKR inhibits pancreatic and hepatic glucokinase (GK) during fasting and is rapidly reversed upon feeding, stimulating insulin secretion and glucose metabolism. In Neoaves, the stringency of switching from gluconeogenesis to glycolysis may be weakened by absence of GCKR and gain in N-glycan branching, perhaps ensuring sufficient glucose flux to ox-PPP under fasting conditions.

Loss of KEAP1 or amplification of NRF2 promotes tumor progression by suppressing the ox-stress associated with increased metabolic rates driven by oncogenesis (129). O-GlcNAcylation of KEAP1 activates binding to and ubiquitination of NRF2, linking nutrient sensing via HBP to redox stress signaling in cancer cells (130, 131). Deletion of KEAP1 disconnects NRF2 from this negative regulation, while preserving HBP regulation of the responses to stress through N-glycosylation. Notably, the stress-induced peptide:N-glycanase (NGLY1) serves a sensor of proteosome insufficiency by removing N-glycans from the NRF2 homolog, NRF1 (also known as NFE2L1) which upregulates SREBF2, thereby the mevalonate pathways and proteasome biogenesis (132–134). The mevalonate pathway is required for lipid-linked oligosaccharide synthesis and N-glycosylation as well as surveillance and repair of mitochondrial damage (135). In Neoaves, NRF1, NRF2, SCAP, MBTPS1/2 and SREBF2 are present; key genes for prevention and alleviation of proteotoxic stress, while SREBF1 is absent, a gene encoding a transcriptional regulator of BCKDK causally associated with elevated BCKA/BCAA in nonalcoholic steatohepatitis and obesity (136).

### Naked mole rats lack the ubiquitination sites in SLC3A2

If Neoave longevity is due, in part, to enhanced ROS suppression and dampened mTOR signaling, then perhaps the naked mole-rat accomplishes a similar feat (∼10 fold longer lifespan for a rodent of its size) without gene purging (101), by living underground with low oxygen consumption and metabolic rate. Naked mole-rat is resistant to cancer, shows negligible senescence, and maintain healthy vascular function longer into their lifespan than other rodents (137, 138). Extremely high-molecular-mass hyaluronan, a linear polymer of glucuronate β1-3/4GlcNAc (139), extremely error-free protein synthesis (140), increased expression of CDK inhibitors and genes mitigating ox-stress have been documented (141). The extracellular domain of SLC3A2 has five N-glycans including the primates-derived site at N381, which is expected to stabilize SLC3A2 in the galectin lattice. However, the ubiquitination sites are absent in a divergent cytoplasmic sequence (Fig. 9B). Together, the galectin lattice and GL-Lect mediated endosomal recycling may enhance SLC3A2 heterodimer stability, rather than loss to lysosomal proteolysis (73, 142). The naked mole-rat is a thermoconformer that thrives at lower body temperatures, while birds are 3-4°C warmer than mammals. The dynamics of interactions between SLC3A2*SLC7A exchangers and AA/Na^+^ symporters in the galectin lattice (ie. phase-transition condensate) are expected to increase exponentially as a function of temperature and distance (17, 143), which may in part, explain the evolution of SLC3A2 NXS/T and ubiquitination sites in these divergent vertebrate species. Finally, with apparent serendipity, SLC3A2 in hominids acts as a receptor for an envelope glycoprotein encoded by the endogenous retrovirus HML-2, which stimulates mTOR activity required in brain development and maintenance of brain stem cells (144). Overexpression of HML-2 is associated with tumorigenesis and neurodegeneration, similar in this regard to overexpressing of SLC3A2 in cancers more widely (45, 46).

In summary, our data suggests that N-glycans on primate SLC3A2 are positioned to promote surface expression and clustering of SLC3A2*SLC7A with N-glycosylated AA/Na^+^ symporters, and thereby regulate AA exchange, redox balance and mTOR/AMPK signaling. The evolved position of N-glycans on SLC3A2 determines Golgi remodeling, sensitivity to HPB and to lipid-linked oligosaccharide levels in the ER, which regulate Gal3 lattice retention at the cell surface and GL-Lect mediated endocytosis. Intrigued by the absence of SLC3A2 and many of its transporter partners in Neoaves, our preliminary analysis points to wider gene purging that rewiring energetics, while reducing burden of regulatory redundancies. The expansion of genes and their paralogues in early vertebrates may have maximized molecular interactions allowable by their bioenergetics, beyond which systemic instability increases (145, 146). Some genes and molecular interactions that were advantages in the past may now represent a health risk given the rapidly changing environment of our modern life. Comparative analysis of Neoave genomes and proteomes may reveal how natural selection has inadvertently extended their lifespans, providing clues to improve human health and aging.

## Acknowledgments

The authors thank Bret Larsen and Karen Colwill for assistance with SWATH and LC-MS/MS. Aldis Krizus for SLC3A2 expression vectors, Drs. Ran Kafri, Hakon Leffler and Kathryn Wellen for helpful discussions. J.W.D. acknowledges research support from Canadian Institutes of Health Research (CIHR) (MOP-62975). LJ gratefully acknowledges Mizutani Foundation for Glycosciences (reference no 200014), Agence National de la Recherche ANR (ANR-19-CE13-0001-01 and ANR-20-CE15-0009-01), and Fondation pour la Recherche Médicale (EQU202103012926). LJ also acknowledges the Cell and Tissue Imaging core facility (PICT IBiSA), Institut Curie, member of the French National Research Infrastructure France-BioImaging (ANR10-INBS-04).

## Author Contributions

J.W.D. designed the study, wrote the manuscript and prepared some figures. M.S.Z. and E.D. performed all endocytosis experiments. C.Z. performed and analyzed N-glycans and glycopeptide analysis as well as the SLC3A2 interactions by mass spectrometry. J.P performed the metabolite analysis by mass spectrometry as well as cell culture experiments. L.J. and M.S-Z. designed and performed the endocytosis experiments. Authors reviewed and approved the manuscript. D.W.J.N. assembled data on genes missing in birds and bats. A-C.G. L.P. and J.L. assisted with reagents and experimental advice on the fluvastatin experiments. G.G.H. designed the AA exchange using BSA.

## METHODS

### Materials

Antibodies to FLAG-M2 and tubulin were purchased from Sigma-Aldrich (St. Louis, MO). Metabolite standards and reagents were obtained from Sigma-Aldrich (St. Louis, MO) with minimal purity of 98%. ^13^C-uniformly labeled glucose and glutamine were purchased from Cambridge Isotope Laboratories,Inc. (Andover, MA). All organic solvents and water used in sample and LC–MS mobile phase preparation were HPLC grade and obtained from Fisher Scientific (Fair Lawn, NJ). N-Glycosidase F (PNGase F, EC 3.5.1.52, recombinant cloned from Flavobacterium meningosepticum and expressed in Escherichia coli) was purchased from Roche Diagnostics (Mannheim, Germany).

Antibodies for Western blots are rabbit anti-phospho-AMPK (T172) and AMPK with Cell Signaling #2531S and #2532S, respectively; rabbit anti-phospho-eIF2a Cell Signaling #9712S; rabbit anti-SLC3A2 (Santa Cruz Biotech. #361-375), CD98 (C-20) and sc-7095(Santa Cruz Biotech). Anti-human CD98 antibody, clone MEM-108, Biolegend #315602. Anti-EEA-1 antibody, BD Bioscience (#610456). Fluvastatin was purchased from US Biologicals (F5277-76). CRISPR/CAS9 Reagents: FastDigest BbsI (#FD1014) from Thermo Scientific; T7 DNA ligase (#M0318S), T4 PNK (#M0201S), ATP (#P0756S) from NEB; Plasmid-Safe™ ATP-Dependent DNase (#E3101K) from Epicentre; PolyJet™ reagent from SignaGen Laboratories; CRISPR-CAS9 vector pX459. Lactose, Sigma (#L3750), and Galectin-3 inhibitor compound from Galecto, Inc. (#GB0149-03), Home-produced Gal-3-488 recombinant protein. Genes lost and duplicated in Birds Protein sequences were for ∼250 Ave species were accessed through UniProt Taxonomy - Aves [8782]. The presence of genes in the 62 Ave species was also interrogated in OrthoDB which included Passeriformes, Galloanserae, Falconiformes, and Pelecaniformes and numbering 15,6, 5, 5 respectively, and 31 Neoaves. SLC3A2 is absent in Neoaves and present in truncated forms in Aves with only 21 of 250 aligned sequences showing a TM domain. Similarly, the KEAP1 gene is absent in Neoaves, and present in Galloanserae and some Passeriformes. Other Ave genes of interest were examined in the same manner and classified as absence if not found or severely truncated (>50%) in all or most Neoaves by UniProt Taxonomy; Aves [8782]. g:Profiler was used to compare GO and KEGG modules in Human, Serinus canaria (canary), and Myotis lucifugus (microbat).

### HeLa cells

HeLa Flp-In™ T-REx™ cells were a kind gift from Dr Stephen Taylor (University of Manchester). HeLa Flp-In T-REx cell lines were cultured at 37°C and 5% CO2 in a humidified atmosphere in high glucose DMEM (Wisent, St. Bruno, Quebec) supplemented with 10% fetal bovine serum (FBS), 2 mM Gln, penicillin/streptomycin, unless indicated otherwise. For HeLa cells with insertions of tetracycline-inducible FLAG-SLC3A2 WTseq and variants at the recombination site, expression was induced with the addition of 0.2 µg/ml doxycycline for 48-72 hours. This was followed by the addition of 0, 5, or 10 μM fluvastatin as indicated, and cells were cultured for a further 48 hours.

### CRISPR/Cas9 mutation of SLC3A2

Human SLC3A2 is a type II transmembrane glycoprotein with amino acids 1 – 184 cytoplasmic, 185 – 205 transmembrane and 206 – 630 extracellular. SLC3A2 has four isoforms; P08195-1 with the canonical sequence of 630 amino acids; P08195-2 is missing the first 102 residues; P08195-3 is missing 38-99; and P08195-4 has an insertion of 32 amino acids after position 98. Two guide RNA (sgRNA) sequences were made (https://zlab.bio/guide-design-resources) to target exon 4 and the flanking intron for removal of 110bp from the SLC3A2 gene. sgRNA#1: CAGATTCAACCGGAGGTACC, and sgRNA#2: CCGCGTTGTCGCGAGCTAC. The two sgRNA sequences were separately cloned into the CRISPR-CAS9 vector px459 (147). HeLa Flp-In T-REx cells were transfected with equal amounts of eachCRISPR-CAS9 px459 (sgRNA1and sgRNA2) plasmid using Lipofectamine 2000 according to manufacturer’s instructions. Following transfection, cells were cultured in DMEM (high glucose, 10% FBS) containing 1 μg/mL puromycin.

Colonies were expanded and analyzed by PCR and Western blotting. Genomic DNA was extracted using Genomic DNA Mini Kit (Geneaid). PCR products were then analyzed by agarose gel electrophoresis. Deletions were confirmed by PCR using Fwd: AGGAGGTGGAGCTGAATGAGT Rev: CGAGACTCAGAGGAGCTGATGT primers flanking the expected deletion:707bp without deletion, or 117bp with deletion. PCR deletion products were sequenced. Cells lysates in Tris pH 7.5 50mM and 2% SDS were applied to SDS-PAGE followed by Western blotting using anti-SLC3A2 (Santa Cruz, 1:5000) and anti-ϒ-tubulin antibodies (1:10,000). The transfer membranes were then probed with HRP-conjugated secondary antibodies.

### Real-time reverse transcriptase quantitative polymerase chain reaction (RT-qPCR)

RNA from cell pellets was extracted using Qiagen RNeasy kit, quantified using Nanodrop, and reverse transcribed with Superscript II. PCR was performed using the ABI Prism 7900HT Sequence Detection System (Applied Biosystems). Amplification mixture contained 1 µL cDNA and 10 µL SYBR Green PCR Master Mix (4364344; Invitrogen). Each reaction was performed in triplicate. Quantification data generated by PCR software SDS2.2.2 (Applied Biosystems) was analyzed using the ΔΔCt analysis method against GAPDH expression (148). Primers used were as follows:

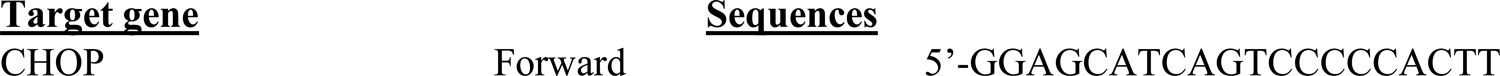
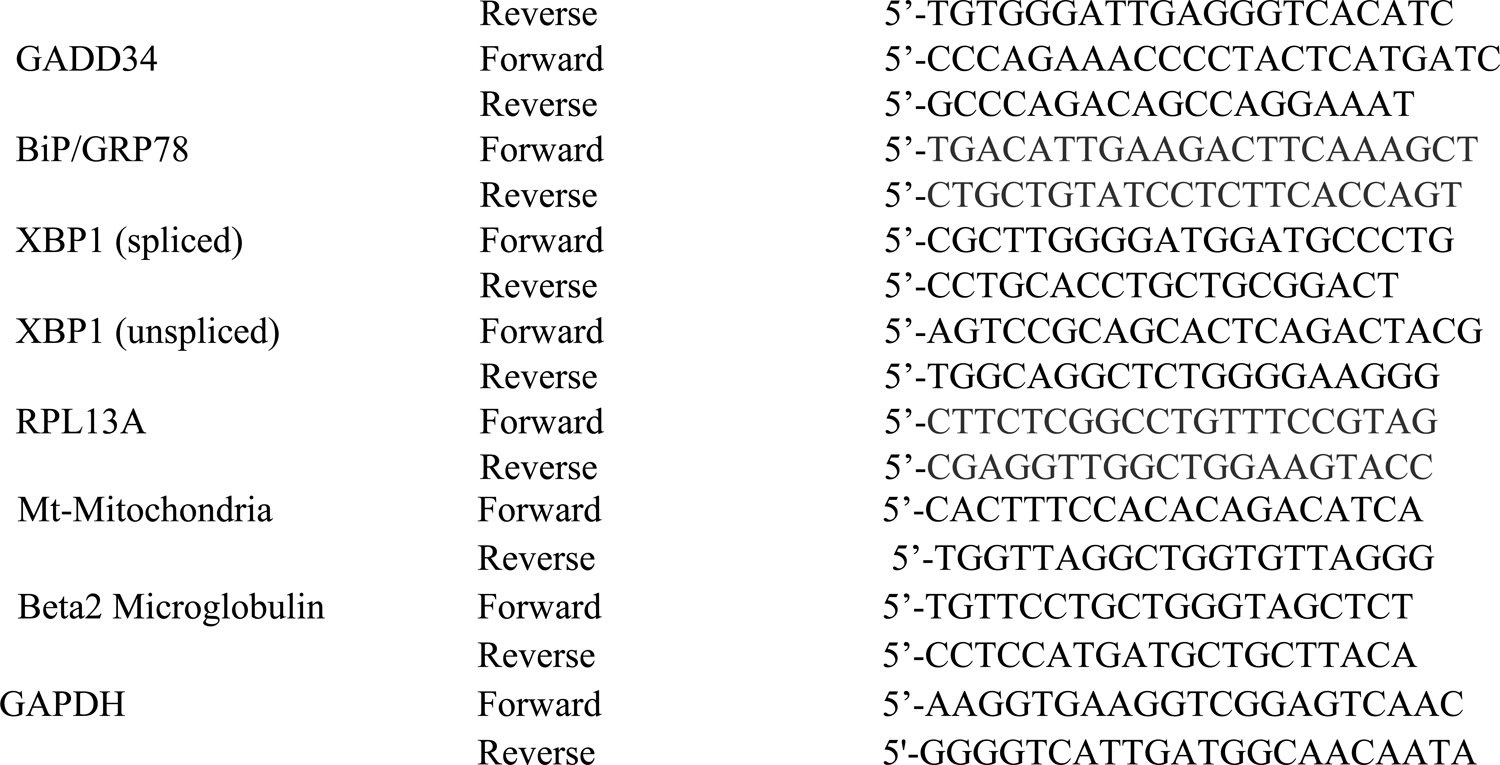

### Re-expression of SLC2A3 wild type and glycosylation site variants

SLC3A2 has 4 isoforms that differ in first ∼100 amino acids at the cytosolic N-terminus, and HeLa cells express the multiple isoforms. The N-terminal sequence is dispensable for interactions with SLC7A5 as revealed by X-ray crystallography and cryo-electron microscopy (58). To limit experimental variation in N-glycan profiling and AP-MS/MS analysis, we made the NXS/T(X≠ P) site variants in one FLAG-tagged isoform in the HeLa Flp-In T-REx SLC3A2 KO cells cultured under standard high glucose culture conditions. Human SLC3A2 cDNA encoding amino acids 66-630 was FLAG-tagged at the N-terminus and cloned into the pcDNA5/FRT/TO expression vector. N-glycosylation site mutations were introduced by a combination of fusion PCR and restriction enzyme cloning using the NEB Q5® Site-Directed Mutagenesis Kit. All mutations (site deletion at N365D and N381D and site additions at A273N, D355N, D492N) were confirmed by DNA sequencing. Each construct was cloned into the pcDNA5/FRT/TO plasmid and integrated into the genome of the HeLa Flp-In T-REx SLC3A2 KO cells described above, at a pre-integrated FRT recombination site, by co-transfection with Flp recombinase encoding POG44 plasmid, using PolyJet™ reagent (SignaGen Laboratories) and DMEM media without FBS. Following selection in 200 µg/mL of hygromycin, resistant clones were tested for inducible FLAG-SLC3A2 expression following 48h of culture with 0.2 µg/mL doxycycline (dox). Cell lysates were analyzed by Western-blotting using polyclonal anti-CD98 (C-20).

### Metabolite profiling by LC-MS/MS (Metabolomics)

Cells were seeded in 6-well plates with 6 technical replicates per experimental condition and cultured for 24 h, followed by two quick washes of the wells with warm PBS (∼37°C), then the plates were flash frozen in liquid nitrogen (149). The metabolites were immediately extracted by adding 1 ml of extraction solution (40% acetonitrile, 40% methanol, and 20% water) and then the cells were scraped and collected in 1.5 mL vials. The mixture was shaken for 1 hour at 4°C and 1400 rpm in a Thermomixer (Eppendorf, Germany). The samples were spun down at 14000 rpm, for 10 minutes at 4°C (Eppendorf, Germany), and supernatants were transferred into fresh tubes to be evaporated to dryness in a CentreVap concentrator at 40°C (Labconco, MO). The dry extract samples were stored at −80°C until LC-MS/MS analysis. Following reconstitution in 100 µL of water containing internal standards (500µg/ml and 300µg/ml of D7-Glucose and ^13^C9^15^N-Tyrosine, respectively), samples were injected by auto-sampler (Dionex Corporation, CA) onto an HPLC column in line with a triple-quadrupole mass spectrometer (AB Sciex 5500Qtrap, Toronto, ON, Canada). Metabolites were separated through a guard column (Inertsil ODS-3, 4 mm internal diameter x 10 mm length, 3 μM particle size) and analytical column (Inertsil ODS-3, 4.6 mm internal diameter, 150 mm length, and 3-μM particle size; GL Sciences, Japan) for both polarity modes. In positive mode analysis, the organic portion (acetonitrile) of the mobile phase (0.1% acetic acid) ramped from 5% to 90% in 16 minutes, held for 1 minute at 90%, then returned within 1 minute to 5% acetonitrile in mobile phase for column regeneration. In negative mode, the acetonitrile composition ramped from 5 to 90% in 10 minutes, held for 1 min at 90%, then the gradient ramped back over 3 minutes to 5% acetonitrile in mobile phase (0.1% tributylamine, 0.03% acetic acid, 10% methanol), to regenerate the column for the next run. The total runtime for each sample in both modes was 20 minutes and the samples were maintained at 4°C in the auto-sampler, and the injection volume was 10 μL. An automated washing procedure was developed before and after each sample to avoid sample carryover. The mass spectrometric data acquisition time for each run was 20 minutes, and the dwell time for each MRM channel was 10 milliseconds. Settings for negative and positive modes with electrospray ionization (ESI) were optimized. MultiQuant software (AB Sciex, Version 2.1) was used for peak analysis of ∼120 targeted metabolites, with standards run consecutively. Peaks representing targeted masses and LC retention times were confirmed manually. Common mass spectrometric parameters: GS1 and GS2 were 50 psi; CUR was 20 psi, and CAD was 3 and 7 for positive and negative modes, respectively, and source temperature (TEM) was 400°C. Signal was normalized to internal standard and cell number. MetaboAnalyst was used for preliminary statistical analysis of the LC-MS/MS data (https://www.metaboanalyst.ca/MetaboAnalyst/ModuleView.xhtml)(150). The LC-MS/MS system does not resolve hexose and hexosamine isomers including glucose/galactose and GlcNAc/GalNAc. To monitor trends in metabolic pathways, we refer to these isomers in their glucose (Glc) forms.

### Cell migration by scratch-wound assay

Cells were seeded into 6-well plates and serum starved for 24 hours, as described. A P10 pipette tip was used to create a linear wound, and healing by cell migration was monitored for 24 hours using time-lapse microscopy via an inverted microscope (DMIRE2; Leica Microsystems) with a motorized stage and live-cell apparatus (37 °C humidified chamber with 5% CO2; Applied Scientific Instrumentation, Eugene, OR, USA). Images were captured every 15 min using an ORCA Hamamatsu CCD camera (Hamamatsu Corporation, Bridgewater, NJ, USA) with a × 10 lens. Image analysis was carried out by Velocity 3D Image Analysis Software (Improvision) by measuring the total wound area in three fields per condition, at t = 0, 4, 6, 8,12 and 24 hours. Measurements at each location were averaged to yield a mean wound area. The residual wound area was expressed as a percent of original wound.

### Cell growth

Cell lines were plated in 96-well Nunc 165305 Optical-Bottom plates at 1000 cells/well and growth was measured daily as a fraction of well confluence using the Celigo Cell Cytometer’s label-free bright field imaging. Scans were repeated daily until confluence reached approximately 100% and data graphed as a fraction of confluence with time. Cell growth was also analyzed using Alamar Blue reagent (Invitrogen), and plate reader (Gemini Fluorescence from Molecular Devices) at fluorescent excitation 590 nm and emission at 544 nm.

The induced response to ox-stress was also measured with Alamar Blue, which reacts with reducing-potential (i.e. NAD(P)H). HeLa cells with insertions of tetracycline-inducible FLAG-SLC3A2 at the recombination site, were induced by the addition of 0.2 µg/ml doxycycline for 48 hours, after which they were plated at 7000 cells per well in 96 well optical plates in rows of 12 replicates. Control wells of non-Tet induced cells were usually included in each assay. After 24 hours in the 96 well plates, freshly diluted H_2_O_2_ was titrated at 0 to 600 uM across the 12 replicate wells and incubated for 16h. Alamar Blue and Hoescht were added to each well and the Alamar Blue signal was read as above after 1 and 4 hours and normalized to cell number by Hoescht staining and counting of nuclei by InCell imaging.

### Export and recovery of amino acid from the medium in restricted conditions

Media samples for metabolomics analysis were prepared from 1.2 x 10^6^ cells plated in 6 well plates, allowed to adhere in complete media for 12 hours. Cells were briefly rinsed with PBS (with Ca2+/ Mg2+) and then incubated with AA-free DMEM/F12 media (US Biological) for 3 hours. Fresh amino acid-free DMEM/F12 supplemented as indicated with BSA or BSA+ Gln and then media aliquots were taken at the indicated time points. Aliquots were centrifuged at 16100 g for 5 minutes, transferred to fresh tubes, and stored at −80°C until further processing. For targeted metabolomics by LC-MS/MS, 10 μL media aliquots (thawed just until melted) were extracted with 500 μL ice cold extraction solvent (40% acetonitrile:40% methanol:20% H_2_O) and vortexed for 1 minute, then shaken in an Eppendorf shaker (Thermomixer R) at 1400 rpm, 4°C for 20 minutes and centrifuged at 4°C for 15 minutes at 16100 g. Supernatants were transferred to a clean Eppendorf tube, evaporated to dryness in a CentreVap concentrator at 40°C (Labconco, MO) and stored at −80°C, then resuspended in 180 μL of water containing the Internal Standards and analyzed by LC-MS/MS as described above.

### Antibody binding and uptake experiment

#### Binding

Cells were shifted on ice for 10 min on ice (4 °C) and washed 3 times with ice-cold DMEM. 10 μg/ml of anti-SLC3A2 (MEM-108 clone) prepared in the same medium was added to the cells for 30 min incubation. Excess of antibody was removed by 3 successive washes with ice-cold DMEM medium. Cells are then PFA-4% fixed and immuno-stained against the bound antibody. In the case of co-binding assay (i.e anti-SLC3A2 and Gal3), 5 μg/ml of recombinant Gal3-488 was primarily bound on the cooled cells for 30 min prior to switch to the antibody binding step. Three ice-cold DMEM washes are required between Gal3 binding and antibody binding steps. **Uptake**: continuous antibody uptake was performed by incubating cells with 10 μg/ml of anti-SLC3A2 (MEM-108 clone) for 10 min at 37 °C. Cells were then shifted to 4 °C on ice, 3 times washed with ice-cold-PBS, then PFA-4% fixed, saponin-permeabilized and immuno-stained against the uptaken antibody. For co-uptake experiment, Gal3 and anti-SLC3A2 were sequentially bound on pre-cooled cells, as described above, and then shifted to 37 °C for 10 min internalization.

#### SLC3A2/Gal3 interaction

Experiment was performed as described above for co-binding experiment except that here the cells were lysed. PNS-cleared lysate was then overnight incubated at 4°C with protein G-sepharose beads for SLC3A2 immuno-precipitation (IP). Samples were denatured and eluted from the beads for SDS-PAGE analysis. Co-pulled down Gal3 was further quantified.

#### Gal3 inhibition using GB0149-03 or lactose. Acute treatment

Cells were pre-treated with 10 μM GB0149-03 compound or 150 mM lactose for 5-10 min at 37 °C. The inhibitors were washed out during the following antibody uptake experiment. **Severe treatment**: Cells were pre-incubated with 10 μM GB0149-03 compound or 150 mM lactose for 30 min and 60 min at 37 °C. In contrast with the acute treatment, here the inhibitors were kept during the internalization step.

### Extract of endogenous SLC3A2 glycopeptides from membrane proteins

HeLa cells (∼2 x 10^6^) were suspended in 1 mL of homogenization buffer (0.25 M sucrose, 50 mM HEPES pH 7.5, 5 mM NaF, 5 mM EDTA, 2 mM DTT, cOmplete protease inhibitor), and lysed using a probe sonicator. Homogenate was cleared at 2,000 g for 20 min at 4°C, then ultracentrifuged at 115,000 g for 70 min at 4°C. The pellet was vigorously suspended in 650 μL Tris buffer (0.8% Triton X-114, 50 mM Tris pH 7.5, 0.1 mM NaCl, 5 mM EDTA, 5 mM NaF, 2 mM DTT, cOmplete protease inhibitor). The homogenate was chilled on ice for 10 minutes, incubated at 37°C for 20 minutes, then phase partitioned at 1,950 g for 2 minutes at room temperature. The upper phase was discarded. Membrane proteins in the lower phase were precipitated with 1 mL acetone at −20°C overnight. After centrifugation at 1950 g for 2 minutes, the precipitated membrane proteins were stored at −25°C.

### SDS gel separation of endogenous SLC3A2 and in-gel digest

SDS-PAGE gel loading buffer (250 uL) was added to extracted membrane pellets from HeLa cells, agitated and heated 2 minutes at 100°C to denature proteins. Proteins were separated on 8% SDS-PAGE gels, stained with GelCode Blue, and bands between 80 to 90Da (band 1), 90Da to 110 Da (band 2) and 110 Da to 130 Da (band 3) were cut. The gel pieces were washed with 50 mM ammonium bicarbonate (ABC) and acetonitrile (ACN) 4 times, without reduction and alkylation, followed by in-gel digest with 25 ng/µL trypsin in 50 mM ABC overnight at 37°C. Tryptic peptides were extracted from the gel pieces with 25 mM ABC and 0.5% FA, then dried by speedvac. Tryptic peptides were dissolved in 20 µL 0.5% FA and de-sialidated with sialidase and processed for glycopeptide identification and site specific N-glycan analyses.

### Glycopeptides Enrichment

HILIC microtips were prepared with 10 mg of PolyHYDROXYETHYL A (PolyLC Inc. Columbia MD), in a bed volume of 50 μL, washed with 500 μL of ddH_2_O and equilibrated with 500 μL HILIC solvent (80% ACN, 0.1% TFA). Dried tryptic peptides were dissolved in 100 μL HILIC solvent, loaded slowly onto microtips and spun down at 700-1000 rpm. Sample was loaded a second time into the microtip then washed 3 times with 1 ml HILIC solvent, and glycopeptides eluted with 500 μL of 100 mM ammonium bicarbonate. The eluted glycopeptides were speed vacuumed to dry, and analyzed by LC-MS and MS/MS.

### Glycopeptide preparation for dox-induced FLAG-SLC3A2

Dox-induced FLAG-SLC3A2 variant cell lines (∼10^7^) were lysed in 1 mL of lysis solution (1% Triton100, 20 mM Tris pH7.5, 150 mM NaCl, 1 mM EDTA and 1mM EGTA, with freshly prepared protease inhibitors), and centrifuged at 14,000 rpm for 30 minutes to pellet the nuclei and insoluble material. A 15 μL aliquot was suspended in SDS loading buffer, heated at 100°C for 1 minute and followed by SDS-PAGE and Western blotting to confirm induced expression of FLAG-SLC3A2.

Protein concentration was determined by Thermo BCA protein assay. Normalized protein 1mg lysates were incubated with anti-FLAG M2 antibody-conjugated agarose 20 μL (bed volume, from 40 uL FLAG beads slurry (1:1)) overnight on a rotating platform at 4°C°. The beads were washed 4 times (1 mL) with TBS (50 mM Tris pH7.5, 150 mM NaCl) and 3 times with 50 mM ammonium bicarbonate (ABC, 0.8 mL) before re-suspending in 20 mL ABC (50 mM). For on-beads digest, trypsin (enzyme ∼0.5µg) was directly added to the beads and incubated at 37°C overnight. Next morning, 0.25 ug trypsin was added again and digested for another 2 hours, then heated at 100°C for 1 minute to inactivate trypsin. Peptide was extracted with 50 μL 0.5% formic acid 3 times and dried by speedvac. Samples were dissolved with 10 μL 50 mM ABC with 0.5 μL sialidase and incubated at 37°C overnight. Half of trypsin digest was used for in-solution Asp-N digest (Asp-N (120 ng, 40µg/µL), 37°C overnight digest.) Both trypsin and trypsin plus Asp-N digest samples were injected (0.5 μL) for LC-MS/MS glycopeptide identification and glycopeptide quantification by LC-MS.

### Glycopeptide analysis by LC-MS and MS/MS

Tryptic peptides dissolved in 100 μL HILIC solvent were prepared on HILIC microtips. Samples were then applied to a nano-HPLC Chip using an Agilent 1200 series microwell-plate autosampler, interfaced with an Agilent 6550 iFunnel Q-TOF MS (Agilent Inc., Santa Clara, CA). The reverse-phase nano-HPLC Chip (G4240-62002) had a 40 nL enrichment column and a 75 μm x 150 mm separation column packed with 5 μm Zorbax 300SB-C18. HILIC retains hydrophilic glycopeptides. For HILIC-enriched hydrophilic glycopeptide, mobile phase began with 2% ACN. For hydrophobic glycopeptides, mobile phase began with 8% ACN. The mobile phase was 0.1% formic acid in water (v/v) as solvent A, and 0.1% formic acid in ACN (v/v) as solvent B. The flow rate at 0.3 μL/min with gradient schedule; 2% B (or 8% B) (0−0.2 minute); 2−40% B (or 8-40%) (0.2−35 minute); 40−80% B(35−42 minute); 80% B (42−45 minute) and 80-2% B (45-50 minute). The MS system was in positive ion mode with 2 GHz Extended Dynamic Range mode: V^cap^: 1,800–1,900 V; drying gas flow 5.0 L/minute at T = 280 °C; fragmentor voltage 360 V: precursor selection 10 precursors/cycle; threshold 1,000 counts abs and 0.001% rel; active exclusion after 2 spectrum; and release after 0.5 minutes to start again. Internal reference mass calibration used m/z 1221.9906 (Agilent), 445.1200 and 741.1951 (Polysiloxane). LC-MS/MS targeting specific glycopeptides was done to confirm the glycosylation site and glycan structure. For glycopeptide quantification, run in MS mode only, mass range was set to standard (3200 *m/z*).

**Protein identification:** was done with mass range set to narrow (Low 1,700 *m/z*); auto MS/MS at 8 MS (range 350–1700 *m/z*), 3 MS/MS (range 50–1,700 *m/z*) per/s; narrow isolation width (1.3 *m/z*), with collision energy determined on the fly using a slope of 3.6 and intercept of –4.8

**Glycopeptide MS/MS:** analysis of N-glycan structures used mass range set to standard (3200 *m/z*); auto MS at 8 MS (range 700–2500 *m/z*), MS/MS (range 100–3000 *m/z*) per/s; narrow isolation width (1.3 *m/z*), with collision energy determined on the fly using a slope of 1.8 and intercept of –4.8.

**Targeted glycopeptide analysis:** was done with mass range set to standard (3200 *m/z*); targeted MS/MS was set to different CE for inclusion of targeted glycopeptides.

**Data analysis:** Mascot search was used to identify proteins and peptide sequences. Glycopeptide were identified by the presence of hexose and N-acetylhexosamine using Agilent Masshunter Quantitative Analysis software (B06.01) and glycan structures were predicted for extracted glycopeptides by online GlycoMod (http://web.expasy.org/glycomod/) and confirmed manually.

### Total cellular N-glycan profiling

Precipitated membrane protein (30 μg) was suspended in 60 μL of 0.25% RapiGest SF, 50 mM ammonium bicarbonate, 5 mM DTT, heated at 85°C for 3 minutes, then mixed with 1 μL of PNGase F, 0.7 μL of sialidase, and 20 μL of 50 mM ammonium bicarbonate, and incubated at 42°C for 2 hours followed by 37°C overnight. Released *N*-glycans were extracted with 4-5 volumes of 100% ethanol at −80°C for 2 hours. The supernatant containing released *N-*glycans was speed vacuumed to dry. Pipet tips packed with 10 mg porous graphitized carbon (PGC) for a bed volume of 50 μL were washed with 500 μL of ddH_2_O, 500 μL of 80% acetonitrile (ACN), and equilibrated with 500 μL 0.1% trifluoroacetic acid (TFA). *N-*glycan pellets were dissolved in 50 μL of 0.1% TFA and slowly loaded into the microtips at a flow rate of ∼100 μL/min, washed with 500 μL 0.1% TFA, and *N-*glycans eluted with 500 μL of elution buffer (0.05% TFA, 40% ACN). The eluted *N-*glycans were analyzed by LC-MS/MS. Total N-glycan samples were applied to a nano-HPLC Chip using an Agilent 1260 series microwell-plate autosampler, and interfaced with an Agilent 6550 iFunnel Q-TOF MS (Agilent Technologies, Inc., Santa Clara, CA). The HPLC Chip (glycan Chip) had a 40 nL enrichment column and a 75 μm x 43 mm separation column packed with 5 μm graphitized carbon as the stationary phase. The mobile phase was 0.1% formic acid in water (v/v) as solvent A, and 0.1% formic acid in ACN (v/v) as solvent B. The flow rate at 0.3 μL/minute with gradient schedule; 5% B (0−1 minute); 5−20% B (1−15 minute); 20−70% B(15−16 minute); 70% B (16−19 minute) and 70-5% B (19-20 minute). The MS System was operated in positive ion mode at 2 GHz Extended Dynamic Range, MS mode in low mass range (1700m/z) with MS setting at 8 MS (range 450–1700 *m/z*). **Data analysis:** PNGase F released free N-glycan was identified by Agilent Masshunter Quantitative Analysis software by the presence of hexose and N-acetylhexosamine. N-glycan structure were predict by online GlycoMod (http://web.expasy.org/glycomod/). Finally, Agilent Masshunter Quantitative Analysis software was used to quantify the extracted glycan peaks.

### SLC3A2 interactors by AP-MS

HeLa Flp-In T-REx cells with an insertion of dox-inducible FLAG-SLC3A2 were seeded (2 x 10^6^ cells per 10 cm plate (Nunc) in DMEM high glucose media plus 10% FBS, followed 24 hours later by the addition of 0.2 µg/mL doxycycline for 48 hours. Cells were pelleted and stored at −80°C. Cell were lysed in 250 µL FLAG lysis buffer (0.5% NP40, 50 mM HEPES pH 8.0, 100 mM KCl, 1 mM EDTA, 10% glycerol with fresh protease inhibitors) as described (151). Samples were freeze and thawed once then agitated in 37°C water bath to thaw, and centrifuged at 14,000 rpm for 30 minutes to pellet the nuclei and insoluble material. An aliquot of cell lysate was suspended in SDS loading buffer, heated at 100°C for 1 minute and followed by PAGE and Western blotting with anti-SLC3A2 antibody to check the SLC3A2 expression (protein concentration was determined by Bradford analysis). For proteomics, 1 mg of protein in ∼240 µL lysate was incubated with 40 µL anti-FLAG M2 antibody-conjugated agarose slurry (1:1) for 4 hours on a rotating platform at 4°C. The beads were washed 2 times (1 mL) with FLAG lysis buffer, 2 times with FLAG Rinsing Buffer (20 mM Tris pH 8.0 and 2 mM CaCl_2_) and 2 times with 50 mM ammonium bicarbonate (ABC, 0.8ml) before re-suspending in 20 mL ABC (50 mM) for on-beads trypsin digest.

### Data dependent acquisition (DDA) and SWATH

Samples were prepared for protein identification by DAA and quantification by SWATH using TripleTOF Mass Spectrometer as described (152). Briefly, each sample (5 µL) was directly loaded onto an equilibrated HPLC column at 400 nL/minute flow rate. The peptides were eluted from the column over a 90 minute gradient generated by a NanoLC-Ultra 1D plus (Eksigent, Dublin CA) nano-pump and analyzed on a TripleTOF 600 instrument (AB SCIEX, Concord, Ontario, Canada). The gradient was delivered at 200 nL/minute starting from 2% acetonitrile with 0.1% formic acid to 35% acetonitrile with 0.1% formic acid over 90 minute followed by a 15 minute cleanup at 80% acetonitrile with 0.1% formic acid, and a 15 minute equilibration period back to 2% acetonitrile with 0.1% formic acid, for a total of 120 minute. To minimize carryover between each sample, the analytical column was washed for 3 hours by running an alternating sawtooth gradient from 35% acetonitrile with 0.1% formic acid to 80% acetonitrile with 0.1% formic acid, holding each gradient concentration for 5 minutes. Analytical column and instrument performance were verified after each sample by loading 30 fmol bovine serum albumin (BSA) tryptic peptide standard (Michrom Bioresources Fremont, CA) with 60 fmol a-casein tryptic digest and running a short 30 minute gradient. TOF MS calibration was performed on BSA reference ions before running the next sample to adjust for mass drift and verify peak intensity. The TripleTOF 600 method was set to data dependent acquisition (DDA) mode, which consisted of one 250 ms (ms) MS1 TOF survey scan from 400–1300 Da followed by 20 100 ms MS2 candidate ion scans from 100–2000 Da in high sensitivity mode. Only ions with a charge of 2+ to 4+ that exceeded a threshold of 200 cps were selected for MS2, and former precursors were excluded for 10 seconds after one occurrence. Half of the sample was analyzed by DDA as above, and the other half was analyzed as described below by data independent acquisition (SWATH).

For SWATH, acquisition consisted of one 50 ms MS1 scan followed by 54 dynamic isolation windows covering the mass range of 400–1250 a.m.u. (cycle time of 3.25 s); an overlap of 1 Da between SWATH was preselected. The collision energy for each window was set independently as defined by CE = 0.06 3 m/z + 4, where m/z is the center of each window, with a spread of 15 eV performed linearly across the accumulation time.

### Protein identification

Mass spectrometry data were stored, searched, and analyzed using the ProHits laboratory information management system (LIMS) platform (153). DDA data files were searched using Mascot against human protein database with the RefSeq database (version 57, NCBI) against a total of 72,482 human and adenovirus sequences supplemented with common contaminants from the Max Planck Institute (http://141.61.102.106:8080/share.cgi?ssid=0f2gfuB) and the Global Proteome Machine (GPM; https://www.thegpm.org/crap/index.html). Database parameters were set to search for tryptic cleavages, allowing up to two missed cleavage sites per peptide with a mass tolerance of 40 ppm for precursors with charges of +2 to +4 and a tolerance of ± 0.15 a.m.u. for fragment ions. Deaminated asparagine and glutamine and oxidized methionine were allowed as variable modifications.

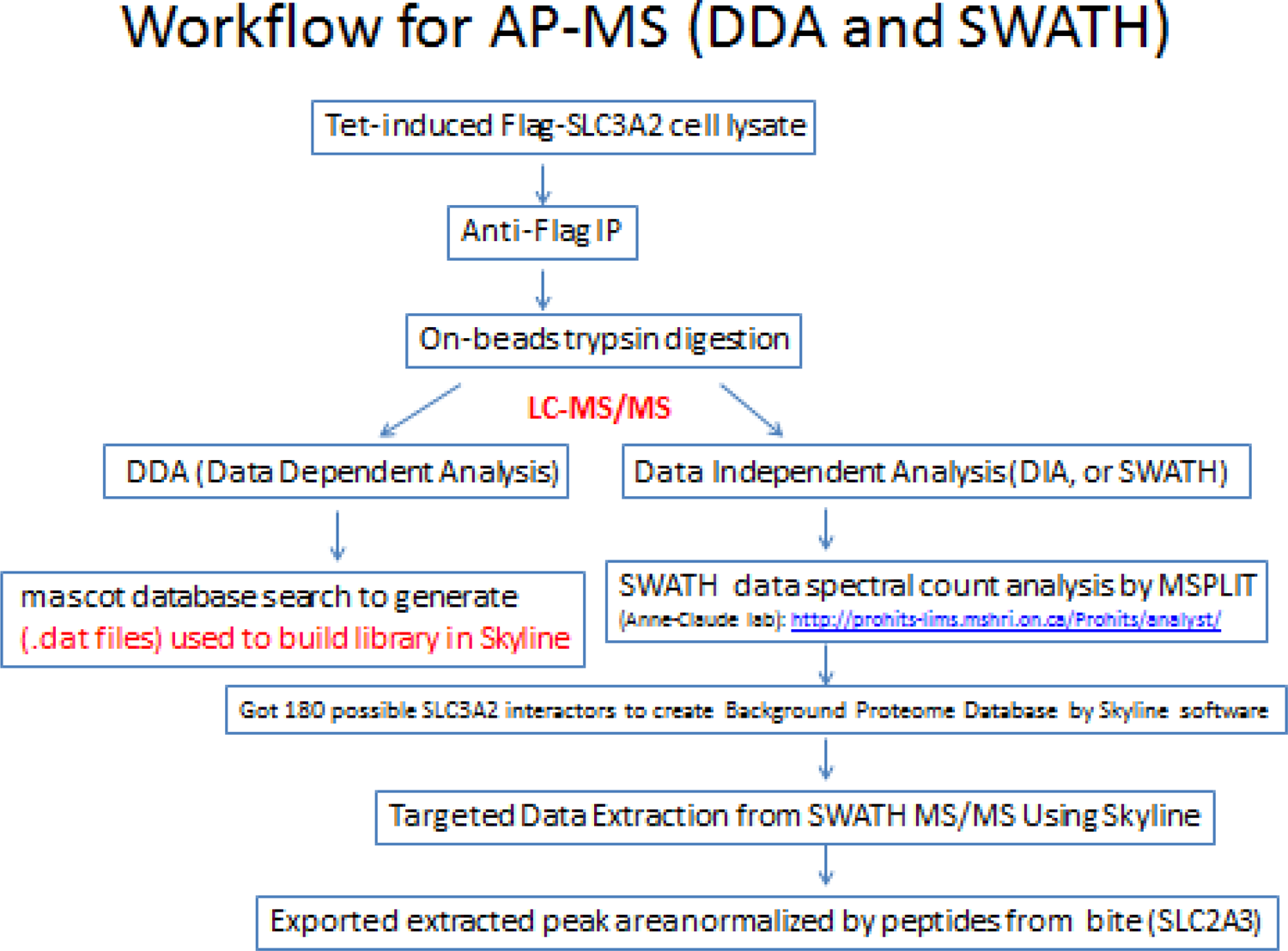

SWATH data file was analyzed with MSPLIT in ProHits. Library was generated in-house using DDA files searched with MS-GFDB for trypsin digest, revealing all cleavage sites with mass tolerance of 50 ppm for parent ions (http://proteomics.ucsd.edu/Software/MSGFDB/) and set as DDA mascot search. Only oxidized methionine was allowed as a variable modification. MSPLIT-DIA parameter settings were: FDA, 0.01; fragment mass tolerance, 50 ppm; retention time window 10 minutes; variable SWATH window from 400 to 1250 (Table S6). After the MSPLIT analysis of SWATH data, spectral counts with 3 or more unique peptides detected per protein in dox-induced SLC3A2 (255I WT) and near absent in SLC3A2 KO cell samples were used to generate a preliminary list of 156 SLC3A2 interactors. Running MSPLIT-DIA prior to targeted extraction by Skyline restricted the search space by providing accurate retention times and a list of the peptides expected to be in the sample.

### Targeted Data Extraction from SWATH MS/MS Using Skyline

Skyline (http://proteome.gs.washington.edu/software/skyline) is an open source, Windows-based software for curating and analyzing data from proteomic experiments. A primary list of 185 sequences of interest were retrieved from Uniprot and uploaded onto Skyline to create a background Proteome Database. Detailed Skyline and data processing settings are given in (154). Briefly, the spectral library generated from DDA files was uploaded in Skyline, and SWATH-MS data files were processed using the full scan MS/MS filtering at a resolving power of 30,000. Unique peptides were refined and curated for reproducible fragment ions. Peak boundaries for each selected peptide were manually supervised and when necessary, adjusted. The reproducibility and reliability of selected peptides and transitions were verified visually by looking at the ion peak to area ratio across the samples. We used at least 3 peptides per protein and at least three fragment ions per peptide for every protein. The extracted transition peak areas were exported to excel as raw intensities and normalized to the intensity of SLC3A2 tryptic peptides. The total intensity of representative peptides for each protein was summed, then divided by the number of transitions, and the resulting average intensity was used to draw comparisons for SLC3A2 interactors (for the same gene, same parent ion, same fragment). The normalized intensity was then converted to a percentage (relative abundance), using the highest value each experiment. The relative abundance of transitions representing the same peptides for each protein were averaged and presented as protein level and used to compare fold change for SLC3A2 interactors across all mutants and replicas. Data was normalized to the highest XIC (extracted ion chromatogram), as a reference to the maximum value across all samples. The Kruskal Wallis test was used to compare data for wild type and all mutations to determine significant differences, then the Dunn test was applied for pairwise comparison eliminating two that were not different. p <=0.05

---

**Figure S1.**
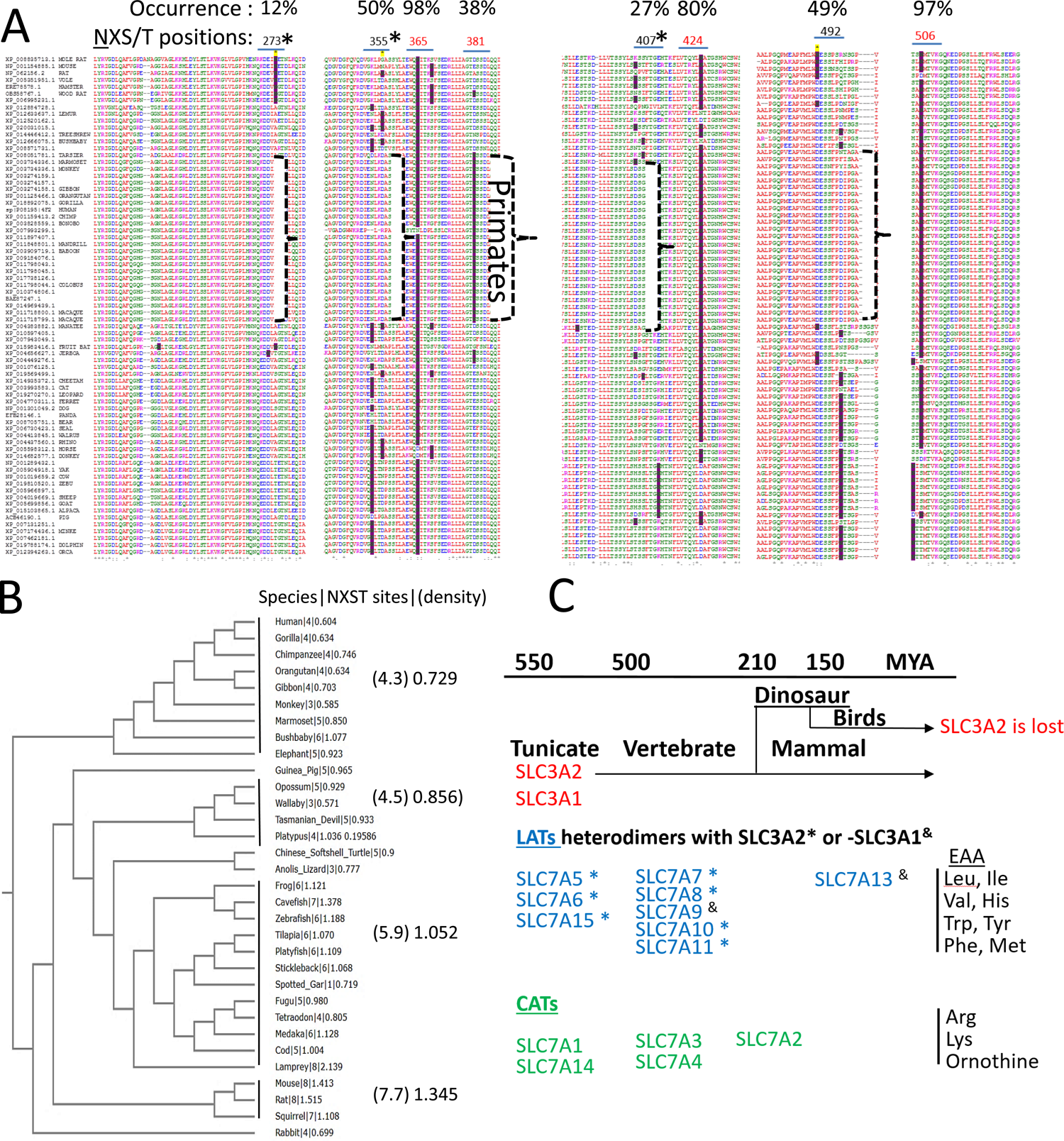
**(A)** Alignment of mammalian SLC3A2 sequences with the N of NXS/T(X≠P) marked (dark red fill) to reveal recent evolution of N-glycosylation sites. Variation in site position are observed in 3-6 AA regions, and four sites have been lost in the human sequence. NXS/T positions in primate are N365, N381, N424, N492. * indicates lost ancestral sites which were added experimentally to human SLC3A2 and expressed in HeLa cells. **(B)** An unrooted phylogenetic tree of vertebrate SLC3A2, indicating the NXS/T sites (number) and density (sites/100 AA) on the right. **(C)** SLC7A family of sodium-independent transporters. Heterodimers with SLC3A2 * and SLC3A1 ^&^. SLC3A2, SLC3A1 and SLC7A1-4, 14 (CATs) are N-glycosylation, but the LATs are not. On the right: an incomplete guide to AA **s**ubstrates for LATs and CATs.

**Figure S2.**
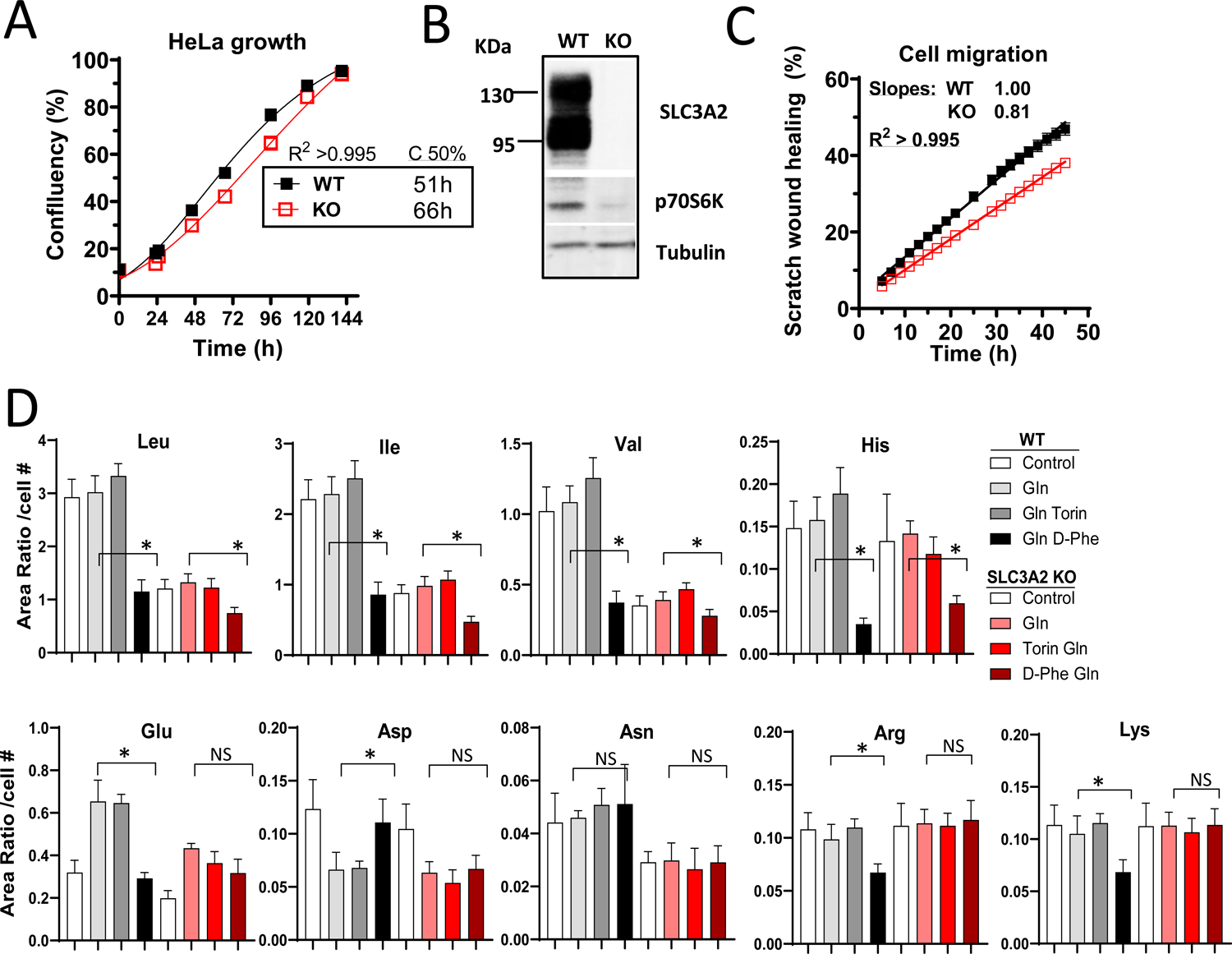
SLC3A2 KO cell phenotype. **(A)** Growth of WT and SLC3A2 KO HeLa cells cultured in normal conditions of DMEM +10% FCS. **(B)** Western blot of cells lysates in log phase growth and growth same conditions. **(C)** Cell motility in a lane-scrape wound healing assay. **(D)** LC-MS/MS analysis of metabolites in WT and SLC3A2 KO cells grown in DMEM + 10% FCS without Gln for 16h then supplemented with Gln (2 mM), Torin (250 nM) and D-phenylalanine (50 mM) in the final hour as indicated.

**Figure S3.**
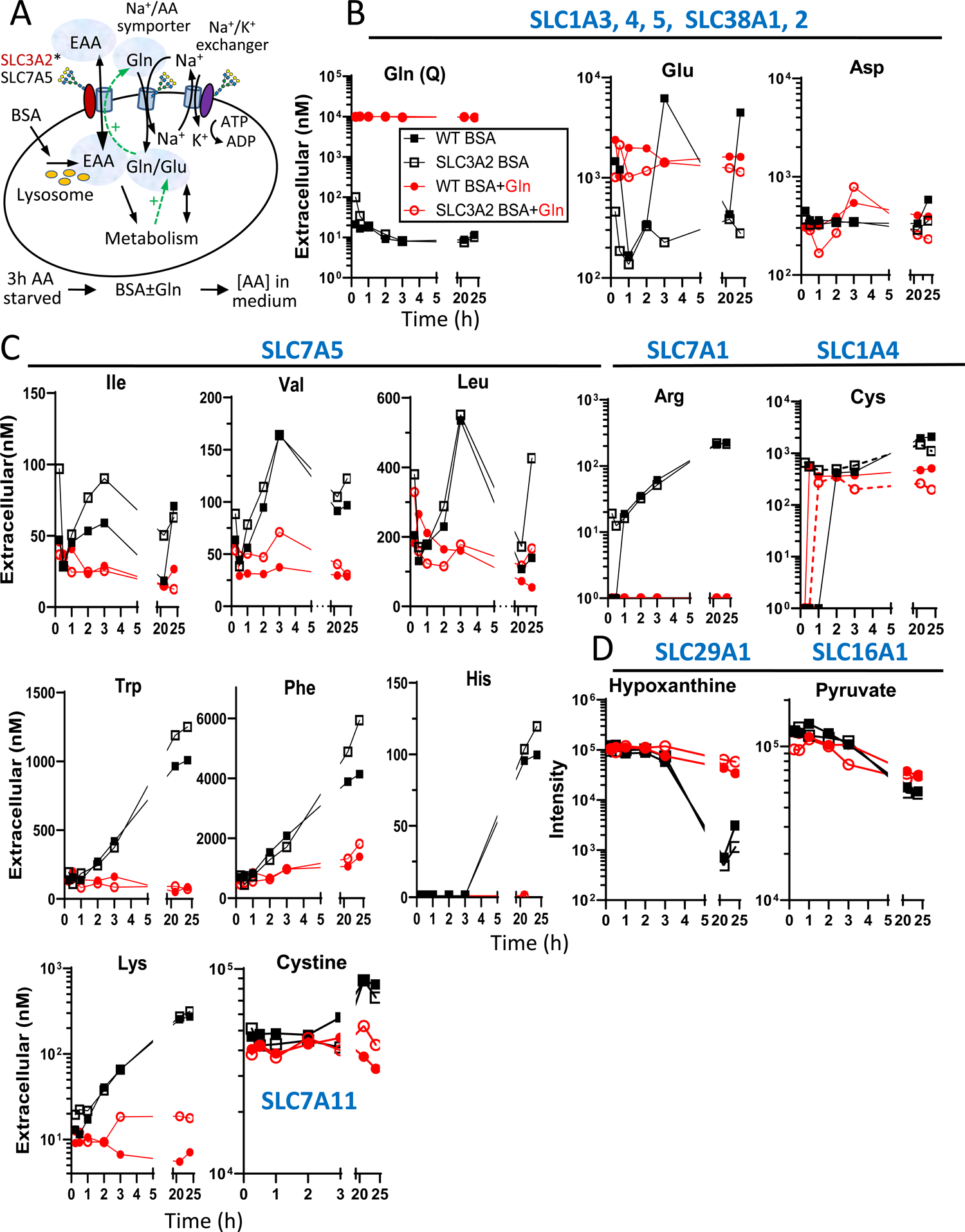
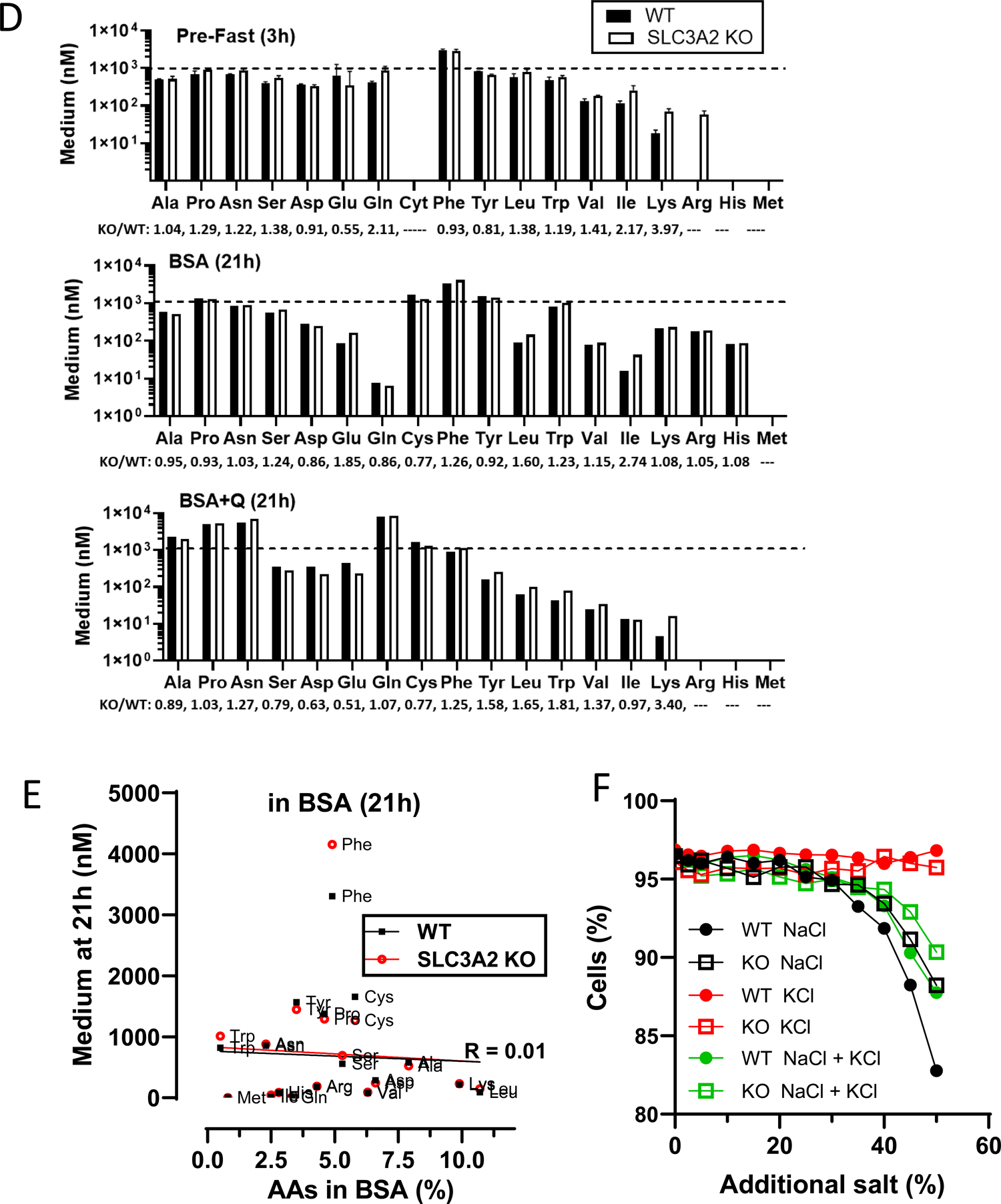
AA exchange with the culture medium. **(A)** Experimental scheme for AA derived from BSA and exchanged with medium depicted as SLC3A2*SLC7A5 may interact with Na^+^/AA symporters and Na^+^/K^+^ ATPase exchange. WT and SLC3A2 KO HeLa cells (6×10^5^ in 2 ml) were grown in AA-free medium for 3h followed by a change of medium and supplements of BSA or BSA+Gln and metabolites were measured in the medium by LC-MS/MS. **(B)** Gln, Glu, Asp. **(C)** BCAA, Trp, Phe, His, Lys, cystine **(D)** Consumption of hypoxanthine and pyruvate which were supplied in the AA-free Time (h) medium. Hypoxanthine salvage was markedly increased for cells in BSA alone indicating that purine biosynthesis is active in HeLa cells and requires Gln. Probable SLC transporters based on interactions with SLC3A2 (Fig. 6). **(D)** Amino acid levels in WT and SLC3A2 KO cells cultured in AA-free medium for 3h, as in panel A. Standard curves for each AA were used to calculate molar amounts. Standard curves for each AA were used to calculate molar amounts. **(E)** The fractional AA content in BSA does <u>not</u> correlate with levels in medium of cells supplemented with BSA. **(F)** SLC3A2 KO cells are more resistant to NaCl than WT cells consistent with reduced SLC7A exchanger interaction with Na+/AA symporter and Na+/K ATPase exchanger. Sensitivity is partially rescued by KCl.

**Figure S4.**
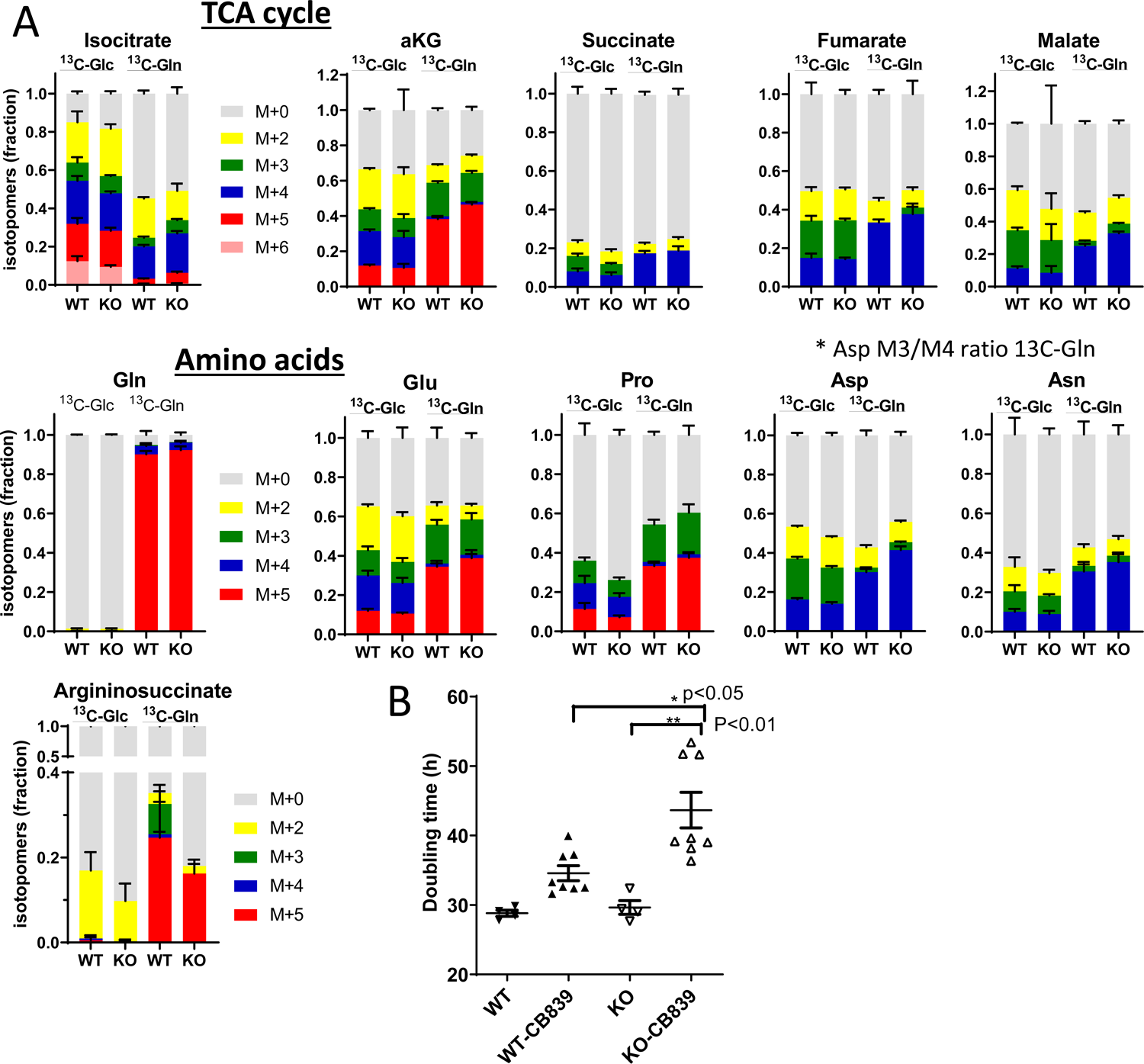
**(A)** Metabolic flux analysis. SLC3A2 WT and KO HeLa cells were cultured in medium with [U-^13^C]-glucose or [U-^13^C]-glutamine for 16h and the ^13^C-labeled fractions were quantified by LC-MS/MS. The results indicate more [U-^13^C]-glutamine and less [U-^13^C]-glucose flux in SLC3A2 KO cells into the TCA cycle in the oxidative direction. [U-^13^C]-glutamine flux to Asp (M4) was higher in SLC3A2 KO cells. *The Asp M3/M4 ratios from [U-^13^C]-glutamine labeling of Asp in KO 0.096 and WT 0.075, fumarate in KO 0.1 and WT 0, malate in KO 0.17 and WT 0.15, indicating a greater contribution of [U-^13^C]-glutamine to reductive conversion of aKG to citrate, and oxaloacetate transamination to Asp. **(B)** Doubling time in DMEM +10% FCS without and with 0.2 μM CB839, a glutaminase inhibitor.

**Figure S5.**
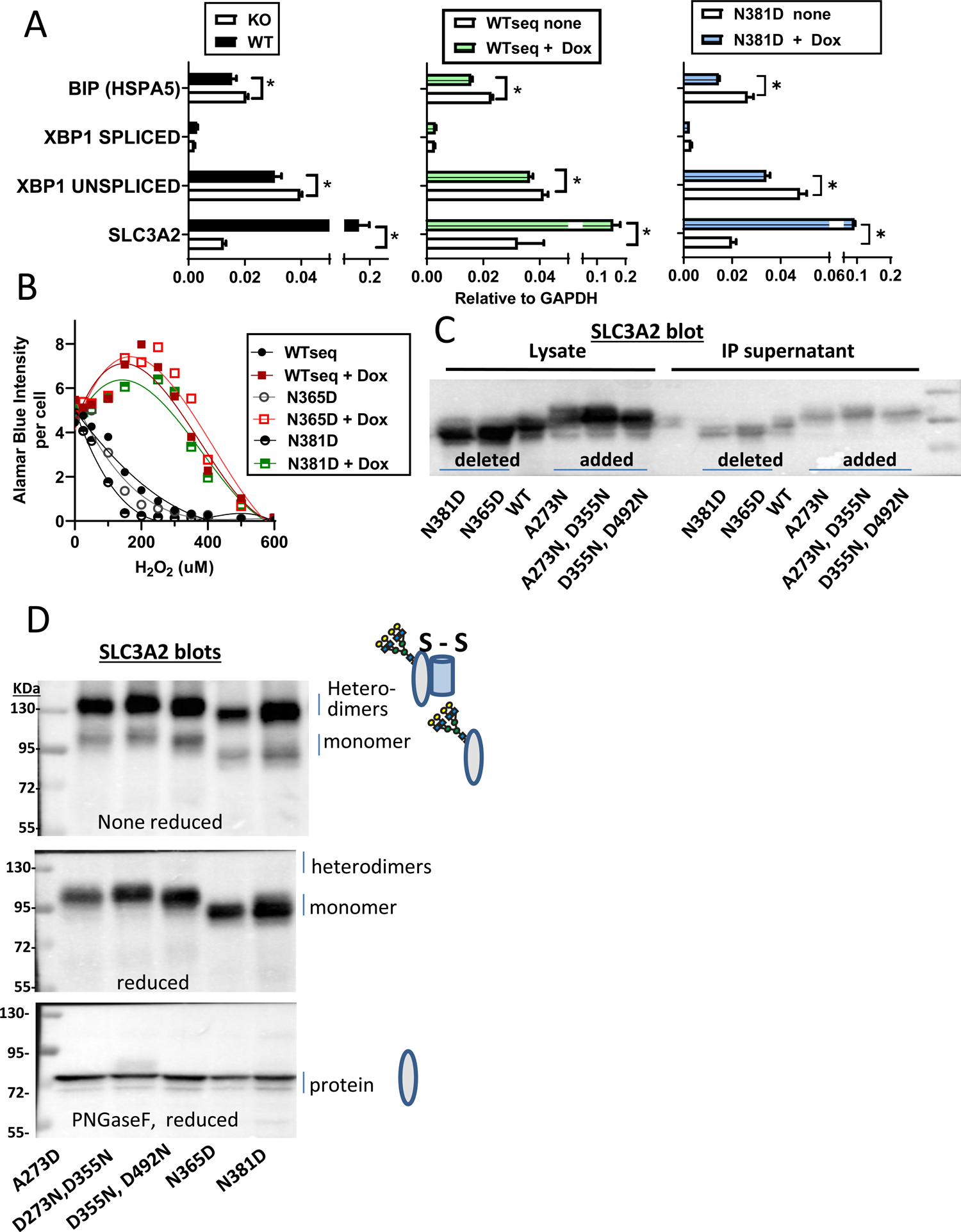
**(A)** Transcript levels by qPCR for stress inducible genes in WT and KO, Dox-induced WTseq and N381D. Cells were cultured in normal DMEM + 10% FCS conditions. **(B)** Sensitivity to H_2_O_2_. WTseq and variants +/- Dox. The Alamar Blue signal was normalized to morphologically-intact cells counted by InCell imaging. **(C**) Similar efficiency of FLAG-SLC3A2 pulldown for WTseq and site variants. **(D)** Western blot of dox-induced FLAG-SLC3A2, non-reduced, reduced, and reduced with PNGase digestion. The majority of FLAG-SLC3A2 is disulfide-linked as heterodimers.

**Figure S6:**
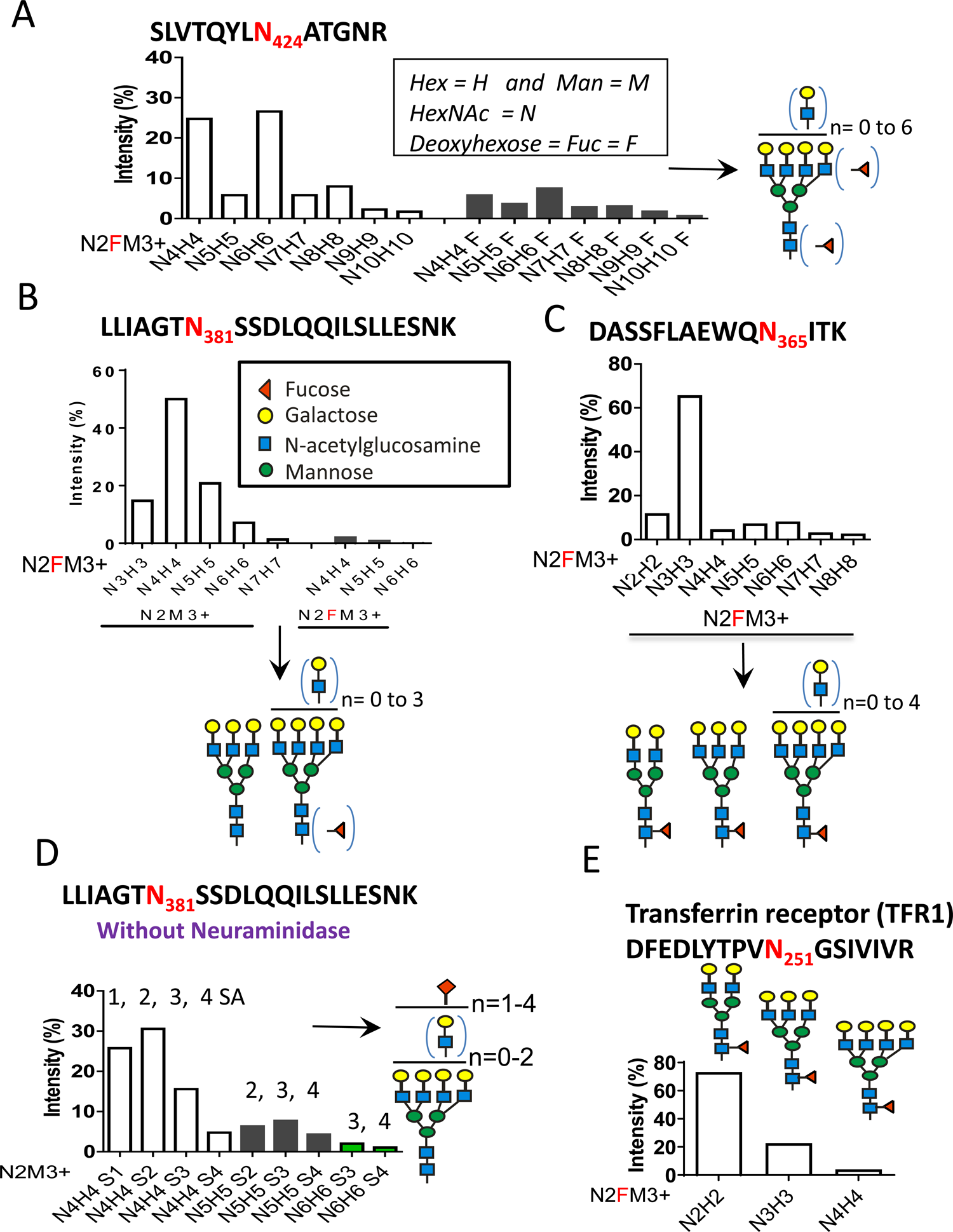
Site specific profiling of N-glycans of endogenous SLC3A2 from HeLa cell. **(A,B,C)** Glycoproteins from cell membrane preparation were separated by PAGE, and native SLC3A2 eluted from gel fractions. Tryptic peptides for three of the four sites in SLC3A2 were accessible to LC-MS/MS analysis. Sialic acid was removed with neuraminidase pre-treatment. **(D)** Analysis of the N381 glycopeptide without removing sialic acid, revealed a profile consistent with panel B. **(E)** By way of comparison, a transferrin receptor N-glycans at N251 from the same HeLa membrane preparation displayed ∼75% bi-, 20% tri and <5% tetra-antennary N-glycans, proportions closer to that of the total N-glycan pool.

**Figure S7:**
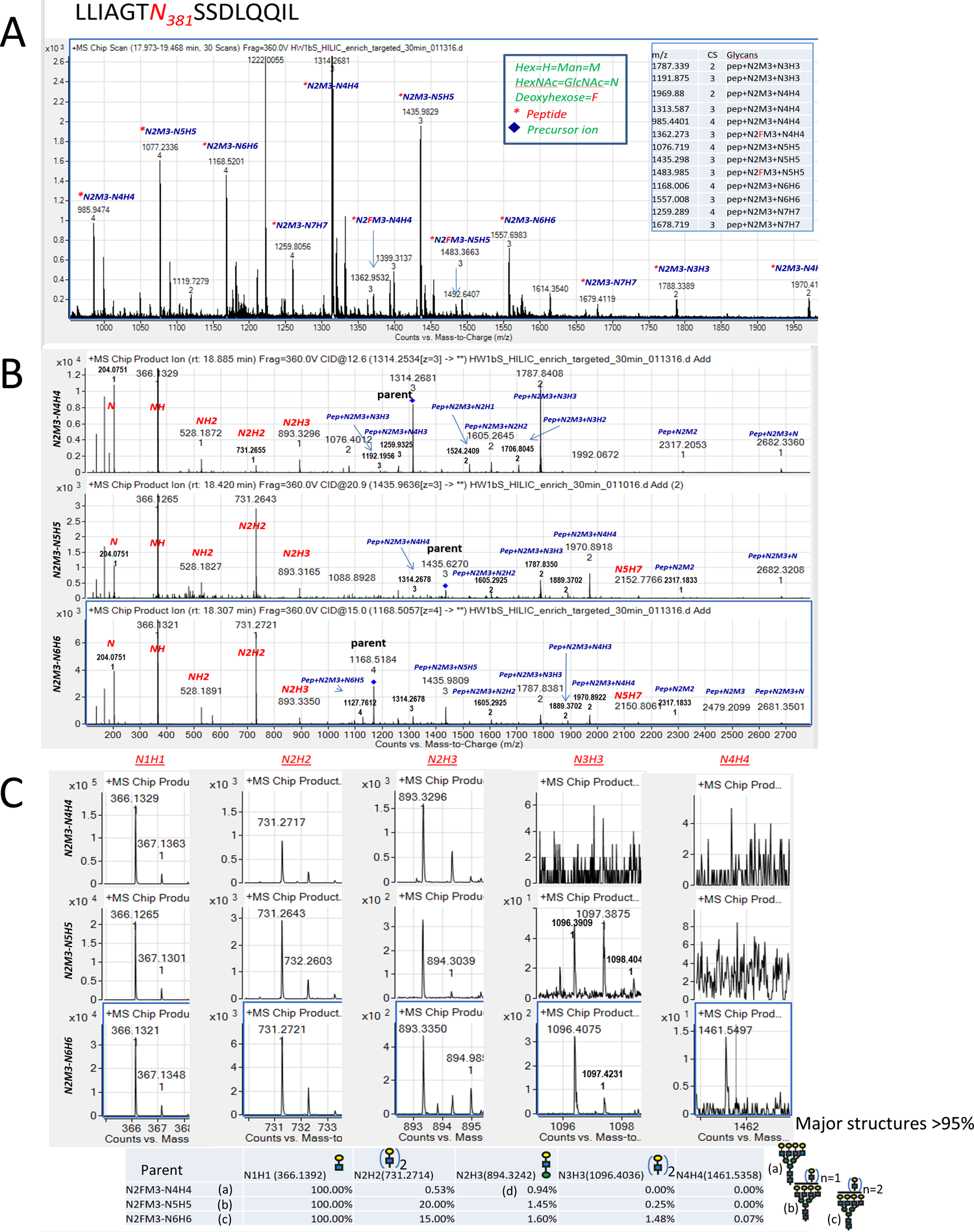
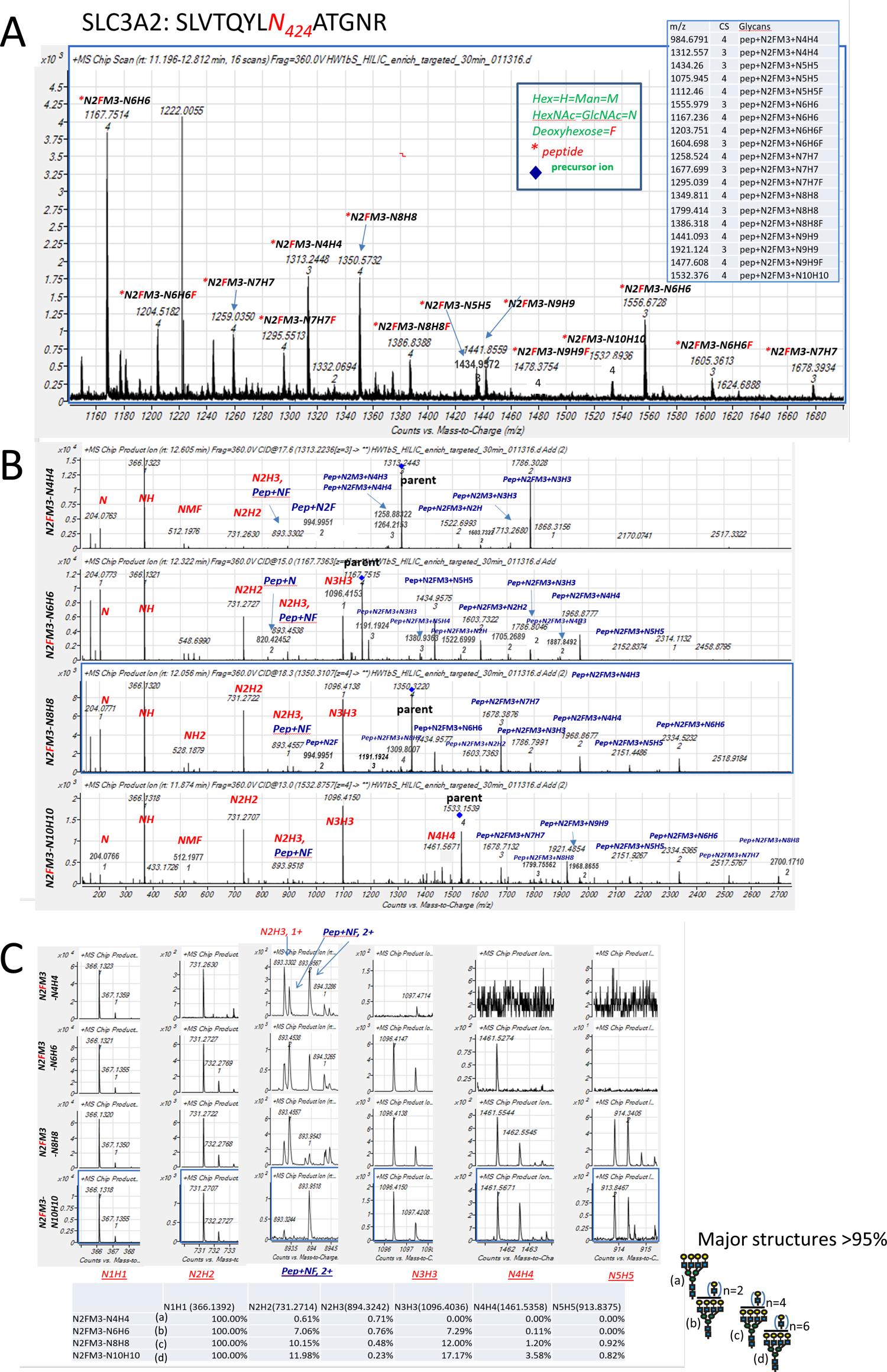
Detailed structural analysis of N-glycans by MS/MS at N381 and N424. **(A)** MS profile to identify major N-glycan species at N381. **(B)** Fragments identify fucose in core N-glycan and poly-LacNAc repeats. **(C)** Expansion of M/Z axis identifies stereoisomers of identical structures. The supplemental tables summarizes an interpretation of the results.

**Figure S8:**
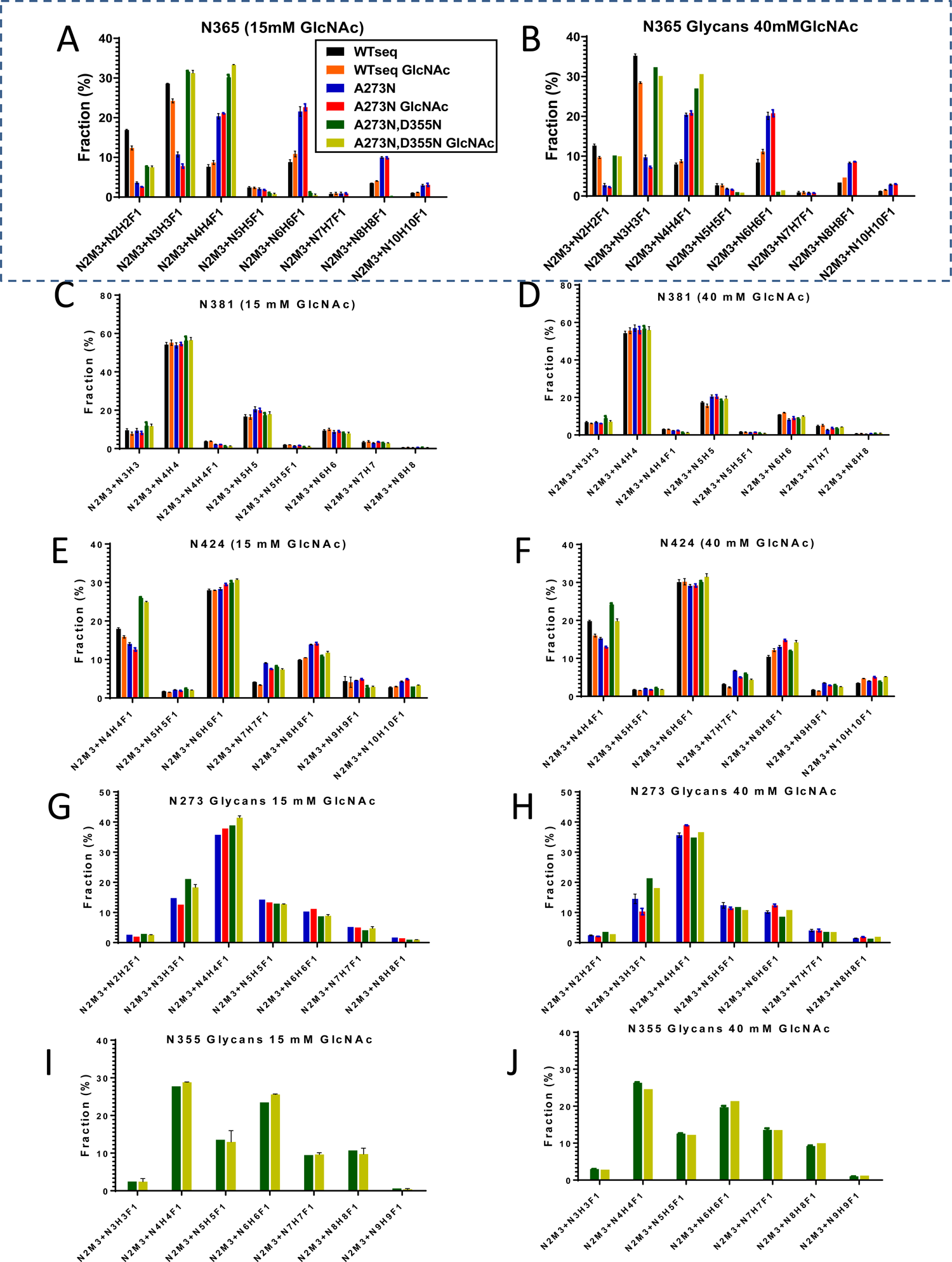
Site-specific analysis of FLAG-SLC3A2 WTseq and variants. Cells were cultured for 48h in medium supplemented with **(A,C,E,G,I)** 15 mM GlcNAc or **(B,D,F,H,J)** 40 mM GlcNAc. GlcNAc treatment increased branching at N365 and had no effect at the other sites. Mean ± SD of 3-4 technical replicates. Selection against sites at A273 and D355 in human SLC3A2 has removed the N-glycans and their influence on Golgi N-glycan processing at N365.

**Figure S9:**
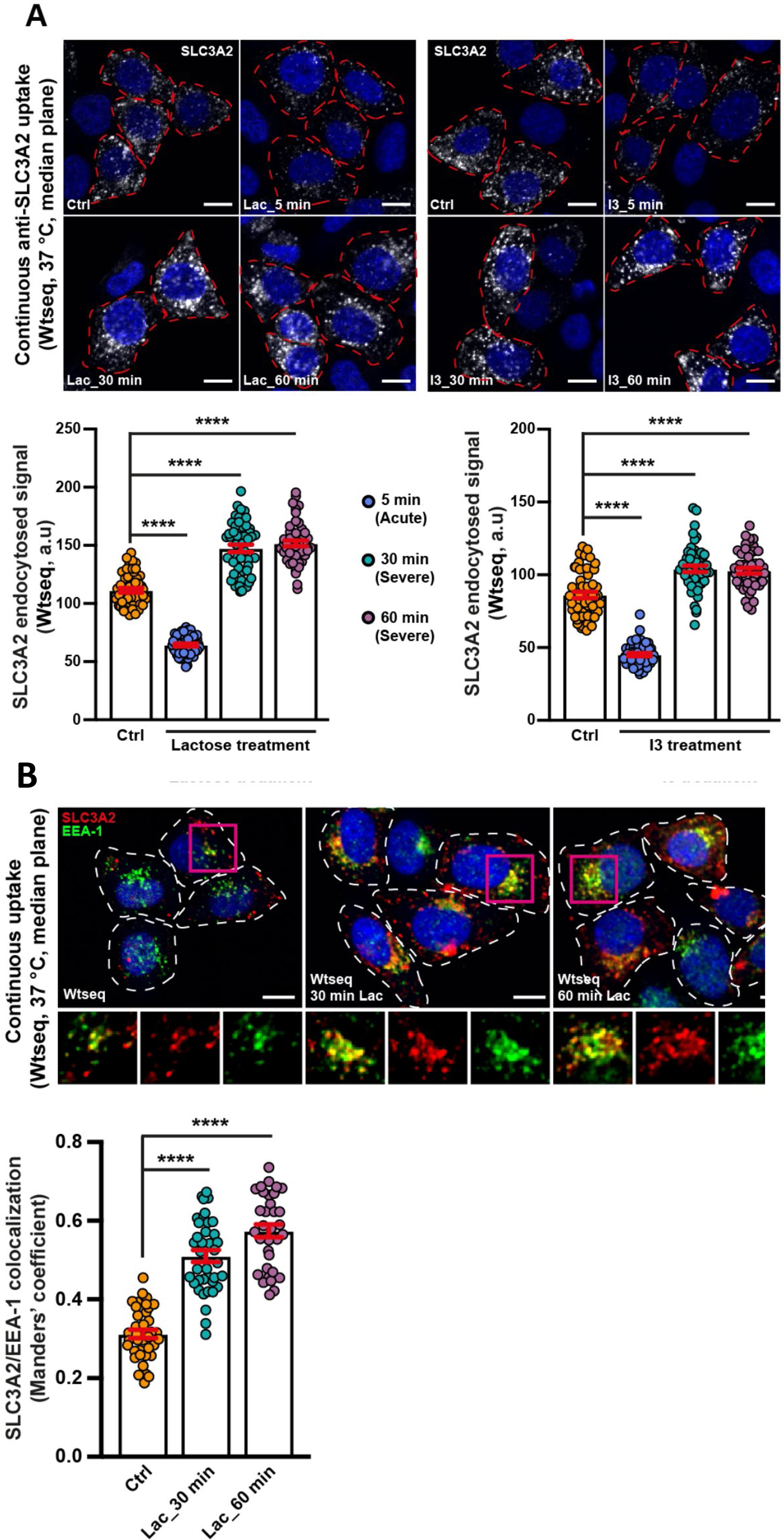
Comparison of acute or prolonged galectin inhibition on SLC3A2 uptake. (A) The internalization of SLC3A2 is affected in opposite ways by acute or prolonged galectin inhibition. Cells expressing the SLC3A2WTseq cells were pretreated for 5 (acute), 30 or 60 min (prolonged) with either **(i)** Gal3 specific inhibitor I3, or **(ii)** the general galectin inhibitor lactose (see methods), followed by anti-SLC3A2 antibody uptake assay (see methods). Internalized SLC3A2 signal intensities were quantified in both I3 and lactose conditions (histograms). With both inhibitors, short acute treatment (5 min) induced a significant inhibition of SLC3A2 endocytosis, whereas SLC3A2 uptake was increased with the prolonged treatment schedules. Note that in the latter cases, the pattern of distribution of internalized SLC3A2 appeared different from control conditions (see below). Means are ± SEM; Statistics are by one-way ANOVA, ****p < 0.0001. A single median plane from confocal imaging were represented. Red dashed lines represent the cell contour. Nuclei in blue (DAPI). Scale bars = 10 μm.. (B) **Prolonged galectin inhibition leads to increased SLC3A2 accumulation in perinuclear EEA-1 positive endosomes.** Experiment as in above on the prolonged lactose treatment condition, with immunolabeling of early endosomes using an anti-EEA-1 antibody (green). SLC3A2 colocalization with EEA1 positive structures was quantified (histogram). Note that in the severe lactose treatment condition, the colocalization of internalized SLC3A2 with EEA-1 was increased, which indicated that the protein was internalized via a different mechanism (see text for discussion). Means are ± SEM; Statistics are by one-way ANOVA, ****p < 0.0001. Zooms (lower panel) are from fuchsia boxed areas in the upper images. A single median plane from confocal imaging are represented. White dashed lines represent the cell contour. Nuclei in blue (DAPI). Scale bars = 10 μm

**Figure S10:**
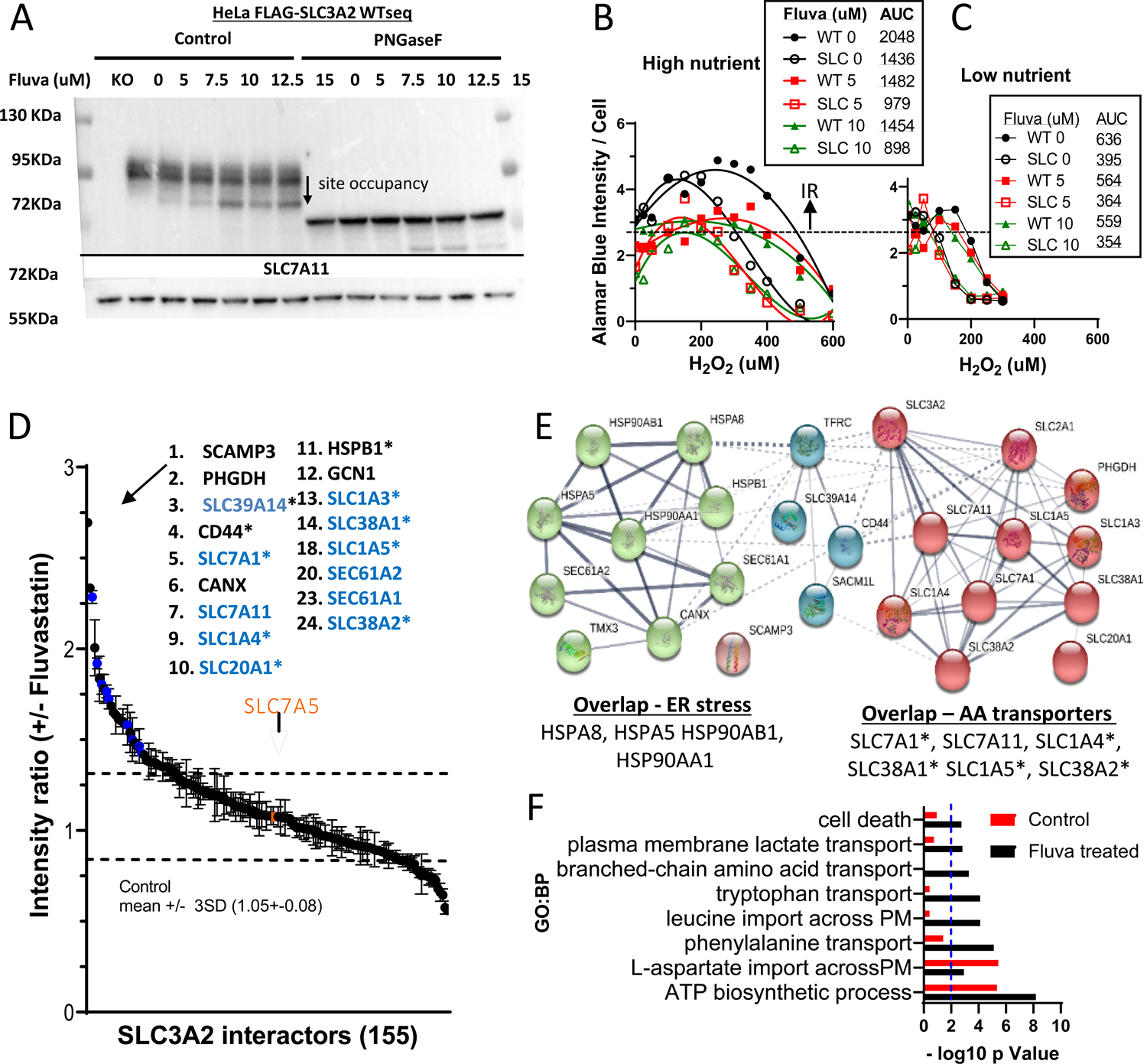
SLC3A2 and site variant interactions in Fluvastatin-stressed cells. **(A)** Western blots for dox-induced FLAG-SLC3A2 (WTseq) and SLC7A11 in lysates from HeLa cells, pretreated for 48h with fluvastatin. Aliquots of lysate were treated with PNGase F (right). **(B,C)** WT and SLC3A2 KO cells were pretreated with fluvastatin for 48 h in high and low glucose/glutamine DMEM (as in Fig. 3). H_2_O_2_ was added for an additional 16h followed by Alamar Blue for 90 min to measure reducing-potential. Cells were stained with DAPI to count morphologically-intact cells by InCell imaging. The early rise in Alamar Blue signal (i.e. NAD(P)H / viable cell) indicates an induced-resistance (IR) to low levels of H_2_O_2_. Area under the curve (AUC) in tet-induced cells ± fluvastatin. **(D)** AP-MS/MS analysis of pulldowns from tet-induced FLAG**-**SLC3A2 WTseq cells (72 h) without and with 10uM fluvastatin for the final 24 h. Data is the mean ± SD of 3 experiments. * N-glycosylated proteins. **(E)** A STRING networks of top 36 interactors with increased intensities upon fluva-treatment (panel D). The overlap is with the A277N variants (green), and with N381D and N365D variants (red) see Figure 7. **(F)** gProfiler multiquery for the AP-MS 155 list, ordered by intensity ratios to compare panel S10D (treated) with Fig 6A (untreated). The GO:BP modules with as a result of changes in gene order caused by Fluvastatin are shown.

**Figure S11:**
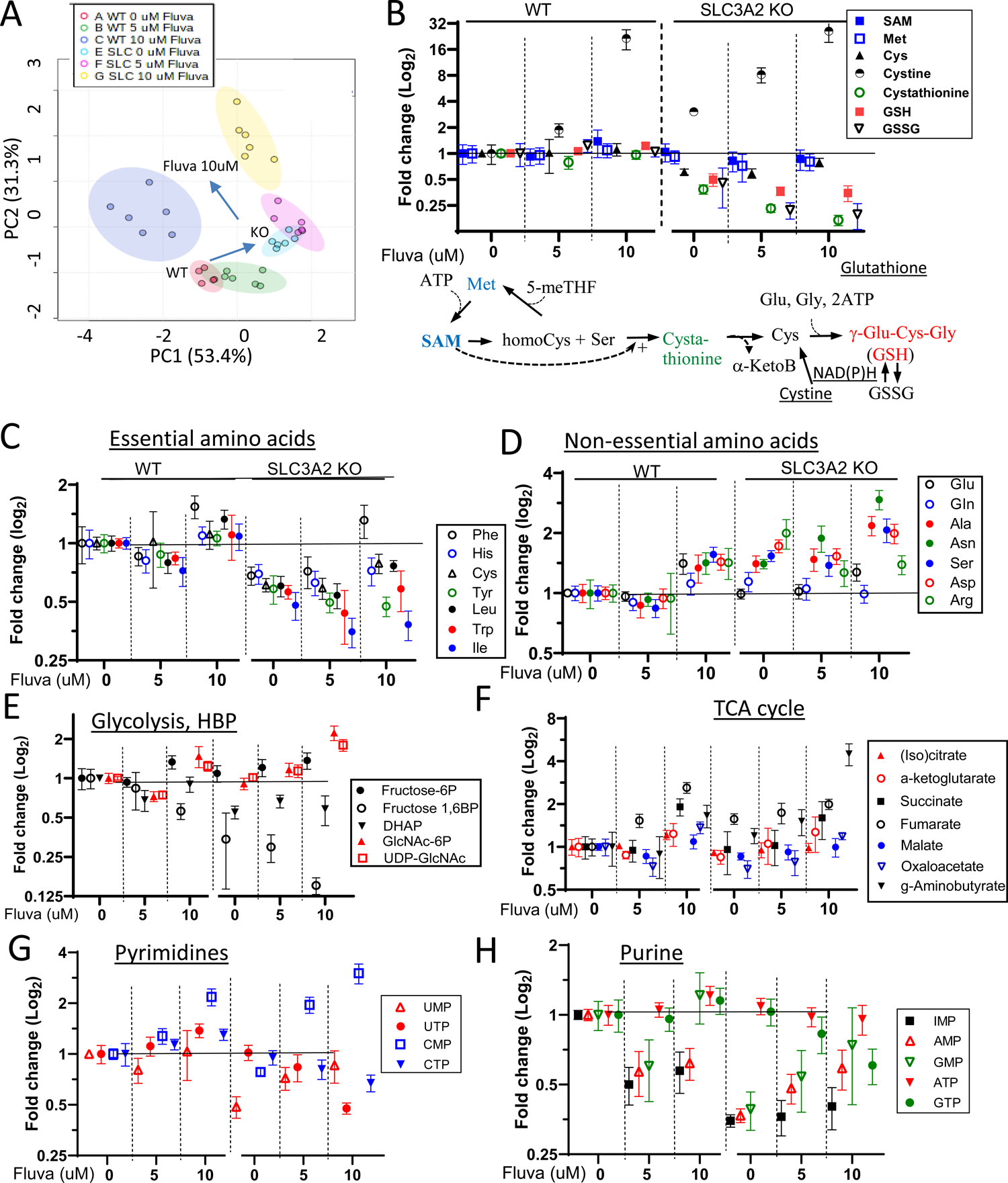
Metabolic profile of Fluvastatin treated WT and SLC3A2 KO cells. **(A)** Principal Component Analysis for metabolites from WT and SLC3A2 KO cells treated with 0, 5, 10 μM fluvastatin for 24h, measured by LC-MS/MS, mean ± SD (n=6). **(B)** Cystine levels were increased in SLC3A2 KO HeLa cells and further upon fluvastatin treatment, but GSH and GSSH levels were depleted, perhaps due to reduced levels of NADH which is required for conversion of Cystine to Cys. **(C-H)** Cells were treated with fluvastatin for 24h. Intracellular metabolites were measured by LC-MS/MS and displayed as KO/WT ratios, mean ± SD (n=6 technical replicates).

**Figure S12.**
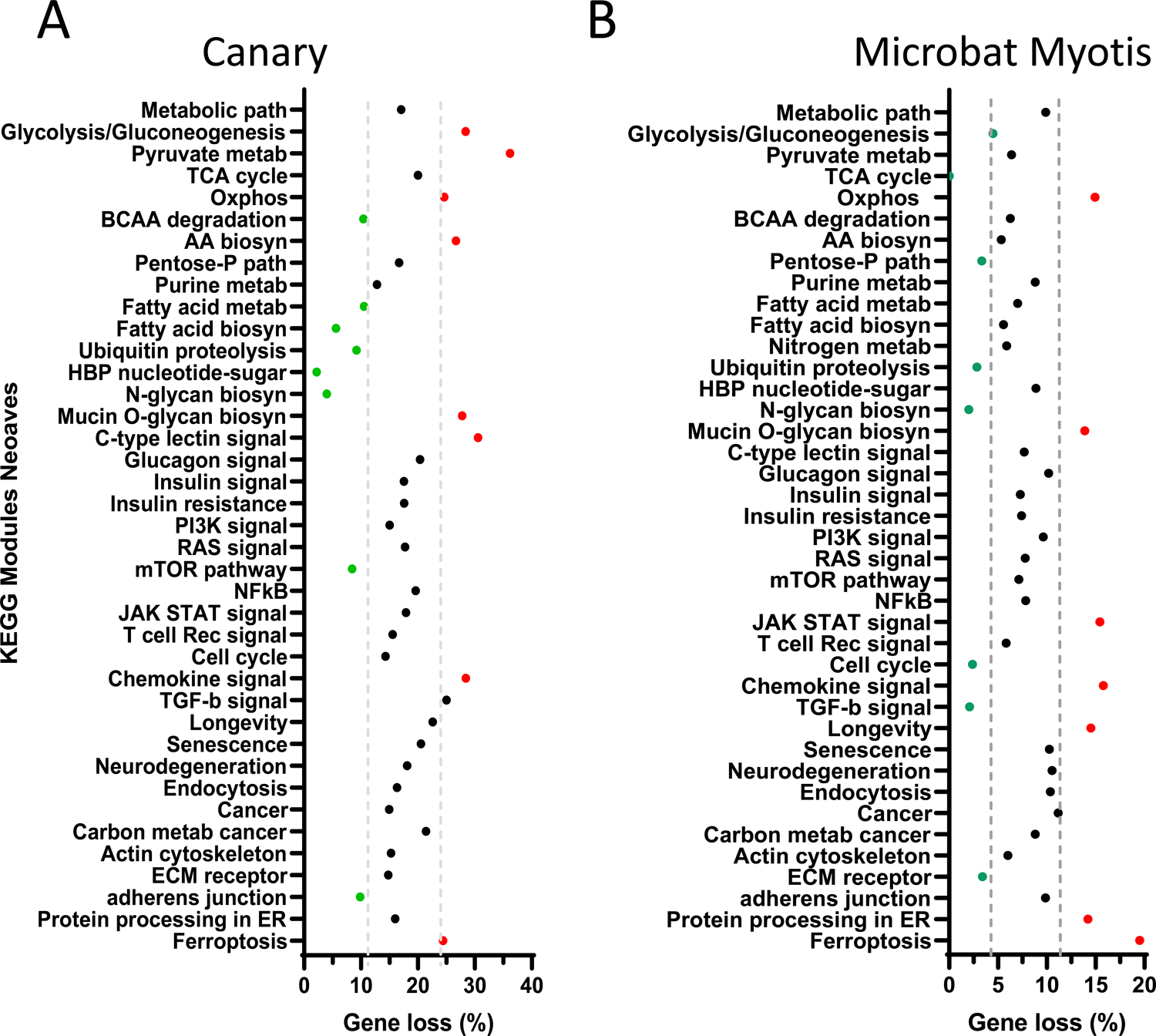
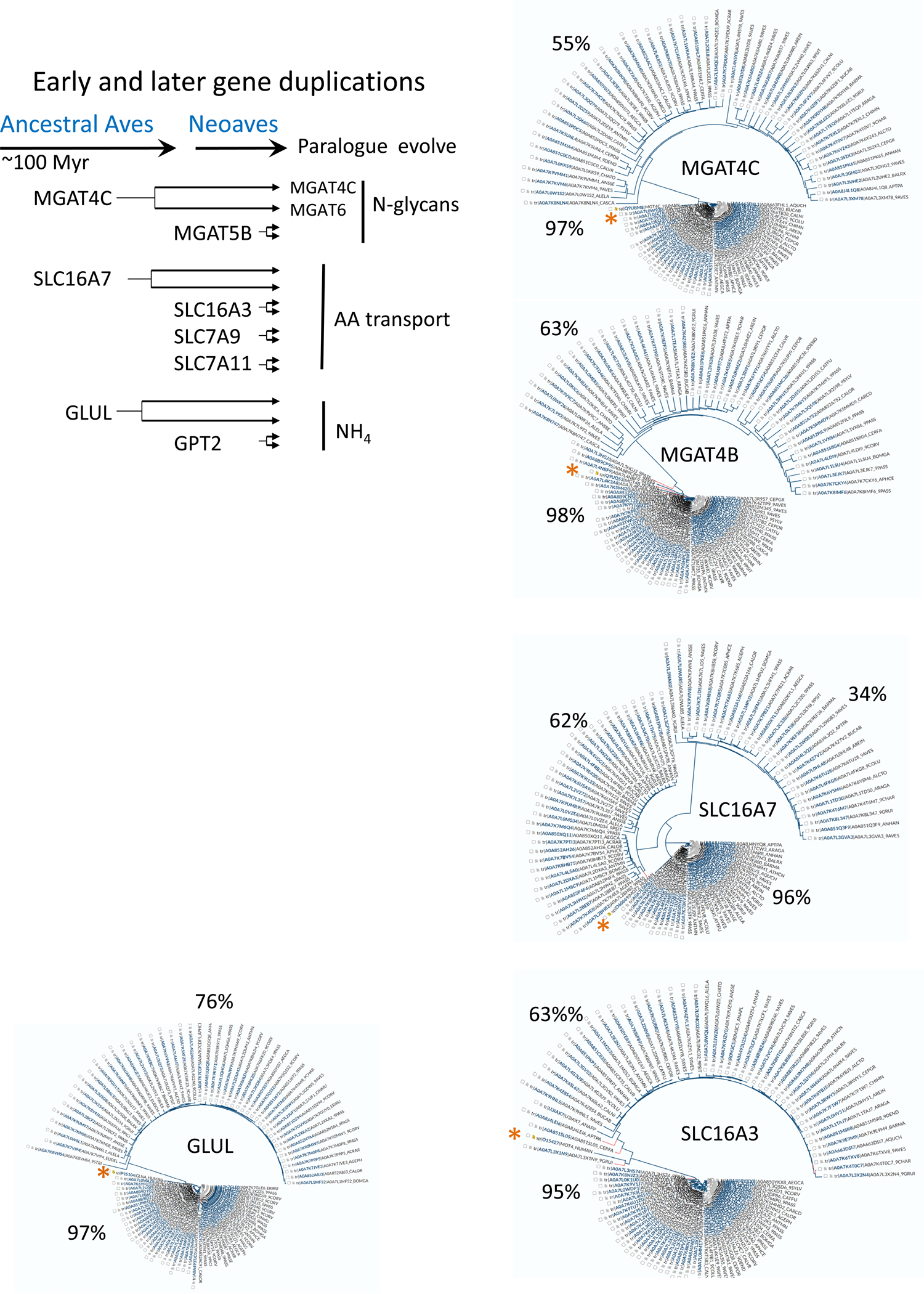
**(A)** gProfiler analysis of 45 KEGG modules for genes lost in Neoaves (n = 3730 genes total; 624 are absent Table S7). % absent for the 45 modules and doted lines is +/- 3 SD. **(B)** same analysis for Microbat. **(C) Examples of genes duplicated leading to Neoaves.** The phylograms show three typical examples with a conversed paralogues closer to the human gene and a rapidly evolving paralogue. SLC16A3 is unusual as the human gene also shows rapid evolution relative to the conserved bird orthologue. The phylograms were generated using Uniprot from a list of bird species (Ave 8782) in taking species by alphabetical order and using the first 100 of ∼500 sequences (2 paralogues per species). Percent identities between paralogues (clusters). Experiential evidence for functional divergence has shown only for MGAT4C. Single orthologues are present in Zebra fish, consistent with ancient origins. *human sequence.

**Figure S13.**
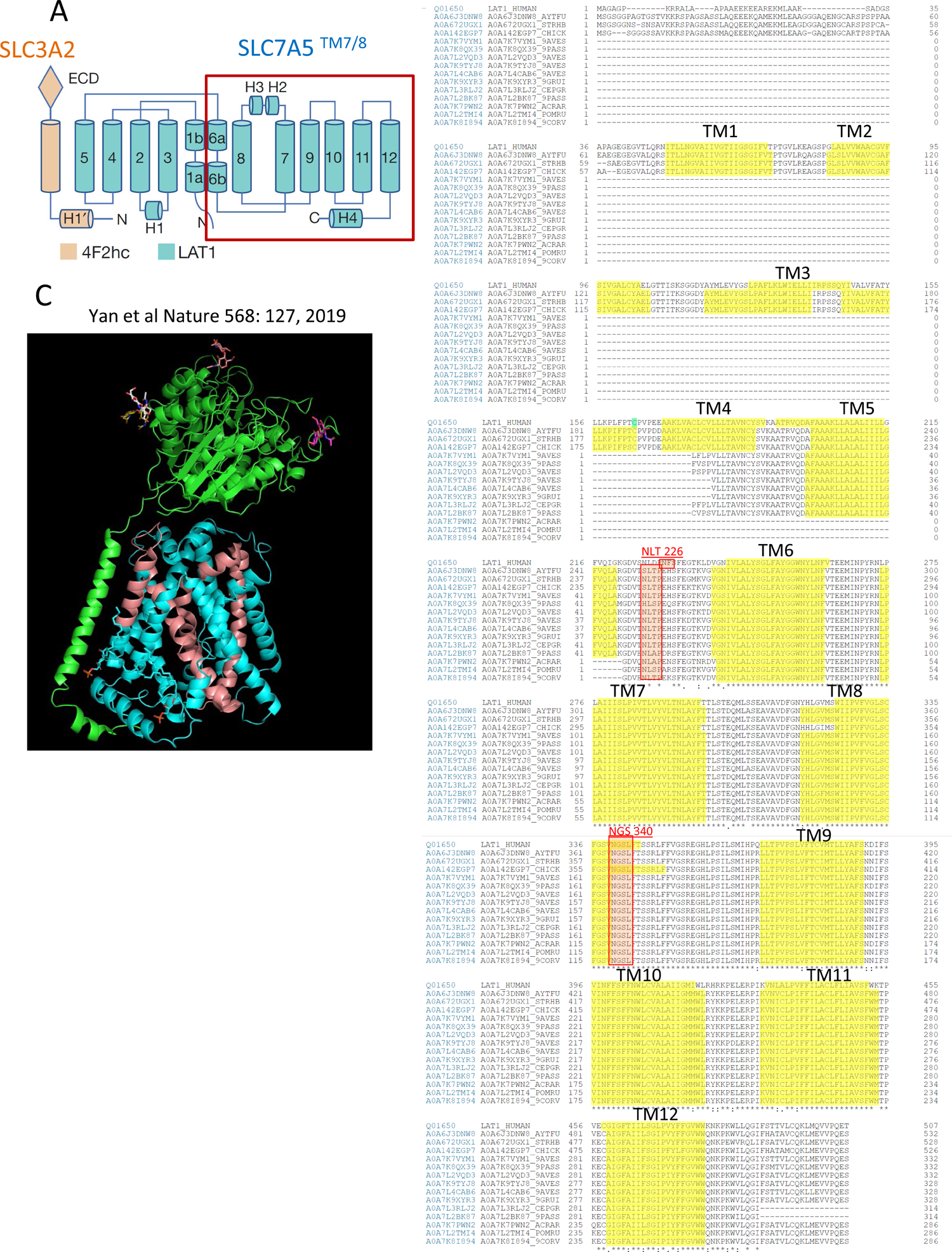
**(A)** SLC3A2 and SLC7A5 domains **(B)** alignment of human SLC7A5 with bird sequences reveals the 12 TM, in 42 ancestral Aves species and truncation at the N-terminus in Neoave species, leaving 8 or 7 TM domains. The gene sequence for 5-12 TM at the COOH end is conserved ∼ 90% identical. Two possible N-glycosylation site are marked (red). **(C)** TMs 1-5 appear to be critical to SLC7A5 fold, and loss has likely disrupted the structure and eliminates activity. A modified activity is of course possible in Neoaves.

**Figure S14:**
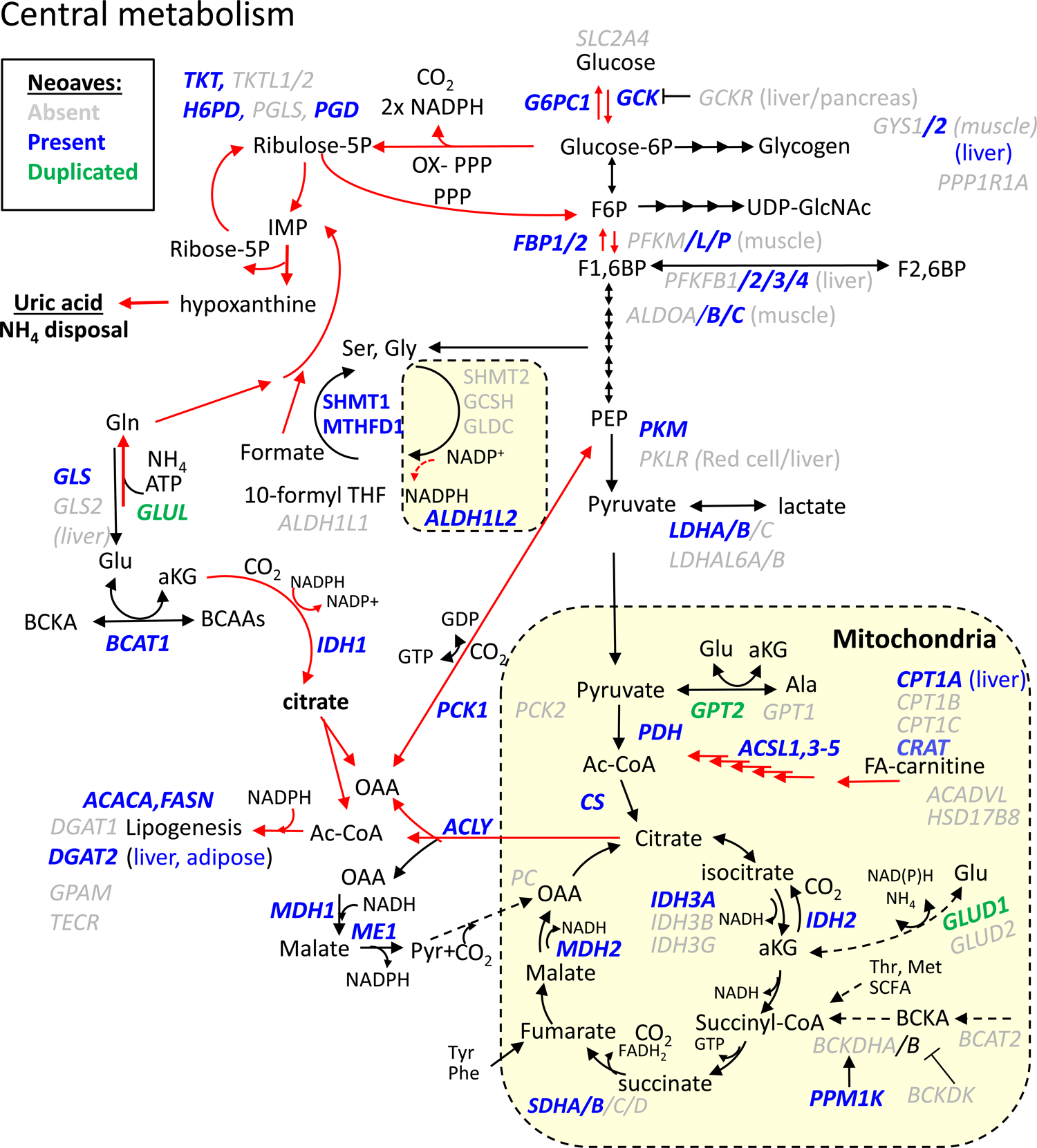
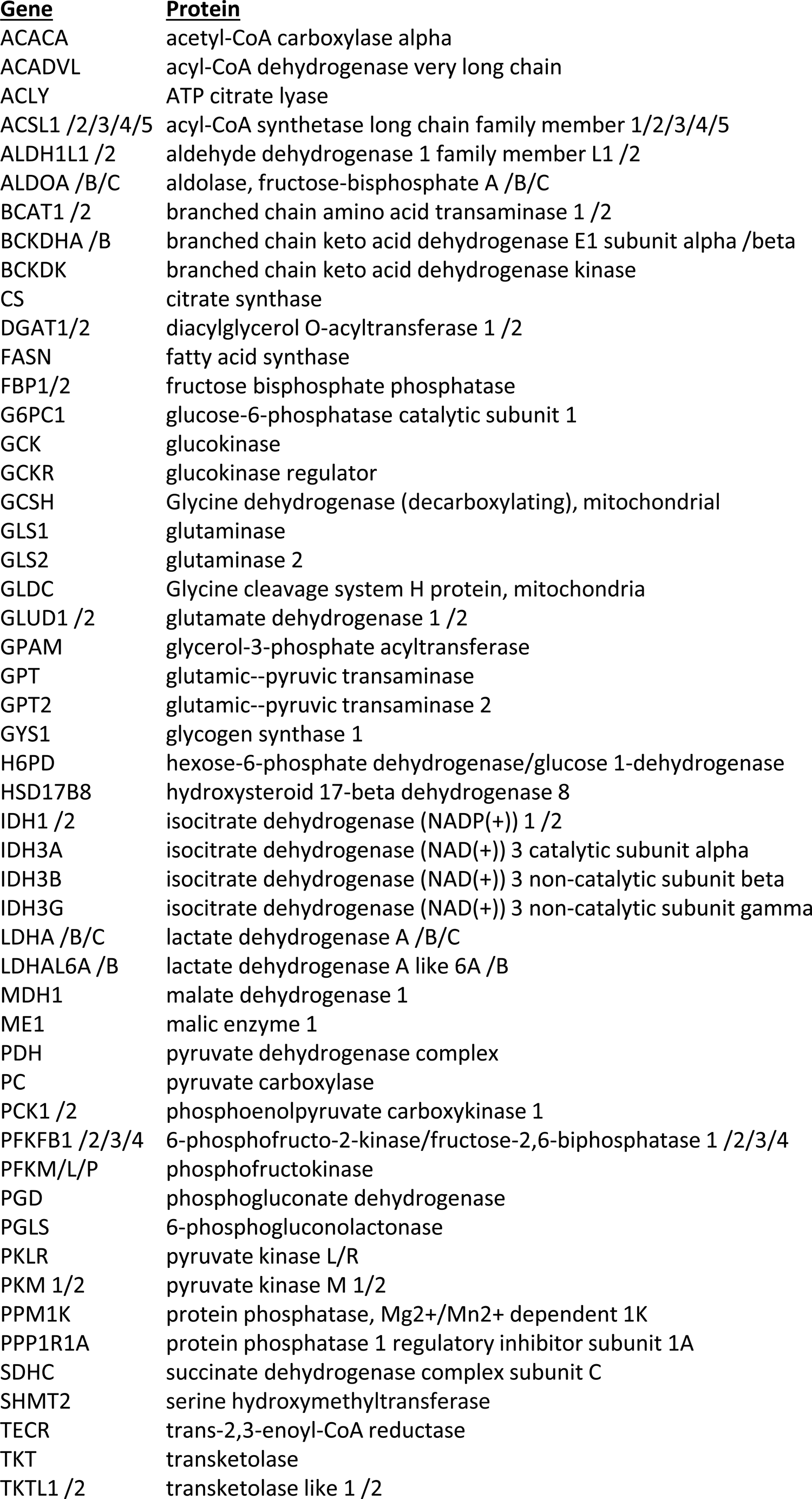
Streamlining in central metabolism. Genes present (blue), absent (grey) or duplicated (green) in Neoaves using Canary as primary reference. Red arrows are likely enhanced routes of flux, due in part to losses of GCKR, PC, PCK2, BCAT2. Other proteins associated with BCAA dehydrogenase complex are present; ACAD8, ACADSB, HIBADH, HIBCH, IVD and DLD, while HSD17B10 is absent. See following page for gene-enzyme list.

## REFERENCES

1. Kaochar, S., and Tu, B. P. (2012) Gatekeepers of chromatin: Small metabolites elicit big changes in gene expression. Trends Biochem Sci 37, 477–483

2. Wellen, K. E., and Thompson, C. B. (2012) A two-way street: reciprocal regulation of metabolism and signalling. Nat Rev Mol Cell Biol 13, 270–276

3. Araujo, L., Khim, P., Mkhikian, H., Mortales, C. L., and Demetriou, M. (2017) Glycolysis and glutaminolysis cooperatively control T cell function by limiting metabolite supply to N-glycosylation. Elife 6

4. Shang, J., Gao, N., Kaufman, R. J., Ron, D., Harding, H. P., and Lehrman, M. A. (2007) Translation attenuation by PERK balances ER glycoprotein synthesis with lipid-linked oligosaccharide flux. J Cell Biol 176, 605–616

5. Yu, R., Longo, J., van Leeuwen, J. E., Zhang, C., Branchard, E., Elbaz, M., Cescon, D. W., Drake, R. R., Dennis, J. W., and Penn, L. Z. (2021) Mevalonate pathway inhibition slows breast cancer metastasis via reduced N-glycosylation abundance and branching. Cancer Res

6. Partridge, E. A., Le Roy, C., Di Guglielmo, G. M., Pawling, J., Cheung, P., Granovsky, M., Nabi, I. R., Wrana, J. L., and Dennis, J. W. (2004) Regulation of cytokine receptors by Golgi N-glycan processing and endocytosis. Science 306, 120–124

7. Demetriou, M., Granovsky, M., Quaggin, S., and Dennis, J. W. (2001) Negative regulation of T-cell activation and autoimmunity by Mgat5 N-glycosylation. Nature 409, 733–739

8. Lagana, A., Goetz, J. G., Cheung, P., Raz, A., Dennis, J. W., and Nabi, I. R. (2006) Galectin binding to Mgat5-modified N-glycans regulates fibronectin matrix remodeling in tumor cells. Mol Cell Biol 26, 3181–3193

9. Johswich, A., Longuet, C., Pawling, J., Rahman, A. A., Ryczko, M., Drucker, D. J., and Dennis, J. W. (2014) N-Glycan Remodeling on Glucagon Receptor Is an Effector of Nutrient Sensing by the Hexosamine Biosynthesis Pathway. J Biol Chem 289, 15927–15941

10. Ohtsubo, K., Takamatsu, S., Gao, C., Korekane, H., Kurosawa, T. M., and Taniguchi, N. (2013) N-Glycosylation modulates the membrane sub-domain distribution and activity of glucose transporter 2 in pancreatic beta cells. Biochem Biophys Res Commun 434, 346–351

11. Abdel Rahman, A. M., Ryczko, M., Nakano, M., Pawling, J., Rodrigues, T., Johswich, A., Taniguchi, N., and Dennis, J. W. (2015) Golgi N-glycan branching N-acetylglucosaminyltransferases I, V and VI promote nutrient uptake and metabolism. Glycobiology 25, 225–240

12. Wellen, K. E., Lu, C., Mancuso, A., Lemons, J. M., Ryczko, M., Dennis, J. W., Rabinowitz, J. D., Coller, H. A., and Thompson, C. B. (2010) The hexosamine biosynthetic pathway couples growth factor-induced glutamine uptake to glucose metabolism. Genes Dev 24, 2784–2799

13. Dennis, J. W., and Brewer, C. F. (2013) Density-dependent lectin-glycan interactions as a paradigm for conditional regulation by posttranslational modifications. Mol Cell Proteomics 12, 913–920

14. Dam, T. K., Gerken, T. A., and Brewer, C. F. (2009) Thermodynamics of multivalent carbohydrate-lectin cross-linking interactions: importance of entropy in the bind and jump mechanism. Biochemistry 48, 3822–3827

15. Lakshminarayan, R., Wunder, C., Becken, U., Howes, M. T., Benzing, C., Arumugam, S., Sales, S., Ariotti, N., Chambon, V., Lamaze, C., Loew, D., Shevchenko, A., Gaus, K., Parton, R. G., and Johannes, L. (2014) Galectin-3 drives glycosphingolipid-dependent biogenesis of clathrin-independent carriers. Nat Cell Biol 16, 595–606

16. Renard, H. F., Tyckaert, F., Lo Giudice, C., Hirsch, T., Valades-Cruz, C. A., Lemaigre, C., Shafaq-Zadah, M., Wunder, C., Wattiez, R., Johannes, L., van der Bruggen, P., Alsteens, D., and Morsomme, P. (2020) Endophilin-A3 and Galectin-8 control the clathrin-independent endocytosis of CD166. Nat Commun 11, 1457

17. Dennis, J. W. (2015) Many Light Touches Convey the Message. Trends Biochem Sci 40, 673–686

18. Stanley, P. (2016) What Have We Learned from Glycosyltransferase Knockouts in Mice? J Mol Biol 428, 3166–3182

19. Aregger, M., Lawson, K. A., Billmann, M., Costanzo, M., Tong, A. H. Y., Chan, K., Rahman, M., Brown, K. R., Ross, C., Usaj, M., Nedyalkova, L., Sizova, O., Habsid, A., Pawling, J., Lin, Z. Y., Abdouni, H., Wong, C. J., Weiss, A., Mero, P., Dennis, J. W., Gingras, A. C., Myers, C. L., Andrews, B. J., Boone, C., and Moffat, J. (2020) Systematic mapping of genetic interactions for de novo fatty acid synthesis identifies C12orf49 as a regulator of lipid metabolism. Nat Metab 2, 499–513

20. Veerababu, G., Tang, J., Hoffman, R. T., Daniels, M. C., Hebert, L. F., Jr., Crook, E. D., Cooksey, R. C., and McClain, D. A. (2000) Overexpression of glutamine: fructose-6-phosphate amidotransferase in the liver of transgenic mice results in enhanced glycogen storage, hyperlipidemia, obesity, and impaired glucose tolerance. Diabetes 49, 2070–2078

21. Kamada, Y., Ebisutani, Y., Kida, S., Mizutani, K., Akita, M., Yamamoto, A., Fujii, H., Sobajima, T., Terao, N., Takamatsu, S., Yoshida, Y., Takehara, T., and Miyoshi, E. (2016) Ectopic expression of N-acetylglucosaminyltransferase V accelerates hepatic triglyceride synthesis. Hepatol Res 46, E118–129

22. Cheung, P., Pawling, J., Partridge, E. A., Sukhu, B., Grynpas, M., and Dennis, J. W. (2007) Metabolic homeostasis and tissue renewal are dependent on beta1,6GlcNAc-branched N-glycans. Glycobiology 17, 828–837

23. Cummings, N. E., Williams, E. M., Kasza, I., Konon, E. N., Schaid, M. D., Schmidt, B. A., Poudel, C., Sherman, D. S., Yu, D., Arriola Apelo, S. I., Cottrell, S. E., Geiger, G., Barnes, M. E., Wisinski, J. A., Fenske, R. J., Matkowskyj, K. A., Kimple, M. E., Alexander, C. M., Merrins, M. J., and Lamming, D. W. (2018) Restoration of metabolic health by decreased consumption of branched-chain amino acids. J Physiol 596, 623–645

24. Ryczko, M. C., Pawling, J., Chen, R., Abdel Rahman, A. M., Yau, K., Copeland, J. K., Zhang, C., Surendra, A., Guttman, D. S., Figeys, D., and Dennis, J. W. (2016) Metabolic Reprogramming by Hexosamine Biosynthetic and Golgi N-Glycan Branching Pathways. Sci Rep 6, 23043

25. Cantor, J. M., and Ginsberg, M. H. (2012) CD98 at the crossroads of adaptive immunity and cancer. J Cell Sci 125, 1373–1382

26. Nicklin, P., Bergman, P., Zhang, B., Triantafellow, E., Wang, H., Nyfeler, B., Yang, H., Hild, M., Kung, C., Wilson, C., Myer, V. E., Mackeigan, J. P., Porter, J. A., Wang, Y. K., Cantley, L. C., Finan, P. M., and Murphy, L. O. (2009) Bidirectional Transport of Amino Acids Regulates mTOR and Autophagy. Cell 136, 521–534

27. Gu, X., Orozco, J. M., Saxton, R. A., Condon, K. J., Liu, G. Y., Krawczyk, P. A., Scaria, S. M., Harper, J. W., Gygi, S. P., and Sabatini, D. M. (2017) SAMTOR is an S-adenosylmethionine sensor for the mTORC1 pathway. Science 358, 813–818

28. Hesketh, G. G., Papazotos, F., Pawling, J., Rajendran, D., Knight, J. D. R., Martinez, S., Taipale, M., Schramek, D., Dennis, J. W., and Gingras, A. C. (2020) The GATOR-Rag GTPase pathway inhibits mTORC1 activation by lysosome-derived amino acids. Science 370, 351–356

29. Napolitano, L., Scalise, M., Galluccio, M., Pochini, L., Albanese, L. M., and Indiveri, C. (2015) LAT1 is the transport competent unit of the LAT1/CD98 heterodimeric amino acid transporter. Int J Biochem Cell Biol 67, 25–33

30. Kantipudi, S., Jeckelmann, J. M., Ucurum, Z., Bosshart, P. D., and Fotiadis, D. (2020) The Heavy Chain 4F2hc Modulates the Substrate Affinity and Specificity of the Light Chains LAT1 and LAT2. Int J Mol Sci 21

31. Cormerais, Y., Giuliano, S., LeFloch, R., Front, B., Durivault, J., Tambutte, E., Massard, P. A., de la Ballina, L. R., Endou, H., Wempe, M. F., Palacin, M., Parks, S. K., and Pouyssegur, J. (2016) Genetic Disruption of the Multifunctional CD98/LAT1 Complex Demonstrates the Key Role of Essential Amino Acid Transport in the Control of mTORC1 and Tumor Growth. Cancer Res 76, 4481–4492

32. Ablack, J. N., Metz, P. J., Chang, J. T., Cantor, J. M., and Ginsberg, M. H. (2015) Ubiquitylation of CD98 limits cell proliferation and clonal expansion. J Cell Sci 128, 4273–4278

33. Liu, C., Li, X., Li, C., Zhang, Z., Gao, X., Jia, Z., Chen, H., Jia, Q., Zhao, X., Liu, J., Liu, B., Xu, Z., Tian, Y., and He, K. (2018) SLC3A2 is a novel endoplasmic reticulum stress-related signaling protein that regulates the unfolded protein response and apoptosis. PLoS One 13, e0208993

34. Shin, C. S., Mishra, P., Watrous, J. D., Carelli, V., D’Aurelio, M., Jain, M., and Chan, D. C. (2017) The glutamate/cystine xCT antiporter antagonizes glutamine metabolism and reduces nutrient flexibility. Nat Commun 8, 15074

35. Jiang, L., Kon, N., Li, T., Wang, S. J., Su, T., Hibshoosh, H., Baer, R., and Gu, W. (2015) Ferroptosis as a p53-mediated activity during tumour suppression. Nature 520, 57–62

36. Bergsbaken, T., Fink, S. L., and Cookson, B. T. (2009) Pyroptosis: host cell death and inflammation. Nat Rev Microbiol 7, 99–109

37. Schwertassek, U., Haque, A., Krishnan, N., Greiner, R., Weingarten, L., Dick, T. P., and Tonks, N. K. (2014) Reactivation of oxidized PTP1B and PTEN by thioredoxin 1. FEBS J 281, 3545–3558

38. Mendelsohn, R., Cheung, P., Berger, L., Partridge, E. A., Lau, K., Pawling, J., and Dennis, J. W. (2007) Control of tumor metabolism and growth by N-glycan processing Cancer Res 67, 9771-9780

39. Gnanapradeepan, K., Leu, J. I., Basu, S., Barnoud, T., Good, M., Lee, J. V., Quinn, W. J., Kung, C. P., Ahima, R., Baur, J. A., Wellen, K. E., Liu, Q., Schug, Z. T., George, D. L., and Murphy, M. E. (2020) Increased mTOR activity and metabolic efficiency in mouse and human cells containing the African-centric tumor-predisposing p53 variant Pro47Ser. Elife 9

40. Leu, J. I., Murphy, M. E., and George, D. L. (2019) Mechanistic basis for impaired ferroptosis in cells expressing the African-centric S47 variant of p53. Proc Natl Acad Sci U S A 116, 8390–8396

41. Tang, C. Y., Man, X. F., Guo, Y., Tang, H. N., Tang, J., Zhou, C. L., Tan, S. W., Wang, M., and Zhou, H. D. (2017) IRS-2 Partially Compensates for the Insulin Signal Defects in IRS-1(-/-) Mice Mediated by miR-33. Mol Cells 40, 123–132

42. Lewis, K. N., Wason, E., Edrey, Y. H., Kristan, D. M., Nevo, E., and Buffenstein, R. (2015) Regulation of Nrf2 signaling and longevity in naturally long-lived rodents. Proc Natl Acad Sci U S A 112, 3722–3727

43. Migliaccio, E., Giorgio, M., Mele, S., Pelicci, G., Reboldi, P., Pandolfi, P. P., Lanfrancone, L., and Pelicci, P. G. (1999) The p66shc adaptor protein controls oxidative stress response and life span in mammals. Nature 402, 309–313

44. Giorgio, M., Migliaccio, E., Orsini, F., Paolucci, D., Moroni, M., Contursi, C., Pelliccia, G., Luzi, L., Minucci, S., Marcaccio, M., Pinton, P., Rizzuto, R., Bernardi, P., Paolucci, F., and Pelicci, P. G. (2005) Electron transfer between cytochrome c and p66Shc generates reactive oxygen species that trigger mitochondrial apoptosis. Cell 122, 221–233

45. El Ansari, R., Craze, M. L., Diez-Rodriguez, M., Nolan, C. C., Ellis, I. O., Rakha, E. A., and Green, A. R. (2018) The multifunctional solute carrier 3A2 (SLC3A2) confers a poor prognosis in the highly proliferative breast cancer subtypes. Br J Cancer 118, 1115–1122

46. Estrach, S., Lee, S. A., Boulter, E., Pisano, S., Errante, A., Tissot, F. S., Cailleteau, L., Pons, C., Ginsberg, M. H., and Feral, C. C. (2014) CD98hc (SLC3A2) loss protects against ras-driven tumorigenesis by modulating integrin-mediated mechanotransduction. Cancer Res 74, 6878–6889

47. Tissot, F. S., Estrach, S., Boulter, E., Cailleteau, L., Tosello, L., Seguin, L., Pisano, S., Audebert, S., Croce, O., and Feral, C. C. (2018) Dermal Fibroblast SLC3A2 Deficiency Leads to Premature Aging and Loss of Epithelial Homeostasis. J Invest Dermatol 138, 2511–2521

48. Broer, S. (2020) Amino Acid Transporters as Targets for Cancer Therapy: Why, Where, When, and How. Int J Mol Sci 21

49. Bowling, S., Di Gregorio, A., Sancho, M., Pozzi, S., Aarts, M., Signore, M., M, D. S., Martinez-Barbera, J. P., Gil, J., and Rodriguez, T. A. (2018) P53 and mTOR signalling determine fitness selection through cell competition during early mouse embryonic development. Nat Commun 9, 1763

50. White, P. J., McGarrah, R. W., Herman, M. A., Bain, J. R., Shah, S. H., and Newgard, C. B. (2021) Insulin action, type 2 diabetes, and branched-chain amino acids: A two-way street. Mol Metab 52, 101261

51. Wang, Q., Holmes, M. V., Davey Smith, G., and Ala-Korpela, M. (2017) Genetic Support for a Causal Role of Insulin Resistance on Circulating Branched-Chain Amino Acids and Inflammation. Diabetes Care 40, 1779–1786

52. Richardson, N. E., Konon, E. N., Schuster, H. S., Mitchell, A. T., Boyle, C., Rodgers, A. C., Finke, M., Haider, L. R., Yu, D., Flores, V., Pak, H. H., Ahmad, S., Ahmed, S., Radcliff, A., Wu, J., Williams, E. M., Abdi, L., Sherman, D. S., Hacker, T., and Lamming, D. W. (2021) Lifelong restriction of dietary branched-chain amino acids has sex-specific benefits for frailty and lifespan in mice. Nat Aging 1, 73–86

53. Neinast, M. D., Jang, C., Hui, S., Murashige, D. S., Chu, Q., Morscher, R. J., Li, X., Zhan, L., White, E., Anthony, T. G., Rabinowitz, J. D., and Arany, Z. (2018) Quantitative Analysis of the Whole-Body Metabolic Fate of Branched-Chain Amino Acids. Cell Metab

54. Crown, S. B., Marze, N., and Antoniewicz, M. R. (2015) Catabolism of Branched Chain Amino Acids Contributes Significantly to Synthesis of Odd-Chain and Even-Chain Fatty Acids in 3T3-L1 Adipocytes. PLoS One 10, e0145850

55. Yoneshiro, T., Wang, Q., Tajima, K., Matsushita, M., Maki, H., Igarashi, K., Dai, Z., White, P. J., McGarrah, R. W., Ilkayeva, O. R., Deleye, Y., Oguri, Y., Kuroda, M., Ikeda, K., Li, H., Ueno, A., Ohishi, M., Ishikawa, T., Kim, K., Chen, Y., Sponton, C. H., Pradhan, R. N., Majd, H., Greiner, V. J., Yoneshiro, M., Brown, Z., Chondronikola, M., Takahashi, H., Goto, T., Kawada, T., Sidossis, L., Szoka, F. C., McManus, M. T., Saito, M., Soga, T., and Kajimura, S. (2019) BCAA catabolism in brown fat controls energy homeostasis through SLC25A44. Nature 572, 614–619

56. Williams, R., Ma, X., Schott, R. K., Mohammad, N., Ho, C. Y., Li, C. F., Chang, B. S., Demetriou, M., and Dennis, J. W. (2014) Encoding asymmetry of the N-glycosylation motif facilitates glycoprotein evolution. PLoS One 9, e86088

57. Dennis, J. W. (2017) Genetic code asymmetry supports diversity through experimentation with posttranslational modifications. Curr Opin Chem Biol 41, 1–11

58. Yan, R., Zhao, X., Lei, J., and Zhou, Q. (2019) Structure of the human LAT1-4F2hc heteromeric amino acid transporter complex. Nature 568, 127–130

59. Gong, D., Chi, X., Ren, K., Huang, G., Zhou, G., Yan, N., Lei, J., and Zhou, Q. (2018) Structure of the human plasma membrane Ca(2+)-ATPase 1 in complex with its obligatory subunit neuroplastin. Nat Commun 9, 3623

60. Feral, C. C., Nishiya, N., Fenczik, C. A., Stuhlmann, H., Slepak, M., and Ginsberg, M. H. (2005) CD98hc (SLC3A2) mediates integrin signaling. Proc Natl Acad Sci U S A 102, 355–360

61. Palm, W., Park, Y., Wright, K., Pavlova, N. N., Tuveson, D. A., and Thompson, C. B. (2015) The Utilization of Extracellular Proteins as Nutrients Is Suppressed by mTORC1. Cell 162, 259–270

62. Nofal, M., Zhang, K., Han, S., and Rabinowitz, J. D. (2017) mTOR Inhibition Restores Amino Acid Balance in Cells Dependent on Catabolism of Extracellular Protein. Mol Cell 67, 936–946 e935

63. de la Ballina, L. R., Cano-Crespo, S., Gonzalez-Munoz, E., Bial, S., Estrach, S., Cailleteau, L., Tissot, F., Daniel, H., Zorzano, A., Ginsberg, M. H., Palacin, M., and Feral, C. C. (2016) Amino Acid Transport Associated to Cluster of Differentiation 98 Heavy Chain (CD98hc) Is at the Cross-road of Oxidative Stress and Amino Acid Availability. J Biol Chem 291, 9700–9711

64. Cano-Crespo, S., Chillaron, J., Junza, A., Fernandez-Miranda, G., Garcia, J., Polte, C., L, R. d. l. B., Ignatova, Z., Yanes, O., Zorzano, A., Stephan-Otto Attolini, C., and Palacin, M. (2019) CD98hc (SLC3A2) sustains amino acid and nucleotide availability for cell cycle progression. Sci Rep 9, 14065

65. Fan, J., Ye, J., Kamphorst, J. J., Shlomi, T., Thompson, C. B., and Rabinowitz, J. D. (2014) Quantitative flux analysis reveals folate-dependent NADPH production. Nature 510, 298–302

66. Ducker, G. S., and Rabinowitz, J. D. (2017) One-Carbon Metabolism in Health and Disease. Cell Metab 25, 27–42

67. Sullivan, L. B., Gui, D. Y., Hosios, A. M., Bush, L. N., Freinkman, E., and Vander Heiden, M. G. (2015) Supporting Aspartate Biosynthesis Is an Essential Function of Respiration in Proliferating Cells. Cell 162, 552–563

68. Zhang, C. S., Hawley, S. A., Zong, Y., Li, M., Wang, Z., Gray, A., Ma, T., Cui, J., Feng, J. W., Zhu, M., Wu, Y. Q., Li, T. Y., Ye, Z., Lin, S. Y., Yin, H., Piao, H. L., Hardie, D. G., and Lin, S. C. (2017) Fructose-1,6-bisphosphate and aldolase mediate glucose sensing by AMPK. Nature 548, 112–116

69. Vara-Ciruelos, D., Russell, F. M., and Hardie, D. G. (2019) The strange case of AMPK and cancer: Dr Jekyll or Mr Hyde? (dagger). Open Biol 9, 190099

70. Koppula, P., Zhang, Y., Shi, J., Li, W., and Gan, B. (2017) The glutamate/cystine antiporter SLC7A11/xCT enhances cancer cell dependency on glucose by exporting glutamate. J Biol Chem 292, 14240–14249

71. Ren, P., Yue, M., Xiao, D., Xiu, R., Gan, L., Liu, H., and Qing, G. (2015) ATF4 and N-Myc coordinate glutamine metabolism in MYCN-amplified neuroblastoma cells through ASCT2 activation. J Pathol 235, 90–100

72. Deprez, P., Gautschi, M., and Helenius, A. (2005) More than one glycan is needed for ER glucosidase II to allow entry of glycoproteins into the calnexin/calreticulin cycle. Molecular Cell 19, 183–195

73. Eyster, C. A., Cole, N. B., Petersen, S., Viswanathan, K., Fruh, K., and Donaldson, J. G. (2011) MARCH ubiquitin ligases alter the itinerary of clathrin-independent cargo from recycling to degradation. Mol Biol Cell 22, 3218–3230

74. Lau, K. S., Partridge, E. A., Grigorian, A., Silvescu, C. I., Reinhold, V. N., Demetriou, M., and Dennis, J. W. (2007) Complex N-glycan number and degree of branching cooperate to regulate cell proliferation and differentiation. Cell 129, 123–134

75. Maeda, K., Tasaki, M., Ando, Y., and Ohtsubo, K. (2020) Galectin-lattice sustains function of cationic amino acid transporter and insulin secretion of pancreatic beta cells. J Biochem 167, 587–596

76. Stegmayr, J., Zetterberg, F., Carlsson, M. C., Huang, X., Sharma, G., Kahl-Knutson, B., Schambye, H., Nilsson, U. J., Oredsson, S., and Leffler, H. (2019) Extracellular and intracellular small-molecule galectin-3 inhibitors. Sci Rep 9, 2186

77. Salameh, B. A., Cumpstey, I., Sundin, A., Leffler, H., and Nilsson, U. J. (2010) 1H-1,2,3-triazol-1-yl thiodigalactoside derivatives as high affinity galectin-3 inhibitors. Bioorg Med Chem 18, 5367–5378

78. Eyster, C. A., Higginson, J. D., Huebner, R., Porat-Shliom, N., Weigert, R., Wu, W. W., Shen, R. F., and Donaldson, J. G. (2009) Discovery of new cargo proteins that enter cells through clathrin-independent endocytosis. Traffic 10, 590–599

79. Schroeder, N., Chung, C. S., Chen, C. H., Liao, C. L., and Chang, W. (2012) The lipid raft-associated protein CD98 is required for vaccinia virus endocytosis. J Virol 86, 4868–4882

80. Shafaq-Zadah, M., Dransart, E., and Johannes, L. (2020) Clathrin-independent endocytosis, retrograde trafficking, and cell polarity. Curr Opin Cell Biol 65, 112–121

81. Go, C. D., Knight, J. D. R., Rajasekharan, A., Rathod, B., Hesketh, G. G., Abe, K. T., Youn, J. Y., Samavarchi-Tehrani, P., Zhang, H., Zhu, L. Y., Popiel, E., Lambert, J. P., Coyaud, E., Cheung, S. W. T., Rajendran, D., Wong, C. J., Antonicka, H., Pelletier, L., Palazzo, A. F., Shoubridge, E. A., Raught, B., and Gingras, A. C. (2021) A proximity-dependent biotinylation map of a human cell. Nature 595, 120–124

82. Youn, J. Y., Dyakov, B. J. A., Zhang, J., Knight, J. D. R., Vernon, R. M., Forman-Kay, J. D., and Gingras, A. C. (2019) Properties of Stress Granule and P-Body Proteomes. Mol Cell 76, 286–294

83. Forbes, K., Shah, V. K., Siddals, K., Gibson, J. M., Aplin, J. D., and Westwood, M. (2015) Statins inhibit insulin-like growth factor action in first trimester placenta by altering insulin-like growth factor 1 receptor glycosylation. Mol Hum Reprod 21, 105–114

84. Siddals, K. W., Marshman, E., Westwood, M., and Gibson, J. M. (2004) Abrogation of insulin-like growth factor-I (IGF-I) and insulin action by mevalonic acid depletion: synergy between protein prenylation and receptor glycosylation pathways. J Biol Chem 279, 38353–38359

85. Cavallini, G., Sgarbossa, A., Parentini, I., Bizzarri, R., Donati, A., Lenci, F., and Bergamini, E. (2016) Dolichol: A Component of the Cellular Antioxidant Machinery. Lipids 51, 477–486

86. Ishimoto, T., Nagano, O., Yae, T., Tamada, M., Motohara, T., Oshima, H., Oshima, M., Ikeda, T., Asaba, R., Yagi, H., Masuko, T., Shimizu, T., Ishikawa, T., Kai, K., Takahashi, E., Imamura, Y., Baba, Y., Ohmura, M., Suematsu, M., Baba, H., and Saya, H. (2011) CD44 variant regulates redox status in cancer cells by stabilizing the xCT subunit of system xc(-) and thereby promotes tumor growth. Cancer Cell 19, 387–400

87. Tamada, M., Nagano, O., Tateyama, S., Ohmura, M., Yae, T., Ishimoto, T., Sugihara, E., Onishi, N., Yamamoto, T., Yanagawa, H., Suematsu, M., and Saya, H. (2012) Modulation of glucose metabolism by CD44 contributes to antioxidant status and drug resistance in cancer cells. Cancer Res 72, 1438–1448

88. Sigismund, S., Lanzetti, L., Scita, G., and Di Fiore, P. P. (2021) Endocytosis in the context-dependent regulation of individual and collective cell properties. Nat Rev Mol Cell Biol 22, 625–643

89. Castiglione, G. M., Xu, Z., Zhou, L., and Duh, E. J. (2020) Adaptation of the master antioxidant response connects metabolism, lifespan and feather development pathways in birds. Nat Commun 11, 2476

90. Trautman, M. E., Richardson, N. E., and Lamming, D. W. (2022) Protein restriction and branched-chain amino acid restriction promote geroprotective shifts in metabolism. *Aging Cell*, e13626

91. Trefely, S., Lovell, C. D., Snyder, N. W., and Wellen, K. E. (2020) Compartmentalised acyl-CoA metabolism and roles in chromatin regulation. Mol Metab 38, 100941

92. McGarrah, R. W., Zhang, G. F., Christopher, B. A., Deleye, Y., Walejko, J. M., Page, S., Ilkayeva, O., White, P. J., and Newgard, C. B. (2020) Dietary branched-chain amino acid restriction alters fuel selection and reduces triglyceride stores in hearts of Zucker fatty rats. Am J Physiol Endocrinol Metab 318, E216–E223

93. Wagner, G. R., Bhatt, D. P., O’Connell, T. M., Thompson, J. W., Dubois, L. G., Backos, D. S., Yang, H., Mitchell, G. A., Ilkayeva, O. R., Stevens, R. D., Grimsrud, P. A., and Hirschey, M. D. (2017) A Class of Reactive Acyl-CoA Species Reveals the Non-enzymatic Origins of Protein Acylation. Cell Metab 25, 823–837 e828

94. Rezende, E. L., Bacigalupe, L. D., Nespolo, R. F., and Bozinovic, F. (2020) Shrinking dinosaurs and the evolution of endothermy in birds. Sci Adv 6, eaaw4486

95. Guglielmo, C. G. (2010) Move that fatty acid: fuel selection and transport in migratory birds and bats. Integr Comp Biol 50, 336–345

96. Szekely, P., Korem, Y., Moran, U., Mayo, A., and Alon, U. (2015) The Mass-Longevity Triangle: Pareto Optimality and the Geometry of Life-History Trait Space. PLoS Comput Biol 11, e1004524

97. Jarvis, E. D., Mirarab, S., Aberer, A. J., Li, B., Houde, P., Li, C., Ho, S. Y., Faircloth, B. C., Nabholz, B., Howard, J. T., Suh, A., Weber, C. C., da Fonseca, R. R., Li, J., Zhang, F., Li, H., Zhou, L., Narula, N., Liu, L., Ganapathy, G., Boussau, B., Bayzid, M. S., Zavidovych, V., Subramanian, S., Gabaldon, T., Capella-Gutierrez, S., Huerta-Cepas, J., Rekepalli, B., Munch, K., Schierup, M., Lindow, B., Warren, W. C., Ray, D., Green, R. E., Bruford, M. W., Zhan, X., Dixon, A., Li, S., Li, N., Huang, Y., Derryberry, E. P., Bertelsen, M. F., Sheldon, F. H., Brumfield, R. T., Mello, C. V., Lovell, P. V., Wirthlin, M., Schneider, M. P., Prosdocimi, F., Samaniego, J. A., Vargas Velazquez, A. M., Alfaro-Nunez, A., Campos, P. F., Petersen, B., Sicheritz-Ponten, T., Pas, A., Bailey, T., Scofield, P., Bunce, M., Lambert, D. M., Zhou, Q., Perelman, P., Driskell, A. C., Shapiro, B., Xiong, Z., Zeng, Y., Liu, S., Li, Z., Liu, B., Wu, K., Xiao, J., Yinqi, X., Zheng, Q., Zhang, Y., Yang, H., Wang, J., Smeds, L., Rheindt, F. E., Braun, M., Fjeldsa, J., Orlando, L., Barker, F. K., Jonsson, K. A., Johnson, W., Koepfli, K. P., O’Brien, S., Haussler, D., Ryder, O. A., Rahbek, C., Willerslev, E., Graves, G. R., Glenn, T. C., McCormack, J., Burt, D., Ellegren, H., Alstrom, P., Edwards, S. V., Stamatakis, A., Mindell, D. P., Cracraft, J., Braun, E. L., Warnow, T., Jun, W., Gilbert, M. T., and Zhang, G. (2014) Whole-genome analyses resolve early branches in the tree of life of modern birds. Science 346, 1320–1331

98. Satoh, T. (2021) Bird evolution by insulin resistance. Trends Endocrinol Metab 32, 803–813

99. Kapusta, A., Suh, A., and Feschotte, C. (2017) Dynamics of genome size evolution in birds and mammals. Proc Natl Acad Sci U S A 114, E1460–E1469

100. Lovell, P. V., Wirthlin, M., Wilhelm, L., Minx, P., Lazar, N. H., Carbone, L., Warren, W. C., and Mello, C. V. (2014) Conserved syntenic clusters of protein coding genes are missing in birds. Genome Biol 15, 565

101. Kim, E. B., Fang, X., Fushan, A. A., Huang, Z., Lobanov, A. V., Han, L., Marino, S. M., Sun, X., Turanov, A. A., Yang, P., Yim, S. H., Zhao, X., Kasaikina, M. V., Stoletzki, N., Peng, C., Polak, P., Xiong, Z., Kiezun, A., Zhu, Y., Chen, Y., Kryukov, G. V., Zhang, Q., Peshkin, L., Yang, L., Bronson, R. T., Buffenstein, R., Wang, B., Han, C., Li, Q., Chen, L., Zhao, W., Sunyaev, S. R., Park, T. J., Zhang, G., Wang, J., and Gladyshev, V. N. (2011) Genome sequencing reveals insights into physiology and longevity of the naked mole rat. Nature 479, 223–227

102. Ramsey, J. J., Harper, M. E., and Weindruch, R. (2000) Restriction of energy intake, energy expenditure, and aging. Free Radic Biol Med 29, 946–968

103. Swovick, K., Firsanov, D., Welle, K. A., Hryhorenko, J. R., Wise, J. P., Sr., George, C., Sformo, T. L., Seluanov, A., Gorbunova, V., and Ghaemmaghami, S. (2021) Interspecies Differences in Proteome Turnover Kinetics Are Correlated With Life Spans and Energetic Demands. Mol Cell Proteomics 20, 100041

104. She, P., Reid, T. M., Bronson, S. K., Vary, T. C., Hajnal, A., Lynch, C. J., and Hutson, S. M. (2007) Disruption of BCATm in mice leads to increased energy expenditure associated with the activation of a futile protein turnover cycle. Cell Metab 6, 181–194

105. She, P., Zhou, Y., Zhang, Z., Griffin, K., Gowda, K., and Lynch, C. J. (2010) Disruption of BCAA metabolism in mice impairs exercise metabolism and endurance. J Appl Physiol (1985) **108**, 941-949

106. Poncet, N., Mitchell, F. E., Ibrahim, A. F., McGuire, V. A., English, G., Arthur, J. S., Shi, Y. B., and Taylor, P. M. (2014) The catalytic subunit of the system L1 amino acid transporter (slc7a5) facilitates nutrient signalling in mouse skeletal muscle. PLoS One 9, e89547

107. Eggermann, T., Venghaus, A., and Zerres, K. (2012) Cystinuria: an inborn cause of urolithiasis. Orphanet J Rare Dis 7, 19

108. Nagamori, S., Wiriyasermkul, P., Guarch, M. E., Okuyama, H., Nakagomi, S., Tadagaki, K., Nishinaka, Y., Bodoy, S., Takafuji, K., Okuda, S., Kurokawa, J., Ohgaki, R., Nunes, V., Palacin, M., and Kanai, Y. (2016) Novel cystine transporter in renal proximal tubule identified as a missing partner of cystinuria-related plasma membrane protein rBAT/SLC3A1. Proc Natl Acad Sci U S A 113, 775–780

109. Sakamoto, Y., Taguchi, T., Honke, K., Korekane, H., Watanabe, H., Tano, Y., Dohmae, N., Takio, K., Horii, A., and Taniguchi, N. (2000) Molecular cloning and expression of cDNA encoding chicken UDP-N-acetyl-D-glucosamine (GlcNAc): GlcNAcbeta 1-6(GlcNAcbeta 1-2)-manalpha 1-R[GlcNAc to man]beta 1,4N-acetylglucosaminyltransferase VI. J Biol Chem 275, 36029–36034

110. Jouandin, P., Marelja, Z., Shih, Y. H., Parkhitko, A. A., Dambowsky, M., Asara, J. M., Nemazanyy, I., Dibble, C. C., Simons, M., and Perrimon, N. (2022) Lysosomal cystine mobilization shapes the response of TORC1 and tissue growth to fasting. Science 375, eabc4203

111. Ardalan, A., Smith, M. D., and Jelokhani-Niaraki, M. (2022) Uncoupling Proteins and Regulated Proton Leak in Mitochondria. Int J Mol Sci 23

112. Plassais, J., vonHoldt, B. M., Parker, H. G., Carmagnini, A., Dubos, N., Papa, I., Bevant, K., Derrien, T., Hennelly, L. M., Whitaker, D. T., Harris, A. C., Hogan, A. N., Huson, H. J., Zaibert, V. F., Linderholm, A., Haile, J., Fest, T., Habib, B., Sacks, B. N., Benecke, N., Outram, A. K., Sablin, M. V., Germonpre, M., Larson, G., Frantz, L., and Ostrander, E. A. (2022) Natural and human-driven selection of a single non-coding body size variant in ancient and modern canids. Curr Biol 32, 889–897 e889

113. Kraus, C., Pavard, S., and Promislow, D. E. (2013) The size-life span trade-off decomposed: why large dogs die young. Am Nat 181, 492–505

114. Paszek, M. J., DuFort, C. C., Rossier, O., Bainer, R., Mouw, J. K., Godula, K., Hudak, J. E., Lakins, J. N., Wijekoon, A. C., Cassereau, L., Rubashkin, M. G., Magbanua, M. J., Thorn, K. S., Davidson, M. W., Rugo, H. S., Park, J. W., Hammer, D. A., Giannone, G., Bertozzi, C. R., and Weaver, V. M. (2014) The cancer glycocalyx mechanically primes integrin-mediated growth and survival. Nature 511, 319–325

115. Shashar, M., Hod, T., Chernichovski, T., Angel, A., Kazan, S., Grupper, A., Naveh, S., Kliuk-Ben Bassat, O., Weinstein, T., and Schwartz, I. F. (2018) Mineralocorticoid receptor blockade improves arginine transport and nitric oxide generation through modulation of cationic amino acid transporter-1 in endothelial cells. Nitric Oxide 80, 24–31

116. Console, L., Scalise, M., Tarmakova, Z., Coe, I. R., and Indiveri, C. (2015) N-linked glycosylation of human SLC1A5 (ASCT2) transporter is critical for trafficking to membrane. Biochim Biophys Acta 1853, 1636–1645

117. Zandberg, W. F., Benjannet, S., Hamelin, J., Pinto, B. M., and Seidah, N. G. (2011) N-glycosylation controls trafficking, zymogen activation and substrate processing of proprotein convertases PC1/3 and subtilisin kexin isozyme-1. Glycobiology 21, 1290–1300

118. Cheng, C., Ru, P., Geng, F., Liu, J., Yoo, J. Y., Wu, X., Cheng, X., Euthine, V., Hu, P., Guo, J. Y., Lefai, E., Kaur, B., Nohturfft, A., Ma, J., Chakravarti, A., and Guo, D. (2015) Glucose-Mediated N-glycosylation of SCAP Is Essential for SREBP-1 Activation and Tumor Growth. Cancer Cell 28, 569–581

119. Guan, D., Wang, H., Li, V. E., Xu, Y., Yang, M., and Shen, Z. (2009) N-glycosylation of ATF6beta is essential for its proteolytic cleavage and transcriptional repressor function to ATF6alpha. J Cell Biochem 108, 825–831

120. Hong, M., Luo, S., Baumeister, P., Huang, J. M., Gogia, R. K., Li, M., and Lee, A. S. (2004) Underglycosylation of ATF6 as a novel sensing mechanism for activation of the unfolded protein response. J Biol Chem 279, 11354–11363

121. Conrad, M., Lorenz, S. M., and Proneth, B. (2021) Targeting Ferroptosis: New Hope for As-Yet-Incurable Diseases. Trends Mol Med 27, 113–122

122. Shi, Z., and Ruvkun, G. (2012) The mevalonate pathway regulates microRNA activity in Caenorhabditis elegans. Proc Natl Acad Sci U S A 109, 4568–4573

123. Rauthan, M., Ranji, P., Aguilera Pradenas, N., Pitot, C., and Pilon, M. (2013) The mitochondrial unfolded protein response activator ATFS-1 protects cells from inhibition of the mevalonate pathway. Proc Natl Acad Sci U S A 110, 5981–5986

124. Ohtsubo, K., Takamatsu, S., Minowa, M. T., Yoshida, A., Takeuchi, M., and Marth, J. D. (2005) Dietary and genetic control of glucose transporter 2 glycosylation promotes insulin secretion in suppressing diabetes. Cell 123, 1307–1321

125. Lee, Y., Berglund, E. D., Wang, M. Y., Fu, X., Yu, X., Charron, M. J., Burgess, S. C., and Unger, R. H. (2012) Metabolic manifestations of insulin deficiency do not occur without glucagon action. Proc Natl Acad Sci U S A 109, 14972–14976

126. Anthony-Regnitz, C. M., Wilson, A. E., Sweazea, K. L., and Braun, E. J. (2020) Fewer Exposed Lysine Residues May Explain Relative Resistance of Chicken Serum Albumin to In Vitro Protein Glycation in Comparison to Bovine Serum Albumin. J Mol Evol 88, 653–661

127. Chen, J. H., Lin, X., Bu, C., and Zhang, X. (2018) Role of advanced glycation end products in mobility and considerations in possible dietary and nutritional intervention strategies. Nutr Metab (Lond) 15, 72

128. Goodman, R. P., Markhard, A. L., Shah, H., Sharma, R., Skinner, O. S., Clish, C. B., Deik, A., Patgiri, A., Hsu, Y. H., Masia, R., Noh, H. L., Suk, S., Goldberger, O., Hirschhorn, J. N., Yellen, G., Kim, J. K., and Mootha, V. K. (2020) Hepatic NADH reductive stress underlies common variation in metabolic traits. Nature 583, 122–126

129. Robertson, H., Dinkova-Kostova, A. T., and Hayes, J. D. (2020) NRF2 and the Ambiguous Consequences of Its Activation during Initiation and the Subsequent Stages of Tumourigenesis. Cancers (Basel) 12

130. Bonini, M. G., Consolaro, M. E. L., Hart, P. C., Mao, M., de Abreu, A. L. P., and Master, A. M. (2014) Redox control of enzymatic functions: The electronics of life’s circuitry. IUBMB Life 66, 167–181

131. Chen, P. H., Smith, T. J., Wu, J., Siesser, P. F., Bisnett, B. J., Khan, F., Hogue, M., Soderblom, E., Tang, F., Marks, J. R., Major, M. B., Swarts, B. M., Boyce, M., and Chi, J. T. (2017) Glycosylation of KEAP1 links nutrient sensing to redox stress signaling. EMBO J 36, 2233–2250

132. Lehrbach, N. J., Breen, P. C., and Ruvkun, G. (2019) Protein Sequence Editing of SKN-1A/Nrf1 by Peptide:N-Glycanase Controls Proteasome Gene Expression. Cell 177, 737–750 e715

133. Huang, C., Harada, Y., Hosomi, A., Masahara-Negishi, Y., Seino, J., Fujihira, H., Funakoshi, Y., Suzuki, T., Dohmae, N., and Suzuki, T. (2015) Endo-beta-N-acetylglucosaminidase forms N-GlcNAc protein aggregates during ER-associated degradation in Ngly1-defective cells. Proc Natl Acad Sci U S A 112, 1398–1403

134. Zhang, Y., and Manning, B. D. (2015) mTORC1 signaling activates NRF1 to increase cellular proteasome levels. Cell Cycle 14, 2011–2017

135. Liu, Y., Samuel, B. S., Breen, P. C., and Ruvkun, G. (2014) Caenorhabditis elegans pathways that surveil and defend mitochondria. Nature 508, 406–410

136. Grenier-Larouche, T., Coulter Kwee, L., Deleye, Y., Leon-Mimila, P., Walejko, J. M., McGarrah, R. W., Marceau, S., Trahan, S., Racine, C., Carpentier, A. C., Lusis, A. J., Ilkayeva, O., Vohl, M. C., Huertas-Vazquez, A., Tchernof, A., Shah, S. H., Newgard, C. B., and White, P. J. (2022) Altered branched-chain alpha-keto acid metabolism is a feature of NAFLD in individuals with severe obesity. JCI Insight 7

137. Buffenstein, R. (2008) Negligible senescence in the longest living rodent, the naked mole-rat: insights from a successfully aging species. J Comp Physiol B 178, 439–445

138. Csiszar, A., Labinskyy, N., Orosz, Z., Xiangmin, Z., Buffenstein, R., and Ungvari, Z. (2007) Vascular aging in the longest-living rodent, the naked mole rat. Am J Physiol Heart Circ Physiol 293, H919–927

139. Tian, X., Azpurua, J., Hine, C., Vaidya, A., Myakishev-Rempel, M., Ablaeva, J., Mao, Z., Nevo, E., Gorbunova, V., and Seluanov, A. (2013) High-molecular-mass hyaluronan mediates the cancer resistance of the naked mole rat. Nature 499, 346–349

140. Azpurua, J., Ke, Z., Chen, I. X., Zhang, Q., Ermolenko, D. N., Zhang, Z. D., Gorbunova, V., and Seluanov, A. (2013) Naked mole-rat has increased translational fidelity compared with the mouse, as well as a unique 28S ribosomal RNA cleavage. Proc Natl Acad Sci U S A 110, 17350–17355

141. Yu, C., Li, Y., Holmes, A., Szafranski, K., Faulkes, C. G., Coen, C. W., Buffenstein, R., Platzer, M., de Magalhaes, J. P., and Church, G. M. (2011) RNA sequencing reveals differential expression of mitochondrial and oxidation reduction genes in the long-lived naked mole-rat when compared to mice. PLoS One 6, e26729

142. Maldonado-Baez, L., Cole, N. B., Kramer, H., and Donaldson, J. G. (2013) Microtubule-dependent endosomal sorting of clathrin-independent cargo by Hook1. J Cell Biol 201, 233–247

143. Shin, Y., and Brangwynne, C. P. (2017) Liquid phase condensation in cell physiology and disease. Science 357

144. Wang, T., Medynets, M., Johnson, K. R., Doucet-O’Hare, T. T., DiSanza, B., Li, W., Xu, Y., Bagnell, A., Tyagi, R., Sampson, K., Malik, N., Steiner, J., Hadegan, A., Kowalak, J., O’Malley, J., Maric, D., and Nath, A. (2020) Regulation of stem cell function and neuronal differentiation by HERV-K via mTOR pathway. Proc Natl Acad Sci U S A 117, 17842-17853

145. Prud’homme, B., Gompel, N., and Carroll, S. B. (2007) Emerging principles of regulatory evolution. Proceedings of the National Academy of Sciences of the United States of America 104 **Suppl 1**, 8605–8612

146. Helbing, D. (2013) Globally networked risks and how to respond. Nature 497, 51–59

147. Ran, F. A., Hsu, P. D., Wright, J., Agarwala, V., Scott, D. A., and Zhang, F. (2013) Genome engineering using the CRISPR-Cas9 system. Nat Protoc 8, 2281–2308

148. Livak, K. J., and Schmittgen, T. D. (2001) Analysis of relative gene expression data using real-time quantitative PCR and the 2(-Delta Delta C(T)) Method. Methods 25, 402–408

149. Abdel Rahman, A. M., Ryczko, M., Pawling, J., and Dennis, J. W. (2013) Probing the hexosamine biosynthetic pathway in human tumor cells by multitargeted tandem mass spectrometry. ACS Chem Biol 8, 2053–2062

150. Xia, J., and Wishart, D. S. (2011) Web-based inference of biological patterns, functions and pathways from metabolomic data using MetaboAnalyst. Nat Protoc 6, 743–760

151. Hesketh, G. G., Youn, J. Y., Samavarchi-Tehrani, P., Raught, B., and Gingras, A. C. (2017) Parallel Exploration of Interaction Space by BioID and Affinity Purification Coupled to Mass Spectrometry. Methods Mol Biol 1550, 115–136

152. Lambert, J. P., Picaud, S., Fujisawa, T., Hou, H., Savitsky, P., Uuskula-Reimand, L., Gupta, G. D., Abdouni, H., Lin, Z. Y., Tucholska, M., Knight, J. D. R., Gonzalez-Badillo, B., St-Denis, N., Newman, J. A., Stucki, M., Pelletier, L., Bandeira, N., Wilson, M. D., Filippakopoulos, P., and Gingras, A. C. (2019) Interactome Rewiring Following Pharmacological Targeting of BET Bromodomains. Mol Cell 73, 621–638 e617

153. Liu, G., Knight, J. D., Zhang, J. P., Tsou, C. C., Wang, J., Lambert, J. P., Larsen, B., Tyers, M., Raught, B., Bandeira, N., Nesvizhskii, A. I., Choi, H., and Gingras, A. C. (2016) Data Independent Acquisition analysis in ProHits 4.0. J Proteomics 149, 64–68

154. Jamwal, R., Barlock, B. J., Adusumalli, S., Ogasawara, K., Simons, B. L., and Akhlaghi, F. (2017) Multiplex and Label-Free Relative Quantification Approach for Studying Protein Abundance of Drug Metabolizing Enzymes in Human Liver Microsomes Using SWATH-MS. J Proteome Res 16, 4134–4143

